# Dog-human gene network colocalization links synaptic signalling and ubiquitination to compulsive disorders

**DOI:** 10.64898/2026.10.12.758413

**Authors:** Katarina Tengvall, Vista Sohrab, Matthew J. Christmas, Gustaf Brander, Eric Pederson, Raphaela Pensch, Brittney Kenney, Chengcheng Song, Ola Wallerman, Elisabeth Sundström, Chao Wang, Åsa Karlsson, Linnéa L. Monsén, Nordic OCD and Related Disorders Consortium (NORDiC), PGC TS/OCD working group, Kathryn A. Lord, Nora I. Strom, Christian Rück, David Mataix-Cols, Manuel Mattheisen, James J. Crowley, Jennifer R. S. Meadows, Elinor K. Karlsson, Kerstin Lindblad-Toh

## Abstract

Obsessive-compulsive disorder (OCD) is highly polygenic, and the largest genome-wide association study (GWAS), with 53,660 cases and 2,044,417 controls, identified only 30 loci, a small fraction of loci estimated to explain its heritability – too few to resolve biological pathways^1^. Pet dogs develop a compulsive disorder (dogCD) similar to OCD^2^. In a GWAS of dogCD with 2,584 dogs from the community science project Darwin’s Ark, each dog locus explains at least 40-fold more phenotypic variance than each human OCD locus, giving power equivalent to the human GWAS with 80-fold fewer individuals. Combining the two GWAS identifies a 165-gene network undetectable in humans at current sample sizes. This network is enriched for genes expressed in neurons, particularly in the amygdala and cortex, and comprises four biological processes: excitatory-inhibitory synaptic transmission, axon guidance and neuron recognition, protein ubiquitination, and adenylyl cyclase-activating G-protein-coupled receptor signaling. Like most complex human traits^3^, dogCD risk is likely driven by regulatory variation: fine-mapped dogCD SNPs are non-coding, and dogCD loci are enriched in candidate cis-regulatory elements (cCREs) in a new dog epigenomic atlas spanning eight brain regions. The cross-species network implicates an inherited risk architecture largely outside current OCD pharmacotherapy, and establishes pet dogs as a model for gene network discovery, particularly for conditions underpowered in human studies.

## Introduction

Obsessive-compulsive disorder (OCD) is a psychiatric disorder affecting 1–3% of the global population with profound personal and societal costs^4–6^, and new treatment options are needed. Current first-line treatments are selective serotonin reuptake inhibitors (SSRIs), which are ineffective in up to 60% of patients^7^, and cognitive-behavioural therapy (CBT), to which a substantial proportion of patients show no clinically meaningful response^8^. However, efforts to develop new treatments have been hampered by an incomplete understanding of OCD’s underlying neurobiology^9^.

Dysfunction in the cortico-striatal-thalamo-cortical (CSTC) circuits has repeatedly been implicated in OCD pathophysiology^10^. Converging evidence indicates abnormalities across multiple interconnected networks, with altered glutamatergic excitation and GABAergic inhibition thought to contribute to impaired behavioural control and compulsive symptoms^11,12^. Consistent with this model, glutamate-modulating agents show promise in treatment-refractory cases^13,14^. However, the specific molecular networks mediating this excitatory-inhibitory imbalance remain poorly characterised, limiting the development of more precise therapeutic targets^9^.

Genome-wide association studies (GWAS) offer an opportunity to define the underlying networks for psychiatric diseases through unbiased discovery of risk-conferring variants, but require very large sample sizes^15^. For example, for schizophrenia, GWAS (76,755 cases and 243,649 controls) identified 287 genome-wide significant loci, and 120 genes, that functionally converge on synaptic biology in excitatory and inhibitory neurons^16^. This includes the gene *DRD2*, the target of every current antipsychotic, providing genetic validation for an existing drug class. The other genes constitute a large set of novel candidates for therapeutic development, as genetic support for a drug target more than doubles its probability of being approved^17^.

OCD GWAS have not yet reached a scale sufficient for robust inference of the underlying molecular networks. The largest OCD GWAS to date (53,660 cases and 2,044,417 controls, including samples from the 23andMe Research Institute) identified 30 genome-wide significant loci. Consistent with previous work implicating the CSTC circuit^1^, OCD genetic risk is enriched in genes expressed in excitatory neurons of the cortex and hippocampus, and two classes of striatal neurons (D1 and D2 medium spiny neurons). However, the 30 loci are a tiny fraction of the 11,500 variants estimated to underlie 90% of OCD’s heritability. This is higher than for schizophrenia (∼9,600), although lower than for major depression (∼14,500)^1^.

To close this gap expediently, strategies beyond simply scaling up the size of the human cohorts are needed. Case recruitment for OCD presents challenges that slow the growth of adequately powered cohorts. OCD is frequently unrecognised clinically, with a mean 8-year delay from symptom onset to treatment^18^, limiting the pool of diagnosed individuals, and population biobanks underrepresent those most severely affected, due to the healthy volunteer effect^19^. Compensating by pooling cases across cohorts that differ in how OCD was ascertained introduces phenotypic heterogeneity that deflates SNP heritability estimates^1^. The same challenges arise with other compulsive and internalising disorders, precluding cross-disorder aggregation as a straightforward solution^1,19^.

The evolutionary conservation of compulsive behaviour across mammals suggests a complementary strategy: using naturally occurring compulsive disorders in other mammals to power the search for the gene networks underlying OCD. Gene regulatory pathways are broadly shared across placental mammals^20,21^, including those active in the brain between dogs and humans^22^. Because of this shared molecular architecture, if statistical frameworks for quantifying gene network colocalization show that the same network is implicated in a given trait across two species, this is evidence for a functional role^23^. For example, combining human and rat GWAS for body mass index using NetColoc revealed convergent synaptic, epigenetic, and hormonal signalling networks that were not apparent from gene-level comparison of either species alone^24^.

The compulsive component of human OCD has strong parallels to canine compulsive disorder (dogCD)^2,25^, presenting an opportunity for network colocalization. In both species, the disorder is characterized by excessive repetition of normal behaviour to the point of causing distress or functional impairment^25–27^. The specific behaviours are species-typical, with dogs displaying flank sucking, tail chasing, and false digging (**Fig. 1a**), rather than checking, ordering, or washing. Onset in dogs occurs during early social maturity (6–12 months), comparable to early-onset human OCD^26^, and dogs are treated with the same combination of SSRIs and behavioural modification used in humans, with comparable SSRI response rates as reported in small dog trials^28,29^. Whether dogs experience the obsessive component of OCD cannot be determined in a non-verbal species; however, compulsions can occur without clearly articulated obsessions in humans too, particularly in early-onset cases^30^. Structural neuroimaging in Doberman pinschers with dogCD has revealed abnormalities in the dorsal anterior cingulate cortex, anterior insula, and corpus callosum that parallel findings in human OCD, implicating the same CSTC circuitry disrupted in the human disorder^31^.

**Fig. 1.**
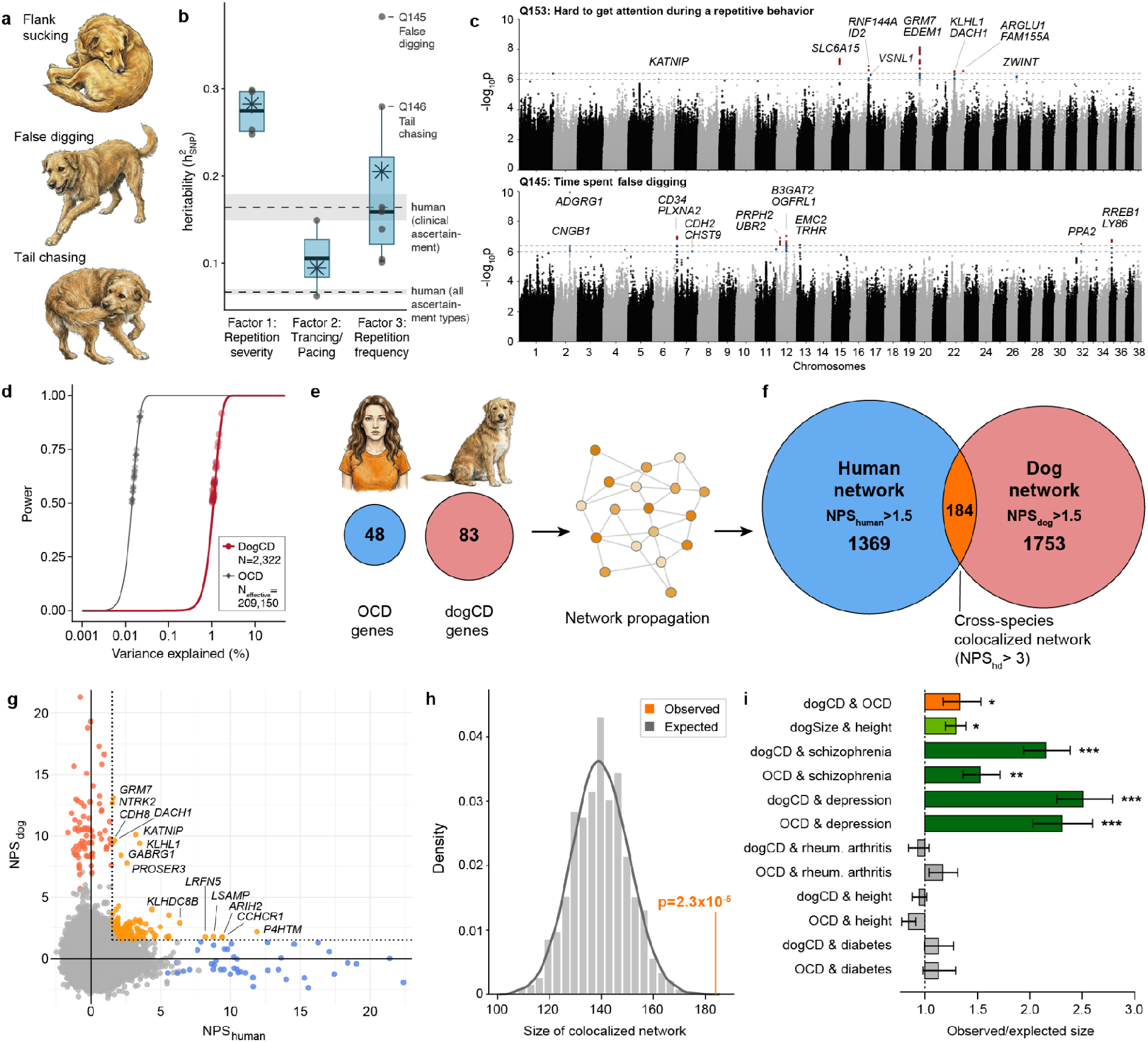
Convergence of human OCD and dogCD genes in shared network. (**a**) Three dog compulsive behaviors. (**b**) LD-corrected heritability of factors (asterisks; unconstrained for factor 2) and underlying survey items (filled grey circles; top 2 Factor 3 items labeled, Q150 not estimable), with median (thick blue line), interquartile range (blue box), and 1.5 × IQR (whiskers). Dashed lines and grey bands show human OCD h²SNP ± s.e.^1^. (**c**) Manhattan plots for two dogCD items. (**d**) Power to detect a locus in GWAS for dogCD (p ≤ 4x10^-7^) and OCD (p ≤ 5x10^-8^). Points are significant loci. (**e**) OCD and dogCD seed gene sets used in network propagation are distinct. **(f)** Genes exceeding species-specific and combined (orange) NPS thresholds after network propagation (NPS_hd_ = NPS_human_ × NPS_dog_). NPS thresholds defined by sensitivity analysis (Methods): p = 2.1x10^-5^. (**g**) NPS_human_ vs NPS_dog_ for all PCNet2.0 genes (N = 19,267), including colocalized network genes (orange; seed genes labeled), human-only (blue), dog-only (red), and other genes (grey). (**h**) Observed (orange, n = 184) versus expected (grey distribution; mean = 138.98, s.d. = 10.98) size of the colocalized gene network (10,000 permutations). (**i**) Observed/expected colocalized network size for dogCD & OCD (orange) and other trait pairs with (green) and without (grey) genetic overlap. *p≤2.3x10^-5^, **p≤3.2x10^-14^, ***p≤1.4x10^-68^. Error bars show 95% confidence intervals.

Previous genetic studies in dogCD reinforce the parallels between dogs and humans. Four dogCD risk genes have been identified in dogs, all with synaptic functions: *PGCP* and *CDH2*, involved in glutamatergic signalling and synapse formation, and *CTNNA2* and *ATXN1*^32,33^. Cross-species analysis integrating dog, mouse, and human data implicated synaptic adhesion in compulsive disorders^34^, and rare missense variants in *CDH2* — first identified as a dogCD risk gene — are linked to co-occurring OCD and Tourette syndrome in humans^35^. Notably, the most recent human OCD GWAS independently identified *CTNND1* as a top associated gene^1^, a component of the same cadherin-catenin adhesion complex as *CTNNA2*.

Dogs offer practical advantages for GWAS of complex traits, including behavioural disorders. Population bottlenecks and selection have altered the allele frequency spectrum, reducing the sample sizes needed^36–38^, and the close relationship between dogs and humans enables phenotyping through owner surveys and mail-in sample collection^25^. Darwin’s Ark, a community science project, collects genetic information from owner-collected saliva swabs alongside owner-reported phenotypes on thousands of pet dogs, including the dogCD phenotypes used in this study^39^. A previous GWAS drawing on this resource, with 2,155 dogs of diverse breed ancestry, identified loci associated with a complex behaviour, howling propensity, near genes implicated in oral-motor development in humans^38^.

Here, we integrate human OCD GWAS from the largest study to date with dogCD GWAS data from 2,584 pet dogs recruited through Darwin’s Ark to identify molecular networks underlying compulsive behaviour that data from neither species could resolve alone.

## Results

### Three dimensions of dog compulsive behavior

Through the community science project Darwin’s Ark, we assembled a cohort of dogs with dogCD phenotypes based on owner surveys. Darwin’s Ark invites any dog owner to enroll their pet, and its participants are representative of the breed diversity of dogs in the US, with approximately half being mixed breed^38^. After enrolling, owners are asked to complete a series of short behavioral surveys which includes a survey of 11 questions developed by board-certified veterinary behaviorists to assess the frequency, severity, and type of compulsive behaviors characteristic of dogCD^2,40^ and three related questions from other surveys (**Supplementary Table 1**)^41,42^. In total, owners of 25,302 pet dogs in Darwin’s Ark had answered at least one dogCD-related question, and 12,989 had answered all 14.

Through exploratory factor analysis^43^, we resolved the survey responses into three latent dimensions: Factor 1 captures the severity and interruptibility of repetitive behaviors (repetition severity); Factor 2 captures unresponsive or trance-like behaviors, including blank staring and aimless pacing (trancing/pacing); and Factor 3 captures the frequency and duration of specific repetitive motor behaviors such as licking, tail chasing, and light chasing (repetition frequency)^44^. We used item response theory to derive continuous factor scores for each dog from their survey responses.

### DogCD’s genetic architecture enables high-power locus discovery

For 2,584 of the dogCD-phenotyped dogs, we also generated high-density genotypes from owner-collected saliva swabs. Most dogs were selected for genotyping based on enrollment date (i.e., coming off the waitlist as funding became available). We also included 315 dogs based on the dogCD survey questions to ensure dogs with higher symptom severity were represented. The earliest samples were genotyped using a SNP array with 1,011,992 SNPs (Axiom Canine Array A+B; N = 373)^45^, and we subsequently shifted to low-pass sequencing (mean depth = 0.83, s.d. = 0.35; N = 2,211). We imputed genotypes for both data types against the Dog10K reference panel^46^, yielding variant calls for up to 9,885,182 SNPs with MAF ≥ 1% (**Supplementary Table 2**). We confirmed that imputed genotypes are 98.6% concordant with genotypes from deep sequencing of 11 dogs (**Supplementary Table 3**).

DogCD traits are heritable, with SNP-based heritability estimates for nearly all traits exceeding those reported for human OCD^1^. Using restricted maximum likelihood with LD-score correction, common SNPs explained 28% ± 8% of variation in repetition severity (Factor 1) and 21% ± 7% in repetition frequency (Factor 3). For trancing/pacing (Factor 2), the LD-constrained model failed to converge; the unconstrained, LD-corrected estimate was 9% ± 5%. All four survey items underlying repetition severity (hard to get attention, hard to interrupt, interferes with daily life, and triggered by excitement) had heritabilities exceeding 25%, as did time spent false digging (38% ± 8%) and time spent tail chasing (28% ± 9%) (**Supplementary Table 4**). Every dogCD trait, except Q93 “paces, circles, or wanders aimlessly” (6% ± 5%) and Q150 “time spent in trance-like behavior” (LD-constrained model failed to converge), exceeded the pooled SNP-heritability reported across all human OCD ascertainment types (h²SNP = 6.7% ± 0.3%)¹, and half of the dogCD traits exceeded those reported for the more narrowly ascertained clinical (h²SNP = 16.4% ± 1.5%) subgroup from the same study^1^ (**Fig. 1b**).

Dozens of genomic regions are significantly associated with at least one of the 14 dogCD survey questions or three factors. We tested for association using two mixed-model approaches that account for population structure: MLMA-LOCO^47^ for the continuous factors and POLMM^48^ for the ordinal survey items. The full summary statistics for all 14 survey questions and three factors are available through an interactive browser^49^ built using a version of PheWeb adapted for the dog genome^50,51^. We then fine-mapped signals into credible-set intervals with SuSiE^52^ and assigned overlapping genes, using the nearest gene within ±1 Mb when an interval contained none. At the dog-specific threshold^53^ of p ≤ 4×10^-7^, 27 regions (46 genes) are genome-wide significant, and 28 additional regions (41 genes) are suggestive (p ≤ 1×10^-6^) (**Fig. 1c**; **Supplementary Table 5**; **Extended Data Fig. 1**). One interval, on chromosome 1, contributed 10 genes, and the others all contributed two or fewer. In total 83 genes with 1:1 human orthologues^54,55^ are in regions with at least a suggestive association to a dogCD trait. None of these candidate dogCD genes are amongst the 48 genes implicated by at least two gene-based mapping approaches in the largest human OCD GWAS (“candidate human OCD genes”)^1^ **(Supplementary Table 6**).

Each dogCD locus explains far more phenotypic variance than each human OCD locus, giving the dogCD GWAS sufficient power at much smaller sample sizes than the human OCD GWAS. DogCD’s significant loci individually explain 1.0-1.7% of phenotypic variance, compared with 0.014-0.023% for human OCD’s significant loci. Thus, dogCD achieves power equivalent to the human OCD GWAS with only 2,322–2,555 dogs, more than 80-fold smaller than the human GWAS’s effective sample size of 209,150 (**Fig. 1d**). This is consistent with population bottlenecks, and potentially selection, driving large-effect risk alleles to higher frequency in dogs. The effect sizes are likely inflated by the winner’s curse phenomenon and require replication^56^, but the disparity in per-locus effect sizes between dogCD and human OCD is one to two orders of magnitude larger than expected even for weaker associations near threshold of significance^57^. Questions with imbalanced responses carry less effective information, as illustrated by Q147 “time spent catching invisible flies” and Q152 “time spent chasing lights or reflections”, where over 90% of dogs fall in a single category; consistent with this, neither item had significant associations (**Supplementary Table 1**).

### Colocalized network links dog and human psychiatric disorders

Network colocalization shows that dogCD and human OCD converge on shared gene networks despite implicating distinct genes. We propagated seed genes from each species through PCNet2.0^58,59^, a composite network of ∼3.85 million pairwise associations among 19,267 human genes (18,725 with 1:1 dog orthologues^55^). As seeds we used all candidate genes present in PCNet2.0: 82 of 83 dogCD genes and all 48 human OCD genes, none of which overlap (**Fig. 1e)**. We then colocalized the two networks with NetColoc^23^, which tests whether two gene sets cluster within overlapping network neighborhoods. We retained genes with species-specific network proximity scores (NPS_dog_ and NPS_human_) > 1.5 and a combined score (NPS_hd_ = NPS_dog_ × NPS_human_) > 3.0. These thresholds were chosen by sensitivity analysis (**Supplementary Table 7**). The resulting cross-species network (184 genes) is larger than expected by chance (p = 2.3x10^-5^; z-test against 10,000 permutations of NPS; **Fig. 1f-h**).

To confirm that dogCD-OCD colocalization reflects trait-specific biology rather than a generic artifact of network propagation, we tested network colocalization in pairs of traits that included both positive (traits with shared genetics) and negative controls (traits lacking shared genetics; **Supplementary Tables 8-10**; **Fig 1i**).

As a positive control, we tested whether network colocalization detects known shared biology for a trait unrelated to OCD: height. A network built from a human height GWAS^60^ colocalizes significantly with a network built from a dog size GWAS in the Darwin’s Ark population^39^, consistent with the well-established conservation of body-size pathways across mammals^61^.

As a further positive control, we tested whether the dogCD network colocalizes with networks for two human psychiatric disorders known to share substantial genetic architecture with OCD: depression and schizophrenia (r_g_=0.36-0.60 with OCD^1^). If colocalization between dogCD and OCD reflects shared biology, we should also see colocalization with other traits that share the same genetic risk factors as OCD. We confirmed significant colocalization for both. In fact, the signal is stronger than observed for OCD itself (depression/dogCD: p = 2.2x10^-122^; schizophrenia/dogCD: p = 1.5x10^-73^), likely reflecting the larger gene sets available for these two disorders.

As negative controls, we tested whether the dogCD network colocalizes with gene networks for traits with no known shared genetic architecture with OCD: rheumatoid arthritis^62^, type 2 diabetes^63^, or height^60^ (r_g_ < 0.05)^1^. The dogCD network does not significantly colocalize with GWAS-derived networks for any of these traits. The lack of colocalization is not because the three traits are less influenced by genetics, as all three traits have heritabilities comparable to OCD (SNP heritabilities h² of ∼0.18^64,65^, up to ∼0.40 for height^66^).

As a final negative control, we confirmed that the human height network that colocalizes with the dog-size network does not colocalize with the dogCD network, showing that cross-species colocalization is not a general property of a high scoring network.

In total, we build seven distinct psychiatric disorder-related networks using NetColoc^23^ and HiDeF^67^: single-species networks for each disorder, and cross-species networks colocalizing dogCD with each human disorder. Each network constitutes the top-level community in the hierarchy of nested communities of interacting genes resolved by HiDeF. Each cross-species network is smaller than the human-only network, and retains fewer seed genes (10–28%, compared with all seeds in the single-species networks; **Fig. 2a**). This reflects different thresholds for inclusion. Genes in cross-species networks are required to exceed the proximity thresholds in both species independently and combined. The OCD and dogCD networks are smaller than those for depression and schizophrenia, possibly reflecting their smaller seed sets.

**Fig. 2.**
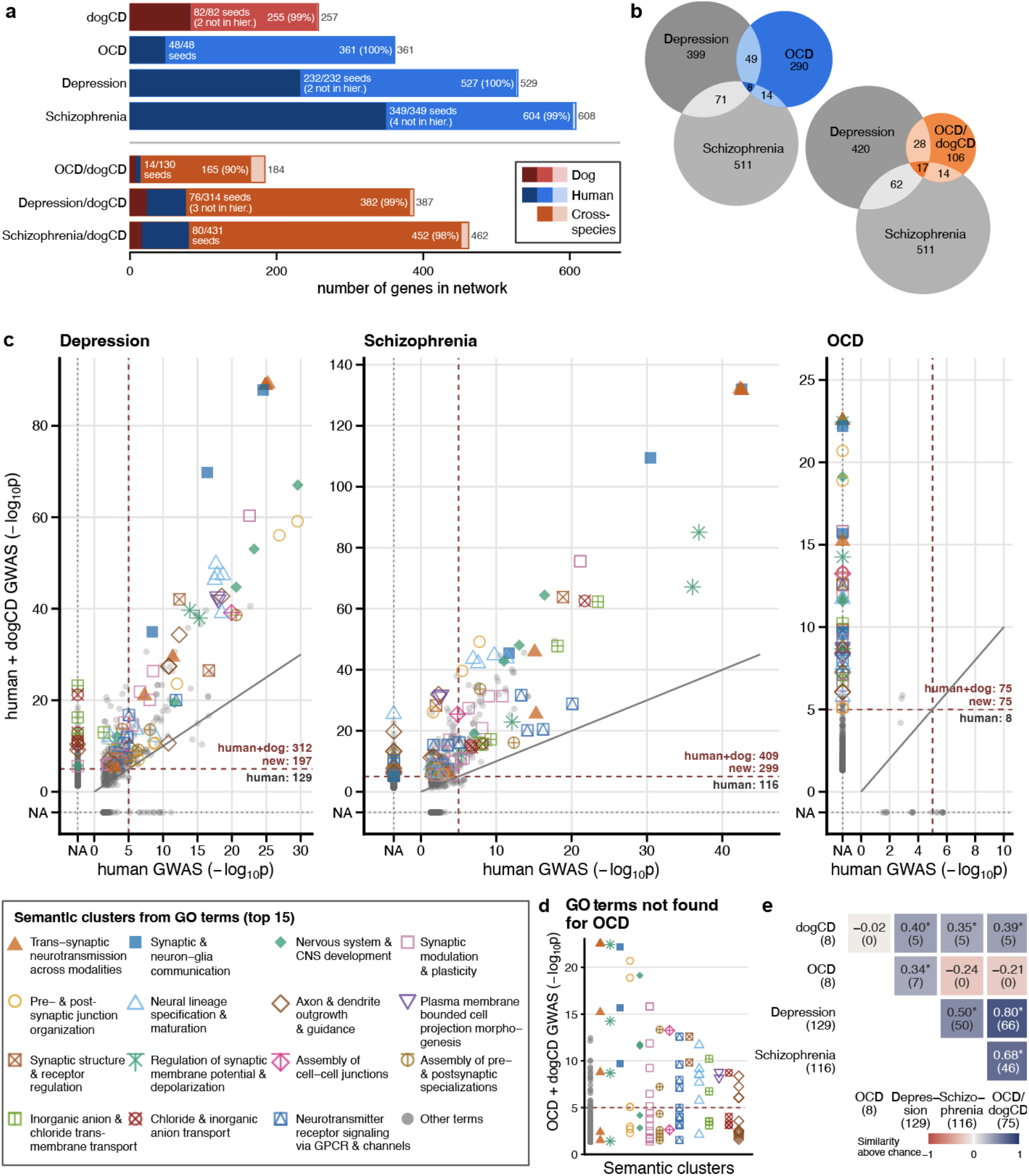
Human-dog colocalized networks of dogCD and three psychiatric disorders. (**a**) Size of each network in the top level hierarchy. Bar length gives the total number of genes per network. Bar colour indicates species, with shade indicating seed genes (dark), other network genes (intermediate) or not in hierarchy (light). Gene counts and excluded seeds noted alongside. (**b**) Area-proportional Venn diagrams of hierarchy-assigned genes shared between the human-only OCD, depression, and schizophrenia networks, and between the cross-species OCD/dogCD, depression, and schizophrenia networks. (**c**) GO term enrichment (−log_10_p) in the cross-species network versus the human-only network for depression, schizophrenia and OCD. Terms are grouped into the top 15 semantically related clusters by GOSemSim^70^ (symbols; remaining clusters grey). The grey dotted line marked ‘NA’ indicates terms without any genes, the red dashed line is the significance threshold (p=1x10^-5^). Numbers are the counts of significant terms in each network, and ‘new’ is the count of terms significant in only the cross-species network. (**d**) GO terms significant in the OCD-dogCD network and with no genes in the OCD network. (**e**) Pairwise semantic similarity between significantly enriched GO Biological Process terms for each network pair, scored relative to chance (Methods). Axis labels give each network’s total significant terms; numbers in parentheses are shared terms. *BH-corrected empirical p < 0.05.

### Cross-species OCD/dogCD network overlaps other psychiatric disorders

For OCD, cross-species colocalization produces a smaller network (165 vs 361 genes) with more overlap with the depression and schizophrenia networks compared to the single-species network (17 vs 8 genes shared with both; **Fig. 2b**; **Supplementary Table 8**). The composition of the shared genes also changes. The 8 genes shared between the OCD, depression and schizophrenia networks are functionally heterogeneous, spanning complement regulation, protein palmitoylation, cell adhesion and transcription. In contrast, the distinct set of 17 genes shared between the OCD/dogCD, depression and schizophrenia networks mostly encode synaptic signalling components, in line with previous mechanistic studies implicating glutamate and GABA receptor genes in compulsive disorders^11,12^. These include six ionotropic and metabotropic glutamate receptors, five GABA-A receptor subunits, the RNA splicing regulator RBFOX1, the presynaptic active-zone scaffold BSN, the adhesion and guidance molecules CDH8, OPCML and SEMA6D and the sodium channel SCN2A.

Cross-species colocalization produces gene networks with greater functional coherence, as measured by gene set enrichment, compared to the human-only networks. We tested each network for enrichment in GO Biological Process terms^68,69^. Semantically related terms were then clustered into higher-level annotations using GOSemSim^70^ (**Fig. 2c,d**; **Supplementary Table 11**). Despite having fewer genes than their human-only counterparts, the cross-species networks find more significant GO terms (terms with p<1x10^-5^) for all three disorders: 2.4-fold for depression (129 to 312), 3.5-fold for schizophrenia (116 to 409), and 9.4-fold for OCD (8 to 75). The enrichments are also stronger: of the terms significant in both networks, most are more significant in the cross-species network (108 of 129 for depression, 110 of 116 for schizophrenia), with median -log_10_p rising from 9.7 to 17.5 and from 7.8 to 25.4, respectively. None of the 8 terms significant in the human-only OCD network are among the 75 significant in the OCD/dogCD network.

The gene set enrichments for the OCD/dogCD network are also semantically more similar to those for depression and schizophrenia than those for the human-only OCD network, suggesting the cross-species network captures more psychiatric-disorder-relevant biology than the human-only network. We use depression and schizophrenia as reference points because their larger seed gene sets yield more robust networks compared to OCD. Because OCD shares genetic risk with both disorders^71^, we expect the underlying pathways to partially overlap. The single-species OCD network overlaps with depression (7 shared terms), but not with schizophrenia (0 shared terms), with which it has less, but still substantial, genetic correlation (r_g_=0.60±SE0.02 and 0.36±SE0.02 respectively)^1^. In contrast, the OCD/dogCD network overlaps both (66 terms with depression, 46 with schizophrenia, and 37 with both), and the GO terms enriched in the OCD/dogCD network are more semantically similar overall to the depression and schizophrenia networks than to either the human-only OCD network or the dog-only dogCD network (**Fig. 2e**). There is no evidence for compulsivity-specific biology, with all GO terms more enriched in both the depression and schizophrenia networks than in the OCD/dogCD network.

### Compulsive behavior communities have synaptic, ubiquitin, and aGPCR themes

Decomposing the cross-species OCD/dogCD network into a hierarchy of subcommunities reveals distinct functional themes that are not apparent from enrichment on the whole network (**Fig. 3a**). We annotated each subcommunity in the hierarchical networks produced by HiDeF by testing for enrichment in GO Biological Process ontology terms^68,69^ and Mammalian Phenotype (MP) Ontology terms^72^, which capture the phenotypic consequences of genetic manipulations in mice (**Fig. 3b**; **Supplementary Table 11**).

**Fig. 3.**
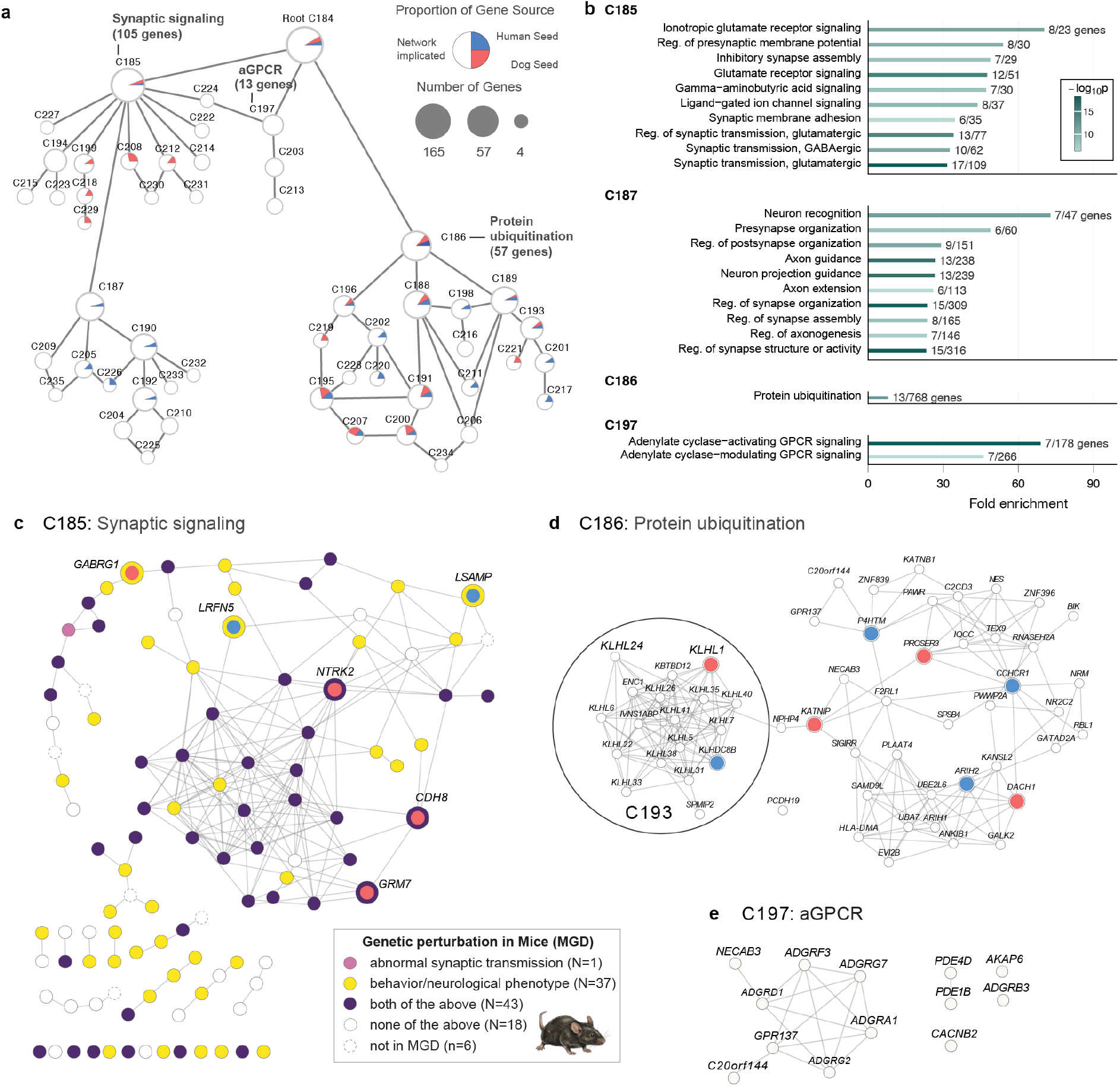
Hierarchical gene communities within the OCD/dogCD network. (**a**) The network consists of 52 gene nested communities (nodes) with between 4 and 165 genes each. Each node is a pie chart showing the fraction of genes in the community that are human seeds (blue), dog seeds (red), or network genes (white). Three higher-level gene communities were annotated based on the enrichment of GO Biological Process terms. (**b**) GO-term fold enrichment for the three major modules from (a), plus subcommunity C187 within the synaptic-signalling module (C185). (**c**) C185 community is enriched for genes, including seed genes from dog (inset red circle) and human (inset blue circle), in two mammalian phenotype (MP) terms. MGD: Mouse Genome Database. (**d**) The community C186 includes four seed genes from dog (red) and two from human (blue), and the subcommunity C193 (circled) is dominated by Kelch-like proteins and ubiquitin enzymes. (**e**) The community C197 includes genes involved in aGPCR. Edges shown in c-e have cosine similarities above 0.95.

#### Synaptic signalling community C185

The largest community (C185; 104 genes; **Fig. 3a-b**) includes both neurotransmitter receptors and the adhesion molecules that organize the synapses containing them. It is enriched for GO terms describing both excitatory (glutamatergic) and inhibitory (GABAergic) receptor signalling. It is also enriched for terms related to synaptic membrane adhesion, driven by a mostly distinct set of genes (only 1 out of 6 overlap). Within C185, HiDeF resolves a nested subcommunity (C187; 43 genes; **Fig. 3a-b**) enriched for axon guidance and neuron projection terms, driven by a nearly distinct set of genes from the receptor- and adhesion-signalling genes enriched in its parent community, C185 (only 4 out of 29 overlap) (**Extended Data Fig. 2)**.

Testing C185 for enrichment in MP terms also discovers a strong synaptic signalling theme, and implicates specific behavioral phenotypes related to OCD. Of the 141 MP terms enriched across the OCD/dogCD network and its HiDeF-defined subcommunities, 133 are associated with C185. The top term is ‘abnormal synaptic transmission’ (44/1239 genes, OR = 10.9, p = 2.8x10^-26^), and C185 is broadly enriched for genes associated with ‘behavior/neurological phenotype’ in mice (80/5760 genes, OR = 7.60, p = 1.7x10^-23^) (**Fig. 3c**). The enriched terms span phenotypic domains relevant to compulsive disorders: synaptic physiology, brain morphology, locomotor behaviour, learning and memory, and anxiety^73^ (**Supplementary Table 12**). They include ‘abnormal grooming behavior’ (8/170 genes, OR = 9.7, p = 3.8x10^-7^) and ‘abnormal anxiety-related response’ (26/818 genes, OR = 7.6, p = 2.2x10^-14^) – both phenotypes described in mouse models for OCD^74,75^. Only 9 of the 133 terms are non-neural phenotypes like abnormal body size and postnatal lethality.

#### Protein ubiquitination community C186

The second community (C186; 57 genes; **Fig. 3b,d**) is enriched for genes involved in protein ubiquitination, the process by which ubiquitin tags proteins for degradation. Dysregulation of this system in the brain contributes to neurodegeneration and neurodevelopmental disorders^76–78^. C186 contains a subcommunity (C193) with five Kelch-like (KLHL) substrate adaptors for the CUL3-RING E3 ubiquitin ligase complex, many of which are expressed in the nervous system^79^, as well as IVNS1ABP (*KLHL39*), a sixth KLHL family member that modulates CUL3 activity indirectly. A seventh KLHL family member, *KLHL1*, is the highest-scoring gene in the OCD/dogCD network (NPS_hd_ = 33.0) and is significantly associated with repetition severity (Factor 1) in the dogCD GWAS.

#### G-protein coupled receptor community C197

Community C197 (13 genes; **Fig. 3e**) implicates adhesion GPCRs (aGPCRs), a family with established roles in neurodevelopment, synapse modulation, and myelination^80^, in compulsive behaviour. Six of the thirteen community genes are aGPCRs, including *ADGRB3*, a critical regulator of synapse development implicated in neurodevelopmental phenotypes and disrupted by perinatal SSRI exposure^80,81^, and *ADGRA1*, linked to anxiety-like behaviour and disrupted hippocampal inhibitory circuit function in mice^82^.

The top GO enrichment term for C197, adenylate cyclase-activating G protein-coupled receptor signalling (**Fig. 3b**), connects the community to experimental models of compulsive behaviour. Five aGPCRs driving this term alongside proteins PDE4D and AKAP6, which degrade cAMP and scaffold PKA (the principal cAMP effector) in neurons^83^. Elevated cortico-striatal cAMP, resulting from reduced PDE4 activity, is linked to stereotypic behaviour in an animal model, with attenuation by fluoxetine, a drug used in dogs and humans to treat compulsive disorders^28^. C197 also includes *CACNB2*, a voltage-gated calcium channel subunit and PKA target associated with multiple psychiatric disorders^84^. Outside the colocalized network, the gene *ADCY2*, which encodes adenylate cyclase 2 and is a validated bipolar disorder risk gene^85^, shows suggestive dogCD association (Q150 “time spent in trance-like behaviour”; p =7x10^-7^), further implicating cAMP signalling in compulsive behaviour.

### GWAS loci support network community themes

Two dogCD GWAS loci independently support the C197 aGPCR/cAMP community, and neither has been previously implicated in psychiatric disease through GWAS in humans. The strongest overall (Q145 “time spent false digging”; p = 1.1x10^-10^) maps to a single gene, *ADGRG1*, an adhesion GPCR like those found in C197, which is required for cortical development in oligodendrocytes^86^, and its loss disrupts frontal brain regions that show altered connectivity in OCD^87^. A second signal is at *ADRA2C* (Q146 “time spent tail chasing”; p = 3.9x10^-8^), which encodes an adrenergic receptor that reduces cAMP by inhibiting adenylyl cyclase, connecting it to the cAMP signalling theme of C197. Mice lacking *ADRA2C* show increased startle, reduced prepulse inhibition, and impaired working memory^88^.

Both dog and human GWAS genes support the C193 ubiquitination community. In dogs, KLHL1, a KLHL-family ubiquitin adapter, has the second-strongest dogCD GWAS signal overall. This two-gene locus (*KLHL1* and *DACH1*) is significantly associated with three dogCD traits: two individual survey items, Q154 “hard to interrupt during a repetitive behavior” (p = 2.2x10^-9^) and Q153 “hard to get attention during a repetitive behavior” (p = 2.7x10^-7^), and one composite factor, repetition severity (CCDF1; p = 1.1x10^-7^). *KLHL1* and the human OCD GWAS gene *KLHDC8B* encode proteins from the same kelch-repeat family; both seeded the network (**Fig. 1f**), and HiDeF propagated them into the kelch-repeat adaptor dominated community C193 (**Fig. 3d**). In human GWAS, *KLHDC8B* is associated with a number of psychiatric traits, including depression^89^, anxiety^90^ and insomnia^91^, the last of which is the only one to also implicate *KLHL1* (**Supplementary Table 6**). *DACH1* has no reported psychiatric disease associations in human GWAS.

One single-gene dogCD GWAS region (Q93 “paces, circles, or wanders aimlessly”; p = 3.8×10^-7^) implicates *PCDH10*, a striatal protocadherin that drives elimination of excitatory synapses by targeting postsynaptic scaffolding proteins for proteasomal degradation. *PCDH10* has no reported psychiatric disease associations in human GWAS. However, the human OCD GWAS gene *MEF2C* transcriptionally activates *PCDH10* in response to neuronal activity^92^, placing them in the same activity-dependent synaptic pruning pathway. *PCDH10* is implicated in autism spectrum disorder and OCD through rare variant studies^93^.

Overall, 12 of 27 dogCD GWAS regions contain at least one gene associated with a psychiatric trait in the GWAS catalog, including *SLC6A15* (Q154 “hard to interrupt during a repetitive behavior”; p = 4.95×10^-9^), which is a replicated depression GWAS gene^89,94^; *GRM7* (Q153 “hard to get attention during a repetitive behavior”; p = 7.44×10^-9^), a metabotropic glutamate receptor associated with depression^95^; *NTRK2* (Q146 “time spent tail chasing”; p = 3.5×10^-7^), a gene associated with anxiety^96^ and depression^94^; *CDH8* (Q150 “time spent in trance-like behaviour”; p = 3.5×10^-7^), a gene associated with ADHD and multiple psychiatric disorders^97–99^; and *PBX1* (Q149 “time spent object/social licking”; p = 2.5×10^-7^), a gene associated with tics and stuttering^100^.

### Functional genomics links OCD/dogCD genes to specific brain circuits

The OCD/dogCD network genes are preferentially expressed in neurons of cortico-striatal and amygdalo-cortical circuitry implicated in compulsive behavior^101^. In both human^102^ and dog brain single-nuclei atlases^103^, the 165 network genes show significantly greater expression in neurons over glia (human: median relative expression 0.24 vs. 0.08; dog: 0.18 vs. 0.08; Wilcoxon rank-sum test, p < 2.2×10^-16^ in both). Cross-species correlations are highest in matching neuronal cell types (Spearman r = 0.80-0.89; **Extended Data Fig. 3**), consistent with conserved cell-type-specific transcriptional programs across placental mammals^104^. In the human atlas, which sampled twice as many brain regions as the dog atlas, expression specificity is enriched in the amygdala and cerebral cortex (permutation test, p = 0.002 and p = 0.03; **Fig. 4a**). Expression is most specific to MGE-derived interneurons, eccentric medium spiny neurons, and amygdala excitatory neurons (**Fig. 4b**).

**Fig. 4.**
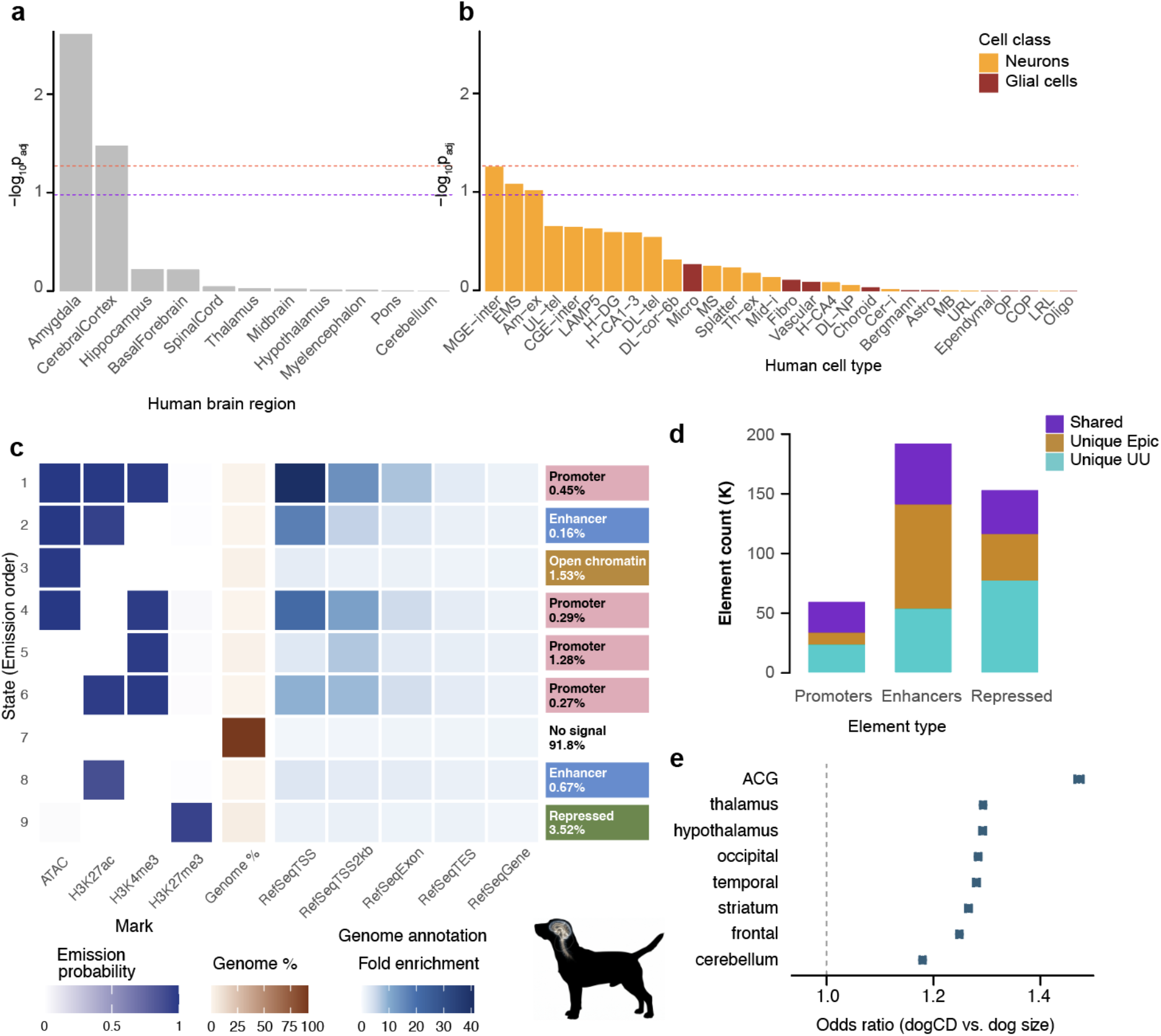
Brain relevance: enrichment of network genes in human single-cell data and of dogCD regions in a new dog epigenomic catalogue. **a-b.** Enrichment of OCD/dogCD network genes in the human brain single-nuclei atlas^102^ in (**a**) brain regions and (**b**) cell types (p<0.05, red dashed line; p<0.10, purple dashed line). **(c)** A new epigenomic catalogue of regulatory elements across eight regions of the adult dog brain. Candidate cis-regulatory elements (cCREs) are defined by nine chromatin states, each supported by a distinct combination of histone marks and ATAC signal, characterized as promoters, enhancers, open chromatin or repressed elements (state type and % genome in that state labeled to right). (**d**) Comparison between our elements (’UU’) and those from the EpiC Dog resource (cerebellum and cerebrum). The UU dataset identifies more promoters, EpiC Dog more enhancers. (**e**) Odds ratios for dogCD versus dog size GWAS region overlap with cCREs across dog brain regions in the UU dataset.

DogCD GWAS regions are enriched in brain regulatory elements, unlike GWAS regions for a non-behavioral trait (dog size) that we used as a comparison. Using the Epigenome Catalog of the Dog (EpiC Dog)^105^, we compared the cCRE basepair overlap of regions associated with dogCD to the regions from the dog size GWAS. DogCD regions are enriched in brain cCREs compared to other tissues (0.33% vs. 0.31% overlap; OR = 1.06, 95% CI 1.058-1.064, Fisher’s exact test, p < 2.2×10^-16^), while dog size regions showed the opposite — significant depletion of brain cCREs relative to other tissues (0.26% vs. 0.31%; OR = 0.84, 95% CI 0.842-0.847, p < 2.2×10⁻^16^; **Supplementary Table 13**). This double dissociation confirms that the brain enrichment observed for dogCD is tissue-specific rather than a generic property of intersecting any well-powered GWAS with a large regulatory annotation.

To annotate functional elements in dogCD GWAS regions that are active in specific brain regions and cell types, we developed a new epigenomic catalogue of regulatory elements of the adult dog brain identifying candidate cis-regulatory elements (cCREs)^106^. The existing dog brain resource, EpiC Dog^105^, covers a broader range of tissues, enabling brain versus non-brain comparisons, but includes only cerebellum and cerebrum as brain regions. We complement this by profiling eight brain regions (anterior cingulate gyrus (ACG), cerebellum, frontal cortex, hypothalamus, occipital cortex, striatum, temporal cortex, and thalamus) using ATAC-seq and histone ChIP-seq for H3K4me3, H3K27ac, and H3K27me3. We annotated promoter-like elements, active enhancers, open chromatin, and repressed states using ChromHMM^107^(**Fig. 4c**). Our catalogue identifies more candidate promoters and repressed states than EpiC Dog, but fewer enhancers, likely reflecting the absence of H3K4me1 profiling (**Fig. 4d**).

Regions associated with dogCD overlap brain cCREs more than regions associated with dog size in every one of the eight profiled brain regions (raw overlap ranging from 5.7% vs. 3.9% in anterior cingulate gyrus (ACG) to 10.7% vs. 8.5% in thalamus; Fisher’s exact test, FDR-corrected p < 2.2x10^-16^ in all regions; **Fig. 4e**; **Supplementary Table 14**). Although thalamus has the highest raw overlap, the relative enrichment (odds ratio for dogCD vs. dog size) is highest in ACG (OR=1.47, 95% CI 1.46-1.48) and lowest in cerebellum (OR = 1.18, 95% CI 1.17-1.18). ACG also has the smallest cCRE annotation of the eight regions (109.8 Mb, versus 147.9-213.8 Mb elsewhere), suggesting its outsized enrichment should be interpreted cautiously. However, annotation size does not explain most of the regional variation in enrichment. Thalamus has the largest annotation (213.8 Mb) yet one of the highest odds ratios (1.29), comparable to much smaller regions. Annotation size and odds ratio are not significantly correlated across the eight regions (Spearman’s ρ = -0.37, p = 0.36), and even less so when ACG is excluded (ρ = -0.05, p = 0.91).

#### Fine-mapped dogCD SNPs in cCREs

Fine-mapping of the 27 genome-wide significant loci using SuSiE identified 148 candidate causal SNPs, none coding, consistent with genetic risk operating through context-specific gene regulation (**Extended Data Fig. 4**; **Supplementary Table 15**). Fourteen SNPs across 7 loci overlap a brain-active cCRE. At *ADGRG1* (Q145 “time spent false digging”; p = 1.1×10^-10^), a single-variant credible set overlaps hypothalamic and striatal enhancers; credible variants at *NTRK2* (Q146 “time spent tail chasing”; p = 3.5×10^-7^), a member of the C185 synaptic network, overlap hypothalamic and thalamic enhancers. Extending to suggestive loci, credible variants at *KATNIP* (Q153 “hard to get attention during a repetitive behavior”; p = 9.4x10^-7^) and *UTP20* (Q155 “repetitive behaviors interfere with daily life”; p = 4.5x10^-7^) fall in promoter elements active across all eight profiled brain regions.

### Network communities implicate convergent neural mechanisms

Glutamatergic synaptic signalling emerges as a molecular mechanism of compulsive disorder conserved across species, driven by the largest community (C185, **Fig. 5a,b**), which is enriched for all three classes of ionotropic glutamate receptors as well as presynaptic and perisynaptic metabotropic glutamate receptors^108,109^. This is consistent with previous work associating glutamatergic signalling genes *PGCP* and *CDH2* with dogCD^32,33^, as well as genetic linkage^110^, gene expression^111^, and neurochemical studies in people with OCD^112^, including magnetic resonance spectroscopy (MRS) findings of elevated glutamate in certain brain regions, particularly in paediatric cases^113^.

**Fig. 5.**
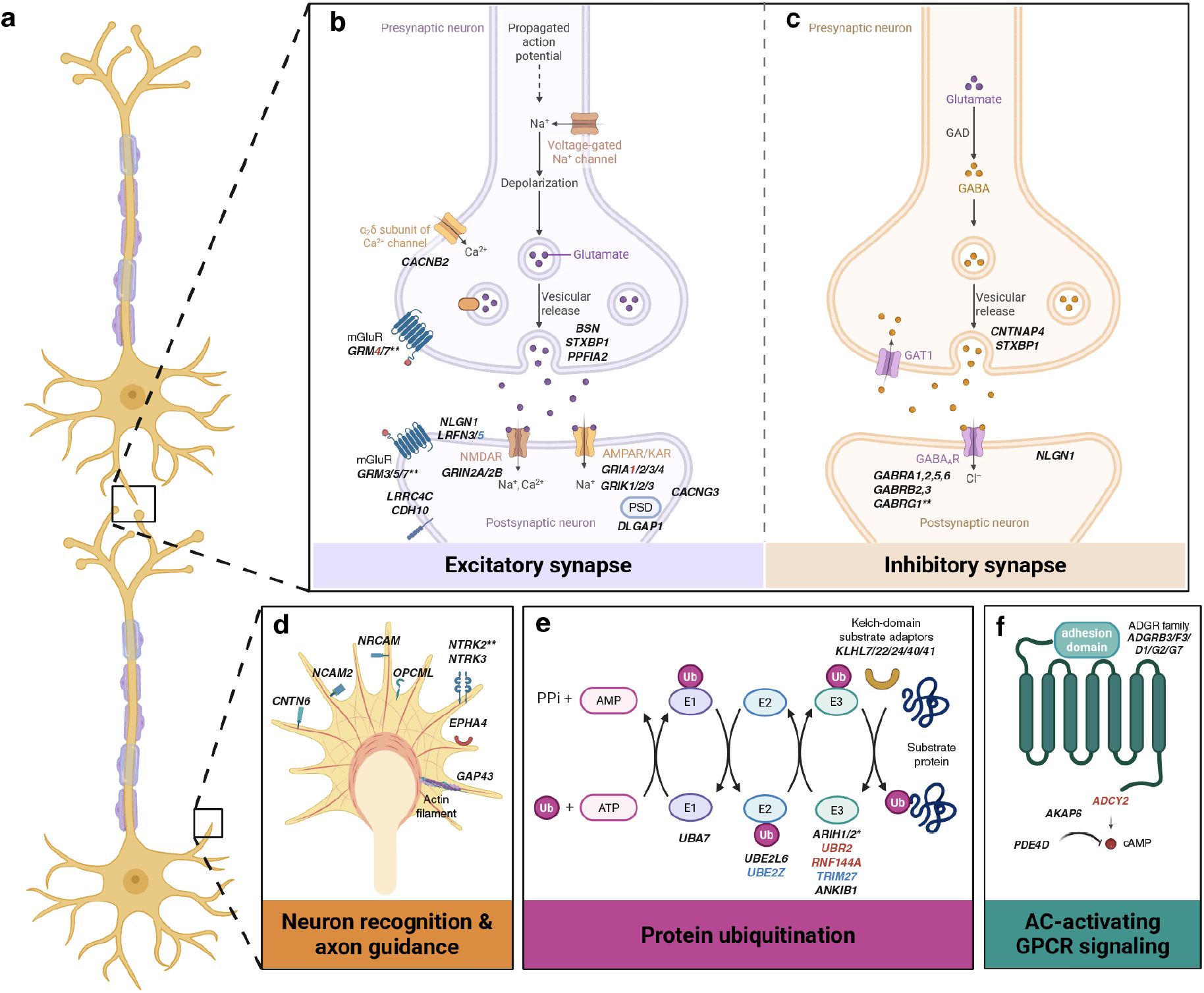
Molecular mechanisms implicated by network communities C185, C187, C186 and C197. (**a**) Neuron schematics highlighting the synaptic (top; expanded in b, c) and growth cone (bottom; expanded in d). (**b**) Excitatory and (**c**) inhibitory synapses, with C185 genes positioned by synaptic compartment (presynaptic release machinery and receptors, postsynaptic receptors, scaffolding and adhesion molecules). (**d**) Growth cone schematic showing C187 genes involved in neuron recognition (GO:0008038) and axon guidance (GO:0007411). (**e**) Protein ubiquitination cascade (C186, GO:0016567), from E1/E2 enzymes through E3 ligases to Kelch-domain substrate adaptors. (**f**) AC-activating GPCR signaling module (C197, GO:0007189), from adhesion GPCRs through cAMP production and degradation. Gene labels are italicized. Black labels are network genes, with asterisks marking those also present as dog (**) or human (*) seed genes. Red and blue labels are seed genes shown for context that are not part of the network (red, dog; blue, human). Created in BioRender, Tengvall, K. (2026), https://BioRender.com/76pmxqm.

The largest community also implicates disrupted excitatory-inhibitory (E/I) balance as the underlying mechanism. Inhibitory GABA-A receptor subunits are as prominent in C185 as excitatory glutamate receptors (**Fig. 5b,c**), consistent with MRS studies reporting an altered glutamate-to-GABA ratio in OCD^11,114^. C185 implicates intrinsic neuronal excitability as well: it includes voltage-gated channel genes, among them the schizophrenia-associated genes *KCNB1* and *CACNB2*, where variants are predicted to alter neuronal spiking and calcium dynamics^115,116^. Many of the genes involved in E/I imbalance are shared across the OCD/dogCD, depression and schizophrenia networks: out of the 17 genes in all three networks, six are ionotropic or metabotropic glutamate receptors and five are GABA-A receptor subunits, providing cross-species genetic support for excitatory-inhibitory imbalance at the cortical synapse as a transdiagnostic feature of psychiatric risk^117^.

Within the largest community, HiDeF resolves a nested subcommunity (C187, **Fig. 5d**) that points to disrupted neurodevelopmental wiring as a contributor to compulsive behaviour. It implicates axon guidance and neuron recognition, both developmental processes that establish corticostriatal circuits by directing growing axons to their targets. The resulting axon bundles, visible as white matter tracts in brain scans, show abnormalities in OCD patients; crucially, these changes are also present in unaffected relatives, suggesting they affect early brain development rather than being consequences of the disorder itself^118^.

Synaptic protein turnover emerges as a complementary layer of control over E/I balance, driven by a smaller community (C186, **Fig. 5e**). This community converges on KLHL-family substrate-recognition adaptors for Cullin-RING E3 ligase complexes, which tag synaptic proteins for proteasomal degradation and thereby control receptor surface expression and synaptic strength^119^. This provides a mechanistic link to C185, the largest community: ubiquitination-mediated trafficking of AMPA and GABA-A receptors is a primary means of maintaining E/I balance at the synapse. C186 is supported by GWAS genes from both species (three from human, three from dog), but its coherence as a ubiquitination programme is apparent only through cross-species colocalization, not from either species’ data alone. KLHL proteins are not yet established drug targets, but their substrate specificity makes them attractive candidates for targeted protein degradation approaches currently under development^120,121^.

Finally, a signalling pathway gains genetic support from one of the smallest communities (C197, **Fig. 5f**). The 12 genes in C197 converge on AC-activating GPCR signalling, with cAMP scaffolding via AKAP6 and degradation by PDE4D. This pathway has direct experimental links to stereotyped behaviour in rodent models^122^, and its independent emergence here in the OCD/dogCD cross-species network supports its relevance to compulsive behaviour.

## Discussion

OCD in humans is polygenic, yet most animal models reduce this complexity to near single-gene disruptions^74,75,123^ or circuit^124^ manipulations. Compulsive disorders in pet dogs, however, like human OCD^2,33^, are complex, environmentally influenced and, crucially, polygenic, allowing us to define networks through GWAS. Using a dog cohort more than 80-fold smaller than the human OCD GWAS’s effective sample size, we exploit this shared genetic architecture through cross-species network colocalization. Each dogCD locus explains 43- to 121-fold more phenotypic variance than each human OCD locus, giving sufficient statistical power despite the smaller sample size. This approach reveals gene communities not apparent from the human OCD GWAS alone, mapping onto four biological processes: excitatory–inhibitory synaptic transmission, axon guidance and neuron projection, protein ubiquitination, and AC-activating GPCR signalling (**Fig. 5**).

In humans, psychiatric disorders are not genetically distinct entities, but instead are largely explained by a small number of shared genomic factors^71,84,125^. The dogCD network overlaps both the depression and schizophrenia networks more than expected by chance, suggesting the dogCD network captures biology relevant to human psychiatric disease. The overlap of dogCD with human OCD is lower, likely because OCD’s GWAS to date has yielded fewer significant genes, so the resulting OCD network captures less of the underlying disease biology. When we colocalize the OCD and dogCD networks, however, the resulting cross-species network has greater overlap with both depression and schizophrenia than either single-species network alone. The cross-species networks between dogCD and all three human diseases are also more functionally coherent: despite having fewer genes than their human-only counterparts, they show more significant GO term enrichments. Together, these results suggest that cross-species colocalization can recover genes and pathways relevant to compulsive behaviour not detected in the human-only OCD GWAS by leveraging convergent biology across species.

The OCD/dogCD network implicates an inherited risk architecture for compulsive disorders that lies largely outside established OCD pharmacotherapy, which pairs SSRIs with antipsychotic augmentation targeting dopamine and serotonin receptors^18^. This points toward new therapeutic targets that act on specific glutamate receptor subtypes rather than broadly modulating glutamate signalling. The clinical evidence already supports this distinction: drugs that broadly modulate glutamate release and clearance have failed in large trials^126,127^, but agents acting on specific receptors are more promising. A network meta-analysis ranked the NMDA-receptor antagonist memantine as the most effective augmentation of SSRIs at moderate certainty^128^, and a single infusion of ketamine, another NMDA-receptor antagonist, can rapidly reduce obsessions^129^. Although the cross-species network contains no serotonin-receptor genes, it is consistent with the efficacy of 5-HT3 antagonists as augmentation^130,131^: 5-HT3 receptors are expressed predominantly on GABAergic interneurons, where their antagonism directly modulates cortical E/I balance through the same ligand-gated ion channel signalling implicated by the network^132,133^.

The convergence we observe between OCD and dogCD across gene community structure, cell-type expression, and brain regulatory enrichment provides molecular-level validation of the dog as a model for compulsive disorder. The model captures the compulsive dimension of OCD; whether it extends to the obsessional component (the intrusive cognitions that also define the human disorder) remains untested. That this convergence is already discernible with fewer than 2,600 dogs demonstrates substantial power at a modest sample size. The dog model is augmented by our new dog brain cCRE atlas, which can help finemap high-confidence causal variants^134^ and connect dog GWAS signals to mechanisms. Expanding it to additional brain regions, such as the amygdala^135,136^, would make it more powerful, and is feasible through partnerships with existing post-mortem tissue donation programmes at veterinary hospitals, paralleling the design of human resources such as PsychENCODE. Dogs and other household pets develop complex, polygenic behavioural disorders in the same environments we do^25,137^. Large-scale GWAS is feasible in these populations: owner participation is demonstrably not the bottleneck, with Darwin’s Ark alone already enrolling 50,329 dogs and 25,019 cats, both of which develop compulsive disorders. As these resources grow, integrating pet and human genetics will help untangle the regulatory genomics of behaviour.

## Nordic OCD and Related Disorders Consortium (NORDiC)

Bjarne Hansen^1^, Christian Rück^2^, David Mataix-Cols^2^, Diana Pascal^2^, Elinor K. Karlsson^3^, Elles de Schipper^2^, Fredrik Johansson^2^, Gerd Kvale^1^, James J. Crowley^4^, Jan Haavik^5^, John Wallert^2^, Julia Bäckman^2^, Kerstin Lindblad-Toh^6^, Kira D. Höffler^7^, Kristen Hagen^8^, Long Long Chen^2^, Manuel Mattheisen^9^, Matthew W. Halvorsen^4^, Nora I. Strom^10^, Paul Lichtenstein^11^, Thorstein Olsen Eide^8^, Josep Pol Fuster^2^, Katarina Tengvall^6^, Stephanie Le Hellard^12^, Gustaf Brander^2^ & Jennifer R. S. Meadows^6^

^1^Department of Clinical Psychology, University of Bergen ^2^Department of Clinical Neuroscience, Karolinska Institutet ^3^Department of Bioinformatics and Integrative Biology, University of Massachusetts Medical School ^4^Department of Genetics, University of North Carolina at Chapel Hill ^5^Department of Biomedicine, University of Bergen ^6^Department of Medical Biochemistry and Microbiology, University of Uppsala ^7^Department of Psychiatry, University of Bergen ^8^Department of Clinical Psychology, Helse Møre og Romsdal HF ^9^Faculty of Computer Science, Dalhousie University ^10^Department of Psychology, Humboldt-Universität zu Berlin ^11^Department of Medical Epidemiology and Biostatisatics, Karolinska Institutet ^12^Department of Clinical Science, University of Bergen

## PGC TS/OCD working group

Nora I. Strom^1,2,3,4^, Zachary F. Gerring^5,6^, Marco Galimberti^7,8^, Dongmei Yu^9,10^, Matthew W. Halvorsen^11^, Abdel Abdellaoui^12^, Cristina Rodriguez-Fontenla^13,14^, Julia M. Sealock^15^, Tim Bigdeli^16,17^, Jonathan R. I. Coleman^18,19^, Behrang Mahjani^20,21^, Jackson G. Thorp^22,23^, Katharina Bey^24^, Christie L. Burton^25^, Jurjen J. Luykx^26,27^, Gwyneth Zai^28,29^, Silvia Alemany^30,31,32^, Christine Andre^33^, Kathleen D. Askland^34^, Nerisa Banaj^35^, Cristina Barlassina^36^, Judith Becker Nissen^37,38^, O. Joseph Bienvenu^39^, Donald Black^40^, Michael H. Bloch^41^, Julia Bäckman^3^, Sigrid Børte^42,43,44^, Rosa Bosch^45,46^, Michael Breen^47,48,49^, Brian P. Brennan^33,50^, Helena Brentani^51^, Joseph D. Buxbaum^20^, Jonas Bybjerg-Grauholm^52^, Enda M. Byrne^53^, Judit Cabana-Dominguez^30,31,32^, Beatriz Camarena^54^, Adrian Camarena^55^, Carolina Cappi^56,57^, Angel Carracedo^58,59,60^, Miguel Casas^61,62^, Maria Cristina Cavallini^63^, Valentina Ciullo^35^, Edwin H. Cook^64^, Jesse Crosby^33,50^, Bernadette A. M. Cullen^65,66^, Elles J. De Schipper^3^, Richard Delorme^67^, Srdjan Djurovic^44,68^, Jason A. Elias^69,70^, Xavier Estivill^71^, Martha J. Falkenstein^33,50^, Bengt T. Fundin^72^, Lauryn Garner^33^, Christina Gironda^33^, Fernando S. Goes^73^, Marco A. Grados^74^, Jakob Grove^4,75,76,77^, Wei Guo^78^, Jan Haavik^79,80^, Kristen Hagen^81,82,83^, Kelly Harrington^84,85^, Alexandra Havdahl^86,87^, Kira D. Höffler^68,88,89^, Ana G. Hounie^90^, Donald Hucks^91^, Christina Hultman^21^, Magdalena Janecka^92,93^, Eric Jenike^33^, Elinor K. Karlsson^94,95^, Kara Kelley^33^, Julia Klawohn^1,96^, Janice E. Krasnow^97^, Kristi Krebs^98^, Christoph Lange^99,100^, Nuria Lanzagorta^101^, Daniel Levey^102,103^, Kerstin Lindblad-Toh^104,95,105^, Fabio Macciardi^106^, Brion Maher^107^, Brittany Mathes^33^, Evonne McArthur^108^, Nathaniel McGregor^109^, Nicole C. McLaughlin^110,111^, Sandra Meier^112^, Euripedes C. Miguel^113^, Maureen Mulhern^24^, Paul S. Nestadt^114^, Erika L. Nurmi^115^, Kevin S. O’Connell^116,117^, Lisa Osiecki^118,119^, Olga Therese Ousdal^79,120^, Teemu Palviainen^121^, Nancy L. Pedersen^21^, Fabrizio Piras^35^, Federica Piras^122^, Sriramya Potluri^33^, Raquel Rabionet^123,124,125^, Alfredo Ramirez^126,127,128,129,130^, Scott Rauch^131^, Abraham Reichenberg^132^, Mark A. Riddle^133^, Stephan Ripke^134,135,136^, Maria C. Rosário^137^, Aline S. Sampaio^138^, Miriam A. Schiele^139^, Anne Heidi Skogholt^140^, Laura G. Sloofman^20^, Jan Smit^141^, María Soler Artigas^30,31,32,142^, Laurent F. Thomas^143,144,145,146^, Eric Tifft^33^, Homero Vallada^147,148^, Nathanial van Kirk^149,150^, Jeremy Veenstra-VanderWeele^151,152^, Nienke N. C. C. Vulink^153^, Christopher P. Walker^154^, Ying Wang^155^, Jens R. Wendland^156^, Bendik S. Winsvold^42,157,158^, Yin Yao^159^, Hang Zhou^160,161,162^, 23andMe Research Team^163^, VA Million Veteran Program^164^, Estonian Biobank^164^, CoGa research team^164^, iPSYCH^164^, HUNT research team^164^, Arpana Agrawal^165^, Pino Alonso^166,167,168,169^, Götz Berberich^170^, Kathleen K. Bucholz^171^, Cynthia M. Bulik^172,21,173^, Danielle Cath^174,175^, Damiaan Denys^176^, Valsamma Eapen^177,178^, Howard Edenberg^179^, Peter Falkai^180,181^, Thomas V. Fernandez^182^, Abby J. Fyer^183,184^, J M. Gaziano^185,186^, Dan A. Geller^187,50^, Hans J. Grabe^188^, Benjamin D. Greenberg^189,190,191^, Gregory L. Hanna^192^, Ian B. Hickie^193^, David M. Hougaard^52,75^, Norbert Kathmann^1^, James Kennedy^29^, Dongbing Lai^194^, Mikael Landén^21,195^, Stéphanie Le Hellard^196,68^, Marion Leboyer^197^, Christine Lochner^198^, James T. McCracken^115^, Sarah E. Medland^199^, Preben B. Mortensen^75,200,201^, Benjamin M. Neale^202,135^, Humberto Nicolini^203,204^, Merete Nordentoft^205,206^, Michele Pato^207^, Carlos Pato^207^, David L. Pauls^50^, John Piacentini^208^, Christopher Pittenger^209^, Danielle Posthuma^210,211^, Josep Antoni Ramos-Quiroga^212,213,214,215^, Steven A. Rasmussen^110^, Margaret A. Richter^216,29^, David R. Rosenberg^217^, Stephan Ruhrmann^218^, Jack F. Samuels^219^, Sven Sandin^21,20^, Paul Sandor^29^, Gianfranco Spalletta^35,220^, Dan J. Stein^221^, S. Evelyn Stewart^222,223,224^, Eric A. Storch^225^, Barbara E. Stranger^226,227^, Maurizio Turiel^228^, Thomas Werge^229,230^, Ole A. Andreassen^231,232^, Anders D. Børglum^4,233,234^, Susanne Walitza^235,236,237^, Kristian Hveem^158,238,239^, Bjarne K. A. Hansen^240,241^, Christian Rück^3^, Nicholas G. Martin^242^, Lili Milani^98^, Ole Mors^243^, Ted Reichborn-Kjennerud^244,245^, Marta Ribasés^30,246,32,142^, Gerd Kvale^82,247^, David Mataix-Cols^3^, Katharina Domschke^139,248^, Edna Grünblatt^249,236,237^, Michael Wagner^24,250,251^, John-Anker Zwart^252,158^, Gerome Breen^253^, Gerald Nestadt^114^, Jaakko Kaprio^254^, Paul D. Arnold^255,256^, Dorothy E. Grice^20^, James A. Knowles^257^, Helga Ask^258,259^, Karin J. H. Verweij^153^, Lea K. Davis^260^, Dirk J. A. Smit^261^, James J. Crowley^11,172,3^, Jeremiah M. Scharf^118,135^, Murray B. Stein^262,263^, Joel Gelernter^264,265^, Carol A. Mathews^266^, Eske M. Derks^267^, Manuel Mattheisen^268,4,2^ & Chris German^163^

^1^Department of Psychology, Humboldt-Universität zu Berlin, Berlin, Germany ^2^Department of Psychiatric Phenomics and Genomics (IPPG), Ludwig-Maximilians University Munich, Munich, Germany ^3^Centre for Psychiatry Research, Department of Clinical Neuroscience, Karolinska Institutet & Stockholm Health Services, Region Stockholm, Stockholm, Sweden ^4^Department of Biomedicine, Aarhus University, Aarhus, Denmark ^5^Department of Mental Health and Neuroscience, Translational Neurogenomics, QIMR Berghofer Medical Research Institute, Brisbane, QLD, Australia ^6^Department of Population Health and Immunity, Healthy Development and Ageing, Walter and Eliza Hall Institute of Medical Research, Melbourne, Victoria, Australia ^7^Department of Psychiatry, Human Genetics, Yale University, New Haven, CT, USA ^8^VA Connecticut Healthcare System, West Haven, CT, USA ^9^Department of Center for Genomic Medicine, Psychiatric and Neurodevelopmental Genetics Unit, Massachusetts General Hospital, Boston, MA, USA ^10^Stanley Center for Psychiatric Research, Broad Institute of MIT and Harvard, Cambridge, MA, USA ^11^Department of Genetics, University of North Carolina at Chapel Hill, Chapel Hill, NC, USA ^12^Department of Psychiatry, Amsterdam University Medical Centers, University of Amsterdam, Amsterdam, The Netherlands ^13^CIMUS (Center for Research in Molecular Medicine and Chronic Diseases), Genomics and Bioinformatics, University of Santiago de Compostela, Santiago de Compostela, A Coruña, Spain ^14^Grupo de Medicina Xenómica, Genetics, FIDIS (Instituto de Investigación Sanitaria de Santiago de Compostela), Santiago de Compostela, A Coruña, Spain ^15^Vanderbilt Genetics Institute, Vanderbilt University, Nashville, TN, USA ^16^Department of Psychiatry and Behavioral Sciences, SUNY Downstate Health Sciences University, Brooklyn, NY, USA ^17^VA NY Harbor Healthcare System, Brooklyn, NY, USA ^18^Social, Genetic and Developmental Psychiatry Centre, Institute of Psychiatry, Psychology, and Neuroscience, King’s College London, London, United Kingdom ^19^National Institute for Health and Care Research Maudsley Biomedical Research Centre, South London and Maudsley NHS Trust, London, United Kingdom ^20^Department of Psychiatry, Icahn School of Medicine at Mount Sinai, New York, NY, USA ^21^Department of Medical Epidemiology and Biostatistics, Karolinska Institutet, Stockholm, Sweden ^22^Mental Health and Neuroscience Program, QIMR Berghofer Medical Research Institute, Brisbane, Queensland, Australia ^23^Faculty of Medicine, School of Biomedical Sciences, University of Queensland, Brisbane, Queensland, Australia ^24^Department of Psychiatry and Psychotherapy, University Hospital Bonn, Bonn, Germany ^25^Department of Neurosciences and Mental Health, Hospital for Sick Children, Toronto, ON, Canada ^26^Department of Psychiatry, Brain, University Medical Center Utrecht, Utrecht, The Netherlands ^27^Second opinion outpatient clinic, GGNet, Warnsveld, The Netherlands ^28^Molecular Brain Science Department, Campbell Family Mental Health Research Institute, Centre for Addiction and Mental Health,, Toronto, ON, Canada ^29^Department of Psychiatry, University of Toronto, Toronto, ON, Canada ^30^Psychiatric Genetics Unit, Group of Psychiatry, Mental Health and Addiction, Vall d’Hebron Research Institute (VHIR), Universitat Autònoma de Barcelona, Barcelona, Spain ^31^Department of Mental Health, Hospital Universitari Vall d’Hebron, Barcelona, Spain ^32^Biomedical Network Research Centre on Mental Health (CIBERSAM), Madrid, Spain ^33^Obsessive-Compulsive Disorder Institute, McLean Hospital, Belmont, MA, USA ^34^Department of Psychiatry & Behavioural Neurosciences, McMaster University, Hamilton, ON, Canada ^35^Laboratory of Neuropsychiatry, IRCCS Santa Lucia Foundation, Rome, Italy ^36^Department of Heath Sciences, University of Milano, Milano, Italy ^37^Department of Child and Adolescent Psychiatry, Aarhus University Hospital, Psychiatry, Aarhus, Denmark ^38^Institute of Clinical Medicine, Health, Aarhus University, Aarhus, Denmark ^39^Department of Psychiatry and Behavioral Sciences, General Hospital Psychiatry, Johns Hopkins University School of Medicine, Baltimore, MD, USA ^40^Departments of Roy J. and Lucille A. Carver College of Medicine, University of Iowa, Iowa City, IA, USA ^41^Department of Child Study Center and Psychiatry, Yale University, New Haven, CT, USA ^42^Department of Research and Innovation, Division of Clinical Neuroscience, Oslo University Hospital, Oslo, Norway ^43^Department of Public Health and Nursing, Faculty of Medicine and Health Sciences, HUNT Center for Molecular and Clinical Epidemiology, Norwegian University of Science and Technology (NTNU), Trondheim, Norway ^44^Department of Medical Genetics, Oslo University Hospital, Oslo, Norway ^45^Department of Child and Adolescent Mental Health, Hospital Sant Joan de Déu, Esplugues de Llobregat, Barcelona, Spain ^46^Instituto de Salut Carlos III, Centro de Investigación Biomédica en Red de Salut Mental (CIBERSAM), Madrid, Spain ^47^Department of Psychiatry, Icahn School of Medicine At Mount Sinai, New York, NY, USA ^48^Seaver Autism Center for Research and Treatment, Icahn School of Medicine At Mount Sinai, New York, NY, USA ^49^The Mindich Child Health and Development Institute, Icahn School of Medicine At Mount Sinai, New York, NY, USA ^50^Department of Psychiatry, Harvard Medical School, Boston, MA, USA ^51^Department of Psychiatry, Universidade De São Paulo, São Paulo, Brazil ^52^Department for Congenital Disorders, Statens Serum Institut, Copenhagen, Denmark ^53^Child Health Research Centre, University of Queensland, Brisbane, Queensland, Australia ^54^Pharmacogenetics Department, Investigaciones Clínicas, Instituto Nacional de Psiquiatría Ramon de la Fuente Muñiz, Mexico City, México ^55^Department of Surgery, Duke, Durham, NC, USA ^56^Department of Psychiatry, Icahn School of Medicine at Mount Sinai Hospital, New York, NY, USA ^57^Department of Psychiatry, University of Sao Paulo, Sao Paulo, Brazil ^58^Center for Research in Molecular Medicine and Chronic Diseases (CiMUS), Genomics and Bioinformatics Group, University of Santiago de Compostela, Santiago de Compostela, Spain ^59^Galiician Foundation of Genomic Medicine, Grupo de Medicina Xenómica, Instituto de Investigación Sanitaria de Santiago -IDIS-, Santiago de Compostela, Spain ^60^Medicina Genómica, Centro de Investigación Biomédica en Red, Enfermedades Raras (CIBERER), Santiago de Compostela, Spain ^61^Programa MIND Escoles, Hospital Sant Joan de Déu, Esplugues de Llobregat, Barcelona, Spain ^62^Departamento de Psiquiatría y Medicina Legal, Universitat Autònoma de Barcelona, Bellaterra, Spain ^63^Department of Psychiatry, Ospedale San Raffaele, Milano, Italy ^64^Department of Psychiatry, University of Illinois Chicago, Chicago, IL, USA ^65^Department of Psychiatry and Behavioral Sciences, Johns Hopkins Medical Institutions, Baltimore, MD, USA ^66^Department of Mental Health, Bloomberg School of Public Health, Baltimore, MD, USA ^67^Child and Adolesccent Psycchiatry Department, APHP, Paris, France ^68^Department of Clinical Science, University of Bergen, Bergen, Norway ^69^Psychiatry, McLean Hospital OCDI, Harvard Medical School, Belmont, MA, USA ^70^Adult Psychological Services, CBTeam LLC, Lexington, MA, USA ^71^Genomics (Quantitative Genomics Laboratories), Esplugues de Llobregat, Barcelona, Spain ^72^Department of Medical Epidemiology and Biostatistics, Center for Eating Disorders Innovation, Karolinska Institutet, Stockholm, Sweden ^73^Department of Psychiatry, Johns Hopkins University, Baltimore, MD, USA ^74^Department of Psychiatry and Behavioral Sciences, Child & Adolescent Psychiatry, Johns Hopkins University, Baltimore, MD, USA ^75^The Lundbeck Foundation Initiative for Integrative Psychiatric Research, iPSYCH, Aarhus, Denmark ^76^Center for Genomics and Personalized Medicine, Aarhus, Denmark ^77^Bioinformatics Research Centre, Aarhus, Denmark ^78^Genetic Epidemiology Research Branch, National Institute of Mental Health, Bethesda, MD, USA ^79^Department of Biomedicine, University of Bergen, Bergen, Norway ^80^Bergen Center for Brain Plasticity, Division of Psychiatry, Haukeland University Hospital, Bergen, Norway ^81^Department of Psychiatry, Møre og Romsdal Hospital Trust, Molde, Norway ^82^Bergen Center for Brain Plasticity, Psychiatry, Haukeland University Hospital, Bergen, Norway ^83^Department of Mental Health, Norwegian University for Science and Technology, Trondheim, Norway ^84^Million Veteran Program (MVP) Coordinating Center, VA Boston Healthcare System, Boston, MA, USA ^85^Department of Psychiatry, Boston University Chobanian & Avedisian School of Medicine, Boston, MA, USA ^86^PsychGen Centre for Genetic Epidemiology and Mental Health, Norwegian Institute of Public Health, Oslo, Norway ^87^Nic Waals Institute, Lovisenberg Diaconal Hospital, Oslo, Norway ^88^Bergen Center for Brain Plasticity, Haukeland University Hospital, Bergen, Norway ^89^Department of Medical Genetics, Dr. Einar Martens Research Group for Biological Psychiatry, Haukeland University Hospital, Bergen, Norway ^90^Department of Psychiatry, University of São Paulo, São Paulo, Brazil ^91^Department of Medicine, Genetic Medicine, Vanderbilt University Medical Center, Nashville, TN, USA ^92^Department of Child and Adolescent Psychiatry, NYU Grossman School of Medicine, New York, USA ^93^Department of Psychiatry, Icahn School of Medicine at Mount Sinai, New York, USA ^94^Department of Bioinformatics and Integrative Biology, University of Massachusetts Medical School, Worcester, MA, USA ^95^Department of Vertebrate Genomics, Broad Institute of MIT and Harvard, Cambridge, MA, USA ^96^Department of Medicine, MSB Medical School Berlin, Berlin, Germany ^97^Department of Psychiatry and Behavioral Sciences, Johns Hopkins University, Baltimore, MD, USA ^98^Estonian Genome Centre, Institute of Genomics, University of Tartu, Tartu, Estonia ^99^Department of Biostatistics, T.H. Chan School of Public Health, Boston, MA, USA ^100^Department of Medicine, Channing Division of Network Medicine, Brigham and Women’s Hospital, Boston, MA, USA ^101^Research Department, Grupo Medico Carracci, Mexico City, Mexico ^102^Department of Psychiatry, Yale University, West Haven, CT, USA ^103^Office of Research & Development, United States Department of Veterans Affairs, West Haven, CT, USA ^104^Department of Medical Biochemistry and Microbiology, Uppsala University, Uppsala, Sweden ^105^SciLifeLab, Uppsala University, Uppsala, Sweden ^106^Department of Psychiatry, University of California, Irvine (UCI), Irvine, CA, USA ^107^Department of Mental Health, Johns Hopkins Bloomberg School of Public Health, Baltimore, MD, USA ^108^Vanderbilt Genetics Institute, Vanderbilt University Medical Center, Nashville, TN, USA ^109^COBRE Center for Neuromodulation, Butler Hospital, Providence, RI, USA ^110^Department of Psychiatry & Human Behavior, Alpert Medical School, Brown University, Providence, RI, USA ^111^Butler Hospital, Providence, RI, USA ^112^Department of Psychiatry, Dalhousie University, Halifax, NS, Canada ^113^Department of Psychiatry, Faculdade de Medicina da Universidade de São Paulo, São Paulo, Brazil ^114^Department of Psychiatry and Behavioral Science, Johns Hopkins University, Baltimore, MD, USA ^115^Department of Psychiatry and Biobehavioral Sciences, Division of Child and Adolescent Psychiatry, University of California, Los Angeles, Los Angeles, CA, USA ^116^Department of Clinical Medicine, Division of Mental Health and Addiction, Oslo University Hospital, Oslo, Norway ^117^NORMENT, University of Oslo, Oslo, Norway ^118^Department of Psychiatry, Massachusetts General Hospital, Boston, MA, USA ^119^Psychiatric and Neurodevelopmental Genetics Unit, Harvard Medical School, Boston, MA, USA ^120^Department of Biomedicine, Haukeland University Hospital, Bergen, Norway ^121^Institute for Molecular Medicine Finland - FIMM, University of Helsinki, Helsinki, Finland ^122^Department of Clinical Neuroscience and Neurorehabilitation, Neuropsychiatry Laboratory, IRCCS Santa Lucia Foundation, Rome, Italy ^123^Department of Genetics, microbiology and statistics, IBUB, Universitat de Barcelona, Barcelona, Spain ^124^CIBERER, Centro de investigación biomédica en red, Madrid, Spain ^125^Department of Human Molecular Genetics, Institut de Recerca Sant Joan de Déu, Esplugues de Llobregat, Barcelona, Spain ^126^Department of Psychiatry and Psychotherapy, Division of Neurogenetics and Molecular Psychiatry, Faculty of Medicine and University Hospital Cologne, University of Cologne, Cologne, Germany ^127^Department of Neurodegenerative Diseases and Geriatric Psychiatry, University Hospital Bonn, Medical Faculty, Bonn, Germany ^128^DZNE Bonn, German Center for Neurodegenerative Diseases (DZNE), Bonn, Germany ^129^Department of Psychiatry and Glenn Biggs Institute for Alzheimer’s and Neurodegenerative Diseases, UT Health San Antonio, San Antonio, TX, USA ^130^Cologne Excellence Cluster for Stress Responses in Ageing-associated diseases (CECAD), University of Cologne, Cologne, Germany ^131^Department of Psychiatry, McLean Hospital, Harvard Medical School, Belmont, MA, USA ^132^Department of Mental disorders, Norwegian Institute of Public Health, New York, NY, USA ^133^Department of Psychiatry and Behavioral Sciences, Child and Adolescent, Johns Hopkins University School of Medicine, Baltimore, MD, USA ^134^Department of Psychiatry and Psychotherapy, Charité - Universitätsmedizin, Berlin, Germany ^135^Stanley Center for Psychiatric Research, Broad Institute of Harvard and MIT, Cambridge, MA, USA ^136^site Berlin-Potsdam, German Center for Mental Health (DZPG), Berlin, Germany ^137^Department of Psychiatry, Child and Adolescent Psychiatry Unit (UPIA), Federal University of São Paulo (UNIFESP), São Paulo, Brazil ^138^Department of Neurosciences and Mental Health, Medical School, Federal University of Bahia, Salvador, Brazil ^139^Department of Psychiatry and Psychotherapy, Faculty of Medicine, University of Freiburg, Medical Center - University of Freiburg, Freiburg, Germany ^140^Department of Public Health and Nursing, HUNT Center for Molecular and Clinical Epidemiology, Trondheim, Norway ^141^Department of Psychiatry, Faculty of Medicine, Location Vumc, Amsterdam UMC, University of Amsterdam, Amsterdam, The Netherlands ^142^Department of Genetics, Microbiology, and Statistics, Faculty of Biology, Universitat de Barcelona (UB), Barcelona, Spain ^143^Department of Clinical and Molecular Medicine, Norwegian University of Science and Technology, Trondheim, Norway ^144^Department of Public Health and Nursing, K. G. Jebsen Center for Genetic Epidemiology, Norwegian University of Science and Technology, Trondheim, Norway ^145^BioCore - Bioinformatics Core Facility, Norwegian University of Science and Technology, Trondheim, Norway ^146^Clinic of Laboratory Medicine, St.Olavs Hospital, Trondheim University Hospital, Trondheim, Norway ^147^Department of Psychiatry, Universidade de Sao Paulo, São Paulo, Brazil ^148^Department of Molecular Medicine and Surgery, CMM, Karolinska Institutet, Stockholm, Sweden ^149^OCD Institute, Division of Depression and Anxiety, McLean Hospital, Belmont, MA, USA ^150^Department of Psychiatry, Harvard Medical School, Belmont, MA, USA ^151^Department of Psychiatry, Division of Child and Adolescent Psychiatry, Columbia University, New York, NY, USA ^152^Department of Child and Adolescent Psychiatry, New York State Psychiatric Institute, New York, NY, USA ^153^Department of Psychiatry, Amsterdam UMC, University of Amsterdam, Amsterdam, The Netherlands ^154^Human Genetics Institute of New Jersey, Rutgers University, Piscataway, NJ, USA ^155^Department of Neurology, The Johns Hopkins University School of Medicine, Baltimore, MD, USA ^156^Laboratory of Clinical Science, NIMH Intramural Research Program, Bethesda, MD, USA ^157^Department of Neurology, Oslo University Hospital, Oslo, Norway ^158^HUNT Center for Molecular and Clinical Epidemiology, Department of Public Health and Nursing, Faculty of Medicine and Health Sciences, Norwegian University of Science and Technology, Trondheim, Norway ^159^Department of Computional Biology, Institute of Life Science, Fudan University, Fudan, China ^160^Department of Psychiatry, Yale School of Medicine, New Haven, CT, USA ^161^Department of Psychiatry, Veterans Affairs Connecticut Healthcare System, West Haven, CT, USA ^162^Section of Biomedical Informatics and Data Science, Yale School of Medicine, New Haven, CT, USA ^163^23andMe, Inc., Sunnyvale, CA, USA ^164^Million Veteran Program^165^Department of Psychiatry, Washington University School of Medicine, Saint Louis, MO, USA ^166^Department of Psychiatry, OCD Clinical and Research Unit, Bellvitge Hospital, Barcelona, Spain ^167^Department of Clinical Sciences, University of Barcelona, Barcelona, Spain ^168^Department of Psychiatry and Mental Health, Bellvitge Biomedical Research Institute IDIBELLL, Barcelona, Spain ^169^CIBERSAM, Mental Health Network Biomedical Research Center, Madrid, Spain ^170^Psychosomatic Department, Windach Hospital of Neurobehavioural Research and Therapy, Windach, Germany ^171^Department of Psychiatry, Washington U. School of Medicine, St Louis, MO, USA ^172^Department of Psychiatry, University of North Carolina at Chapel Hill, Chapel Hill, NC, USA ^173^Department of Nutrition, University of North Carolina at Chapel Hill, Chapel Hill, NC, USA ^174^Departments of Rijksuniversiteit Groningen and Psychiatry, University Medical Center Groninge, Groningen, The Netherlands ^175^Department of Specialized Training, Drenthe Mental Health Care Institute, Groningen, The Netherlands ^176^Department of Psychiatry, Institute of The Royal Netherlands Academy of Arts and Sciences (NIN-KNAW), Amsterdam, The Netherlands ^177^Discipline of Psychiatry and Mental Health, School of Clinical Medicine, UNSW, Sydney, NSW, Australia ^178^Academic Unit of Child Psychiatry South-West Sydney (AUCS), South-West Sydney Clinical School, SWSLHD & Ingham Institute, Sydney, NSW, Australia ^179^Department of Biochemistry and Molecular Biology, Indiana University School of Medicine, Indianapolis, IN, USA ^180^Department of Psychiatry and Psychotherapy, University Hospital LMU, Munich, Germany ^181^Department of Psychiatry, Max Planck Institute, Munich, Germany ^182^Child Study Center and Psychiatry, Yale School of Medicine, New Haven, CT, USA ^183^Department of Psychiatry, New York State Psychiatric Institute, New York, NY, USA ^184^Department of Psychiatry, Columbia University Medical Center, New York, NY, USA ^185^Department of Medicine, VA Boston Healthcare System, Boston, MA, USA ^186^Department of Medicine, Mass General Brigham, Boston, MA, USA ^187^Department of Psychiatry, Child Psychiatry, Massachusetts General Hospital, Boston, MA, USA ^188^Department of Psychiatry & Psychotherapy, University Medicine Greifswald, Greifswald, Germany ^189^COBRE Center on Neuromodulation, Butler Hospital, Providence, RI, USA ^190^Center for Neurorestoration and Neurotechnology, VA Providence Healthcare System, Providence, USA ^191^Department of Psychiatry and Human Behavior, Alpert Medical School, Brown University, Providence, RI, USA ^192^Department of Psychiatry, Child and Adolescent Psychiatry, University of Michigan, Ann Arbor, MI, USA ^193^Brain and Mind Centre, The University of Sydney, Sydney, NSW, Australia ^194^Department of Medical and Molecular Genetics, Indiana University School of Medicine, Indianapolis, IN, USA ^195^Institute of Neuroscience and Physiology, Department of Psychiatry and Neurochemistry, University of Gothenburg, Gothenburg, Sweden ^196^Bergen Center for brain plasticity, Division of Psychiatry, Haukeland University Hospital, Bergen, Norway ^197^Department of Addictology and Psychiatry, Univ Paris Est Créteil, AP-HP, Inserm, Paris, France ^198^Department of Psychiatry, SA MRC Unit on Risk and Resilience in Mental Disorders, Stellenbosch University, Stellenbosch, South Africa ^199^Department of Mental Health, Psychiatric Genetics, QIMR Berghofer Medical Research Institute, Brisbane, Queensland, Australia ^200^National Centre for Register-based Research, Aarhus University, Aarhus, Denmark ^201^Centre for Integrated Register-based Research, Aarhus University, Aarhus, Denmark ^202^Analytic and Translational Genetics Unit, Massachusetts General Hospital, Boston, MA, USA ^203^Department of Psychiatry, Psychiatry, Carracci Medical Group, Mexico City, México ^204^Psiquiatría, Instituto Nacional de Medicina Genómica, Mexico City, México ^205^Mental Health Center Copenhagen, Copenhagen Research Center for Mental Health, Mental Health services in the Capital Region of Denmark, Copenhagen, Denmark ^206^Faculty of Health and Medical Sciences, Department of Clinical Medicine, University of Copenhagen, Copenhagen, Denmark ^207^Department of Psychiatry, Rutgers University, Piscataway, NJ, USA ^208^Department of Psychiatry and Biobehavioral Sciences, Child and Adolescent Psychiatry, UCLA Semel Institute for Neuroscience and Human Behavior, Los Angeles, CA, USA ^209^Department of Psychiatry, Yale University, New Haven, CT, USA ^210^Department of Complex Trait Genetics, Vrije Universiteit Amsterdam, Center for Neurogenomics and Cognitive Research, Amsterdam Neuroscience, Amsterdam, The Netherlands ^211^Department of Child and Adolescent Psychiatric, Section Complex Trait Genetics, VU Medical Center Amsterdam, Amsterdam, The Netherlands ^212^Department of Psychiatry, Hospital Universitari Vall d’Hebron, Barcelona, Spain ^213^Group of Psychiatry, Mental Health and Addictions, Psychiatric Genetics Unit, Vall d’Hebron Research Institute (VHIR), Barcelona, Spain ^214^Biomedical Network Research Centre on Mental Health (CIBERSAM), Barcelona, Spain ^215^Department of Psychiatry and Forensic Medicine, Universitat Autònoma de Barcelona, Barcelona, Spain ^216^Department of Psychiatry, Sunnybrook Health Sciences Centre, Toronto, ON, Canada ^217^Department of Psychiatry and Behavioral Neurosciences, Child and Adolescent Psychiatry, Wayne State University School of Medicine, Detroit, MI, USA ^218^Department of Psychiatry and Psychotherapy, University of Cologne, Cologne, Germany ^219^Department of Psychiatry and Behavioral Sciences, The Johns Hopkins University School of Medicine, Baltimore, MD, USA ^220^Department of Psychiatry and Behavioral Sciences, Division of Neuropsychiatry, Baylor College of Medicine, Houston, TX, USA ^221^Dept of Psychiatry & Neuroscience Institute, SAMRC Unit on Risk & Reslience in Mental Disorders, University of Cape Town, Cape Town, Western Cape, South Africa ^222^Department of Psychiatry, University of British Columbia, Vancouver, BC, Canada ^223^British Columbia Children’s Hospital Research Institute, Vancouver, BC, Canada ^224^British Columbia Mental Health and Substance Use Services Research Institute (BCMHSUS), Vancouver, BC, Canada ^225^Department of Psychiatry and Behavioral Sciences, Baylor College of Medicine, Houston, TX, USA ^226^Department of Pharmacology, Northwestern University Feinberg School of Medicine, Chicago, IL, USA ^227^Center for Genetic Medicine, Northwestern University Feinberg School of Medicine, Chicago, IL, USA ^228^Department of Cardiology, University of Milan, Milan, Italy ^229^Institute of Biological Psychiatry, Mental Health Center Sct. Hans, Copenhagen University Hospital, Mental Health Services (RHP), Copenhagen, Denmark ^230^Institute of Clinical Medicine, University of Copenhagen, Copenhagen, Denmark ^231^Institute of Clinical Medicine, NORMENT Centre, University of Oslo, Oslo, Norway ^232^Division of Mental Health and Addiction, Center for Precision Psychiatry, Oslo University Hospital, Oslo, Norway ^233^The Lundbeck Foundation Initiative for Integrative Psychiatric Research, iPSYCH, Aarhus University, Aarhus, Denmark ^234^Center for Genomics and Personalized Medicine, CGPM, Aarhus University, Aarhus, Denmark ^235^Department of Child and Adolescent Psychiatry and Psychotherapy, Psychiatric University Hospital Zurich (PUK), University of Zurich, Zürich, Switzerland ^236^Neuroscience Center Zurich, University of Zurich and the ETH Zuric, Zurich, Switzerland ^237^Zurich Center for Integrative Human Physiology, University of Zurich, Zurich, Switzerland ^238^HUNT Research Center, Department of Public Health and Nursing, Faculty of Medicine and Health Sciences, Norwegian University of Science and Technology, Trondheim, Norway ^239^Department of Research, Innovation and Education, St. Olavs Hospital, Trondheim University Hospital, Trondheim, Norway ^240^Bergen Center for Brain Plasticity (BCBP), Psychiatry, Haukeland University Hospital, Bergen, Norway ^241^Centre for Crisis Psychology, Psychology, University of Bergen, Bergen, Norway ^242^Department of Genetic Epidemiology, QIMR Berghofer Medical Research Institute, Brisbane, Queensland, Australia ^243^Psychosis Reasearch Unit, Aarhus University Hospital - Psychiatry, 8200 Aarhus N, Denmark ^244^Department of Mental Disorders, Norwegian Institute of Public Health, Oslo, Norway ^245^Institute of Clinical Medicine, University of Oslo, Oslo, Norway ^246^Department of Mental Health, Hospital Universitari Vall d’Hebron, Barcelona, Spain ^247^Department of Clinical Psychology, Faculty of Psychology, University of Bergen, Bergen, Vestland ^248^German Center for Mental Health (DZPG), Partner Site Berlin, Berlin, Germany ^249^Department of Child and Adolescent Psychiatry and Psychotherapy, Psychiatric University Hospital Zurich (PUK), University of Zurich, Zürich, Schweiz ^250^Department of Neurodegenerative Diseases and Geriatric Psychiatry, University Hospital Bonn, Bonn, Germany ^251^German Center for Neurodegenerative Diseases, DZNE, Bonn, Germany ^252^Department of Research and Innovation, Clinical Neuroscience, Oslo University Hospital and University of Oslo, Oslo, Norway ^253^Social, Genetic, and Developmental Psychiatric Centre, Institute of Psychiatry, Psychology, and Neuroscience, King’s College London, London, United Kingdom ^254^Institute for Molecular Medicine FIMM, University of Helsinki, Helsinki, Finland ^255^Department of Psychiatry, The Mathison Centre for Mental Health Research & Education, Cumming School of Medicine, University of Calgary, Calgary, AB, Canada ^256^Program in Genetics and Genome Biology, Hospital for Sick Children, Toronto, ON, Canada ^257^Department of Genetics, Human Genetics Institute of New Jersey, Rutgers University, Piscataway, NJ, USA ^258^PsychGen Center for Genetic Epidemiology, Norwegian Institute of Public Health, Oslo, Norway ^259^PROMENTA Research Center, Department of Psychology, University of Oslo, Oslo, Norway ^260^Department of Medicine, Division of Genetic Medicine, Vanderbilt University Medical Center, Nashville, TN, USA ^261^Department of Psychiatry, Amsterdam UMC location AMC, Amsterdam, The Netherlands ^262^Psychiatry Service, VA San Diego Healthcare System, San Diego, CA, USA ^263^Department of Psychiatry and School of Public Health, University of California San Diego, La Jolla, CA, USA ^264^Department of Psychiatry, Human Genetics (Psychiatry), Yale University School of Medicine, West Haven, CT, USA ^265^Department of Psychiatry, Veterans Affairs Connecticut Healthcare Center, West Haven, CT, USA ^266^Psychiatry and Genetics Institute, Center for OCD, Anxiety and Related Disorders, University of Florida, Gainesville, FL, USA ^267^Department of Mental Health and Neuroscience, QIMR Berghofer, Brisbane, Australia ^268^Department of Community Health and Epidemiology and Faculty of Computer Science, Dalhousie University, Halifax, NS, Canada

## Methods

### Animal approval

Dogs included in GWAS were all privately owned pets enrolled by their owners in Darwin’s Ark, a community-science research program. The study protocol was approved by the UMass Chan Medical School Institutional Animal Care and Use Committee.

Canine brain tissue samples were purchased from Zyagen, USA (Agreement Numbers #13256 and #13403) (**Supplementary Methods 1**). According to the supplier’s documentation, tissues were derived from animals maintained in facilities operating in compliance with the SDA Animal Welfare Act (AWA) and American Kennel Club standards.

### Phenotypic data and Exploratory factor analysis

Exploratory factor analysis (EFA) was performed on survey responses to 14 polytomous items related to repetitive and compulsive behaviour (**Supplementary Table 1**; **Supplementary Methods 2**). These were the full responses from 12,989 pet dogs within the Darwin’s Ark platform^38^ (https://darwinsark.org; Datafreeze: July 15, 2022). Suitability for

EFA was assessed using Bartlett’s test of sphericity^138^ (χ² = 39597.18, df = 91, P = 0) and Kaiser–Meyer–Olkin MSA^139^ (overall = 0.77; range 0.67–0.90). The number of factors was determined using scree test^140^ and parallel analysis^141^ (nFactors::parallel, 1,000 simulations)^142^, both indicating four factors; however, a three-factor solution was retained because all factors had SS-loadings > 1. EFA was conducted in R (v4.5.3) using the psych package (v2.6.5) with polychoric correlations, principal factor extraction (fm = ‘pa’), varimax rotation, max.iter = 200, SMC = FALSE. Reliability was assessed using Cronbach’s alpha (α = 0.75, 95% CI 0.75–0.76, 1,000 bootstrap samples). Factor scores for GWAS were derived using item response theory (IRT) graded response models in mirt (v1.47), with model fit evaluated via M2* (or C2) and item-fit diagnostics; EAP or MAP scoring was used per factor based on standard-error distributions. Unlike EFA, which required complete responses across all 14 items, IRT scoring could accommodate dogs with partial item responses (up to 25,031 dogs received an F1 score); scores were retained only for dogs with at least two answered items for that factor (F1: N = 15,505; F2: N = 19,002; F3: N = 15,525). GWAS was further restricted to the subset of these dogs with genotype data (see below). Full loading matrices, fit indices, are provided in **Supplementary Tables 16-17**.

### Genotypic data

Genotypic analyses used a hybrid dog reference genome consisting of the autosomes and X chromosome of UU_Cfam_GSD_1.0/CanFam4 (GCF_011100685.1) and the ROS_Cfam_1.0 (GCF_014441545.1) Y chromosome contigs in order to match the Dog10K phased imputation panel of 1,929 canines^46^ (**Supplementary Methods 3**).

Three data types were used, high-coverage WGS (n = 11, PRJNA683923, **Supplementary Table 3** for SRA IDs), low-pass sequencing (n = 3,032, PRJNA675863; Gencove), and array data (n = 411, Axiom Canine Genotyping Array Sets A and B, CanFam3.1 reference). High-coverage WGS was mapped as per the Dog10K pipeline (https://github.com/jmkidd/dogmap) with the following tools (bwa-mem2 v2.1, GATK^143^ v4.4.0.0, SAMtools^144^ v1.20, htslib v1.20) and SNPs called with hard filter thresholds (QD < 2.0, QUAL < 30.0, SOR > 3.0, FS > 60.0, MQ < 40.0, MQRankSum < -12.5, ReadPosRankSum < -8.0). The same mapping tools and process were used for the low-pass samples, depth estimated with SAMtools v1.20 “coverage” and found to range between 0.0-4.8X. For the array data, liftover was performed with the LiftOver chain file (canFam3.1 to CanFam4), and the following quality control filters applied with PLINK^145^ v1.90b7.7 QC filters (--geno 0.05, --hwe midp 1e-15, non-ACGT exclusion, --maf 0.01, chr-mismatch removal) (**Supplementary Methods 3**).

Low-pass and array data were imputed with the Dog10K panel (**Supplementary Methods 3**). For low-pass data, GLIMPSE^146^ v1.1.1 and BCFtools^147^ v1.20 were used for SNP extraction, genotype likelihood calculation, chunking, phasing, and ligation. Variants with INFO ≥ 0.8 were retained (SNPs n = 14,274,129). For Array data conform-gt harmonization was performed prior to imputation with Beagle 5.0^148^ (30 burn-in, 40 iterations). Variants with DR2 ≥ 0.8 (14,042,437 variants).

Before merging genotype sets, genotype concordance and low-coverage minimum depth thresholds were determined (**Supplementary Methods 3**). GATK^143^ v4.4.0.0 GenotypeConcordance (Picard v3.1.1, Java 17) was used for all calculations. First, concordance was determined for high-vs low-pass (n = 11 pairs, concordance > 0.97, **Supplementary Table 3**). Next a minimum depth for low-pass was determined with downsampling (10 high-coverage samples reduced to 0.1-1.0X) and set to ≥ 0.3X to ensure a non-reference discordance ≥ 0.95. Finally, array vs low-pass concordance was assessed (5 matched + 1 negative control; sensitivity/concordance > 0.96 vs 0.27/0.53). The final merged dataset was created with PLINK^145^ v1.90b7.7 and the following settings, filters (--maf 0.001, --hwe midp 1e-20 per dataset), merge (--bmerge merge-mode 1, merge-equal-pos) and flip-scan LD QC (iterative removal of problematic variants). A final quality control step (--geno 0.05, --hwe midp 1e-20), yielded 12,518,078 biallelic SNPs from a set of 3,455 samples (**Supplementary Methods 3**).

### Genome-wide association analyses

KING^149^ v2.3.2 was used to assess sample relatedness with the input of LD-pruned genotypes (PLINK^145^ v1.90b7.7, --indep-pairwise 50 5 0.2). Duplicates were removed (kinship ≥ 0.5; n = 13), leaving 3,328 samples for downstream analysis. This number was reduced to 2,584 samples (∼9.9M SNPs after MAF ≥ 0.01 filtering) as not all genotyped samples had complete survey responses for each GWAS phenotype. For this set, the mean age was 5.9 years (range 1–22) and 18% were registered purebred. For each GWAS run, continuous factor traits (n = 3) and ordinal categorical item traits (n = 14), and the control ordinal categorical trait size, the autosomal SNP count, age and number of each sex can be found in **Supplementary Table 2**.

As the genotyping batch was found not to impact GWAS results, minimum models were run for each trait assessed (**Supplementary Methods 4**). Analyses were only performed on autosomes. For continuous factor traits, GCTA^150^ v1.94.1 MLMA-LOCO was run, retaining samples with complete data (PLINK^145^ --fam), filtering for MAF ≥ 0.01, and with sex and age as covariates. For size, GCTA v1.94.1 MLMA-LOCO was run as above, with only sex as a covariate. For categorical behavioural traits, POLMM^48^ was run with GRAB v0.2.5^151^ in R v4.5.3. First, a GRM was built from LD-pruned data (PLINK --indep-pairwise 50 5 0.2), the null model fit with GRAB (GRAB.NullModel, method = “POLMM”, traitType = “ordinal”, LOCO = FALSE, tolTau = 0.2, tolBeta = 0.1) including the phenotype, sex and age. An association analysis was then performed with GRAB.Marker on the non-LD pruned genotypes. The Manhattan and QQ plots of each result were visualised using the qqman^152^ package in R v4.5.3. Genome-wide and suggestive thresholds were set to p ≤ 4×10^-7^ (empirically derived^53^) and p ≤ 1×10^-6^, respectively.

### Credible set fine-mapping

Prior to fine-mapping, low-pass sequencing BAMs were inspected in IGV^153^ v2.16.0 to confirm both alleles were represented by reads as sites representative for significant and suggestive GWAS peaks. Regions of interest were defined with PLINK^145^ v1.90b7.7 clumping (--clump-kb 250, --clump-p1 1×10^-6^, --clump-p2 1×10^-5^, --clump-r2 0.5, --clump-range restricted to genes from the Dog10K^46^ NCBIv106 gene annotations). Variant credible sets were determined by applying SuSiE^154^ (Sum of Single Effects) to each clump ±100kb using finemapper.py in the polyfun environment; where no credible set (CS) was identified within this widened window, SuSiE was instead applied to the clump alone. Genes were assigned to each region from its CS interval specifically using the nearest gene within ±1 Mb when the interval contained none; this keeps gene counts tied to the fine-mapped CS interval. Clumps from different factors or items were fine-mapped independently. For the purpose of reporting the number of distinct genomic regions, clumps whose boundaries — widened by ±100kb — overlapped were counted as a single region, classified as significant if any of its constituent clumps reached genome-wide significance, and as suggestive only if none did.

Regions of interest visualised in R using pairwise LD (r²) calculated between the lead SNP and surrounding markers (PLINK v1.90b7.7 ; --r2 --ld-snp, --ld-window-kb 10000, --ld-window 99999, --ld-window-r2 0). Plots were overlaid with fine-scale recombination rates derived from Dog10K data^155^.

### SNP-based heritability

The estimated variance explained by all SNPs was calculated using GCTA^150^ v1.94.1 GREML (--reml) with full and LD-stratified GRMs (LD scores in 250 kb regions, with 10,000 kb blocks and 5,000 kb overlap; four LD-stratified GRMs), covariates sex and age. The same analyses were applied to both continuous and categorical traits, as according to POLMM developers, the variance components from that method can not be used for accurate estimates of heritability.

### Statistical power comparison between dogCD and human OCD GWAS

To place the dogCD and human OCD discoveries on a common power scale, we first converted the human case-control design (53,660 cases, 2,044,417 controls) into an effective sample size to account for the unbalanced case:control ratio: Neff = 4 / (1/Ncases + 1/Ncontrols). For each locus, we estimated the phenotypic variance explained (R2) from its reported p-value as R2 = χ2/(χ2+N), where χ2 = qchisq(1−p, df = 1); this test-statistic-based estimate does not assume a particular effect-size scale, so it applies equally to dogCD’s continuous and ordinal traits and to human OCD’s case-control design, using Neff in place of N for the latter. Power to reach each study’s discovery threshold (p ≤ 4×10⁻⁷ for dogCD; p ≤ 5×10⁻⁸ for human OCD) was then computed from the non-central chi-squared distribution (1 d.f.) as a function of sample size and R2, using each dogCD trait’s own sample size for its lead loci and the minimum trait sample size (N = 2,322) for the representative power curve shown in the figure. All analyses were performed in R using the tidyverse package.

### Candidate cis-regulatory element identification from dog brain

See **Supplementary Methods 1** for full protocol. Commercial brain samples from eight regions (anterior cingulate gyrus, cerebellum, frontal lobe, hypothalamus, occipital cortex, striatum, temporal cortex and thalamus) were obtained from each of four beagles (two males, two females; Zyagen) (**Supplementary Table 18**). Tissue fragments (25-50mg) were homogenised using Dounce homogenizer, filtered through a 30-µm filter (Miltenyi Biotec) and isolated as a nuclei suspension for downstream analysis.

Open chromatin was profiled via bulk ATAC-seq using custom-assembled Tn5 complexes based on established protocols^156^ (**Supplementary Table 19**). Cleared nuclei were transposed and purified libraries quantified (**Supplementary Table 20**). Histone marks H3K4me3 and H3K27me3 were profiled using the CUTANA ChIC/CUT&RUN Kit (EpiCypher) with specific antibodies (MA5-11199 and PA5-120811, Thermo Fisher Scientific; **Supplementary Tables 21-22**). Active enhancers were profiled by ChIP-seq for H3K27ac using antibody A7253 (Abclonal) after cross-linking and sonication with a Bioruptor Pico (**Supplementary Tables 21 and 23**).

Libraries were sequenced on an Illumina NovaSeq 6000 (paired-end 2x150-bp). ATAC-seq and ChIP-seq data were processed via ENCODE pipelines (v0.3.0^156,157^ and v1.9(https://github.com/ENCODE-DCC/chip-seq-pipeline2), respectively), trimmed with Trimmomatic^158^ v0.41 and aligned to the canine UU_Cfam_GSD_1.0/CanFam4 reference genome^54^ using BWA-MEM v0.7.17. Following duplicate removal and MAPQ filtering (< 30), peaks were called using MACS2^159^ with strict library complexity thresholds (NRF, PBC1, PBC2) and matched input for ChIP-seq (**Supplementary Table 24**). Replicate peak-calling reproducibility was enforced across all epigenetic marks.

ChromHMM^160^ v1.27 was used to model genome-wide chromatin states across all brain regions using a 200-bp bin size. A nine-state model was selected as optimal using the CompareModels function. Functional elements were classified based on combinatorial marking; promoters (presence of H3K4me3; states 1, 4, 5 and 6), active enhancers (presence of H3K27ac, absence of H3K4me3; states 2 and 8), open chromatin (presence of ATAC signal only; state 3) and repressed regions (presence of H3K27me3; state 9) with the remaining state (7) regions lacking detectable signal due to the absence of any peaks (**Supplementary Tables 25-26**). Region-specific overlaps and genomic coverages were analysed in R. State enrichment near expressed hypothalamic genes^54^, Zoonomia phastCons conserved elements^161^ and EpiC Dog^105^ regulatory elements (UCSC liftOver^162^ from CanFam3.1 to UU_Cfam_GSD_1.0/CanFam4) were computed with BEDtools^163^ and the R package ‘valr’^164^.

### GWAS functional annotation and signal enrichment

From the compulsive GWASs, genome-wide significant, suggestive and credible set implicated SNPs were annotated for Dog10K^46^ allele frequencies (N = 1,872 dogs), EpiC Dog^105^ active cCREs brain (2 tissues) and non-brain (9 tissues), active cCREs brain (8 tissues, above), Dog10K curated NCBI v106 gene models^46^ and Zoonomia phyloP constraint scores^165^(sites with a score above the 5% FDR phyloP threshold of 2.27 classed as constrained).

Enrichment of genes taken forward to NetColoc among brain regions and cell types was evaluated using fgsea v. 1.36.2 in R v4.5.3. Genes were ranked by specificity, calculated as mean expression in each cell type or region divided by mean expression across all cells, using Seurat 5.5.1. Significance was assessed with fgsea’s adaptive multilevel permutation test, and empirical P-values were corrected using the Benjamini–Hochberg method; an adjusted P<0.05 was considered significant.

To test whether dogCD GWAS regions are enriched in brain-active regulatory elements relative to dog size GWAS regions, we calculated the base-pair overlap of dogCD and dog size GWAS clump regions (+/-100kb) with EpiC Dog brain versus non-brain cCREs using Fisher’s exact tests. The same analysis was repeated independently for each of the eight profiled brain regions using our own cCRE catalogue; p-values were FDR-corrected across regions.

### Network propagation and colocalization

Human–dog 1:1 orthologues were obtained from TOGA^55^. Seed genes were defined independently for each species (**Supplementary Table 6**). For human OCD, we used the 48 genes implicated by at least two gene-based mapping approaches in the largest published human OCD GWAS^1^. For dogCD, fine-mapped credible intervals were intersected with TOGA 1:1 orthologues; all genes within a credible interval were included, with the nearest orthologue within ±1 Mb used when an interval contained no annotated genes. This yielded 83 dogCD seed genes after excluding four dogCD genes that had no 1:1 human orthologues.

Network proximity scores (NPS) were calculated for all genes in the human (NPS_h_) and dog (NPS_d_) networks by propagating the respective seed gene sets through PCNet2.0 — a composite human protein–protein interaction network of ∼3.85 million pairwise associations among 19,267 genes — using the NetColoc framework^23,24^. Each gene received a combined score NPS_hd_ = NPS_h_ × NPS_d_. Thresholds of NPS_h_ > 1.5, NPS_d_ > 1.5, and NPS_hd_ > 3.0 were established by a sensitivity analysis across threshold combinations (**Supplementary Table 7**; p = 2.1 × 10^-5^ for the selected thresholds) and confirmed with an independent permutation test at the selected thresholds: an empirical null built from 10,000 bin-controlled permutations of NPS_d_, with NPS_h_ held fixed (calculate_expected_overlap), against which the observed network size exceeded chance expectation (p = 2.3 × 10^-5^, z-test). Because each permutation analysis draws its own independent random sample of null orderings, the two p-values are expected to be close but not necessarily identical.

To assess specificity, dogCD seed genes were colocalized with networks derived from human GWAS for depression^89^, schizophrenia^166^, rheumatoid arthritis^62^, type 2 diabetes^63^, and height^60^, and dog size GWAS seed genes were colocalized with the human height network. Colocalization significance for control-trait comparisons was assessed using the same network-propagation framework, with a gene-removal permuted null (1,000 replicates; **Supplementary Table 10**).

Single species networks were defined by genes with NPS >3.0 for dogCD, OCD, depression and schizophrenia, respectively. Cross-species networks used the same NPS_h_ > 1.5 and NPS_d_ > 1.5 thresholds as the dogCD-OCD network; the combined-score threshold (NPS_hd_) was selected independently for each pairing via sensitivity analysis, giving NPS_hd_ > 3.0 for dogCD-schizophrenia (matching dogCD-OCD) and NPS_hd_ > 4.0 for dogCD-depression.

Community detection was performed with HiDeF (v1.1beta, via cdapsutil.CommunityDetection, --maxres 20). Sub-network edges were transformed to cosine similarity scores (network_colocalization.transform_edges; edge weight threshold 0.95), reflecting the extent to which two genes share interactors. Networks were visualised in Cytoscape v3.10.2.

### GO enrichment and community annotation

Network communities were annotated using Fisher’s exact tests against GO Biological Process terms^68,69^. Semantically related terms were collapsed into higher-order functional modules using GOSemSim (Wang semantic similarity) semantic similarity clustering^70^, with average-linkage hierarchical clustering cut at a similarity threshold of 0.6, to reduce redundancy. Enrichment significance thresholds (p < 1 × 10^-5^) and fold-enrichment values were compared between single-species (human-only OCD, depression, schizophrenia) and colocalized (OCD/dogCD) networks to assess the gain in biological resolution from cross-species colocalization. To quantify similarity between each pair of networks’ significantly enriched GO Biological Process terms (p < 1 × 10^-5^), we computed an equal-weighted semantic similarity score using the Wang method in GOSemSim^70^: each network’s terms were matched to their best counterpart in the other network, and the two resulting directional means were averaged, giving both networks equal weight regardless of their number of terms. Scores were chance-corrected as (observed − expected)/(1 − expected), with expected taken from 1,000 permutations of same-sized random term sets drawn, without replacement, from the full genome-wide GO Biological Process annotation (<u>org.Hs.eg.db</u> 3.22.0; 12,285 terms). Significance was assessed empirically against this null distribution and Benjamini-Hochberg-corrected across all ten pairwise comparisons.

The OCD/dogCD network communities were additionally tested for enrichment of MP Ontology terms^72^ (N = 5,446 terms; 10-2,000 genes per term) using Fisher’s exact tests. A term was considered enriched in the OCD/dogCD combined network if it reached significance (p < 5×10^-6^) in the systems-map hierarchy’s root community or any of its immediate subcommunities, and was considered associated with a given subcommunity if independently significant in that community’s own test. Terms annotated to fewer than 10 genes were excluded from the phenotype-domain summary due to unstable p-values at this sample size.

### Cross-species and brain region transcriptomic analysis

Pseudo-bulk average expression matrices were generated for human^102^ and dog (Dog Brain snRNA-seq Expression dataset, DBEx)^103^ single-cell datasets using the AverageExpression function in Seurat v5.5.1, grouping cells by annotated cell type (available as described in Data Availability). Matrices were harmonised by retaining genes with non-zero variance and cell types present in both species. Pairwise Spearman rank correlation coefficients were calculated across all shared cell types and visualised as a heatmap (ComplexHeatmap v2.26.1). To facilitate targeted biological interpretation, axes were ordered according to a predefined biological hierarchy and visually partitioned into neurons and glial cells using the row and column splitting functionalities, allowing for direct evaluation of gene network conservation within specific homologous cell populations. Differences in relative expression between neurons and glia were assessed by Wilcoxon rank-sum test.

OCD/dogCD network genes were assessed for brain cell-type specificity in the DBEx. Gene set enrichment for cell-type and brain-region specificity used the fgsea package (v1.36.2) in R (v4.5.3), ranking genes by the ratio of per-cell-type to pan-cell-type average expression (Seurat v5.5.1 AverageExpression); empirical p-values were FDR-corrected (Benjamini– Hochberg; adjusted p < 0.05).

## Use of AI tools

The authors used Claude (Anthropic) and Perplexity AI to assist with specific tasks during manuscript and code preparation, including drafting code for selected analyses, plots, and modifying PheWeb, editing manuscript text, verifying citation accuracy, and refactoring code for reproducibility in the associated GitHub repository. The core analysis pipeline was designed and implemented by the authors, who reviewed and verified all AI-assisted content and take full responsibility for the accuracy and integrity of the work.

## Data Availability

Genomic data for the 3,032 dogs sequenced at low coverage and the 11 dogs sequenced at high coverage are deposited in NCBI Sequence Read Archive (SRA) under accession numbers PRJNA675863 and PRJNA683923, respectively. Axiom canine genotype array data for 411 dogs (CanFam3.1), lifted, merged and imputed genotypes (UU_Cfam_GSD_1.0), dogCD survey item responses, phenotype, covariate and breed data, and GWAS summary statistics are available at the SciLifeLab Data Repository (Figshare), https://doi.org/10.17044/scilifelab.33339309. GWAS summary statistics for all 17 compulsive disorder traits can also be interactively browsed as Manhattan and LocusZoom plots via a PheWeb instance hosted on SciLifeLab Serve^49^.

Raw ATAC-seq, CUT&RUN and ChIP-seq reads have been deposited in the ENA under accession PRJEB124064. Processed peak files and ChromHMM state assignment bed files are available at Figshare (https://doi.org/10.6084/m9.figshare.31971747).

The canine reference genomes used here were UU_Cfam_GSD_1.0 (GCF_011100685.1) and UU_Cfam_GSD_1.0 supplemented by three Y chromosome sequences (ROS_Cfam_1.0, GCF_014441545.1). Zoonomia phastCons elements were sourced from https://doi.org/10.5281/zenodo.20136916. PacBio Iso-seq hypothalamic expression data are available under accession PRJNA657719. EpiC Dog cCRE raw data were obtained from the NCBI Gene Expression Omnibus (GEO) under accession GSE203104. As the original publication called chromatin states on CanFam3.1, we mapped their data to UU_Cfam_GSD_1.0 and re-called peaks and chromatin states using the exact same methods as in the original publication. Bed files containing the UU_Cfam_GSD_1.0-reference EpiC Dog chromatin states are also deposited in the same Figshare record described above for ChromHMM.

Dog10K variation files were sourced from the European Variation Archive (EVA) PRJEB62420 and https://kiddlabshare.med.umich.edu/dog10K/. NCBI v106 gene models were accessed from https://ftp.ncbi.nlm.nih.gov/genomes/all/annotation_releases/9615/106/. Zoonomia phyloP constraint scores were sourced from https://doi.org/10.5281/zenodo.20137136.

Orthologs were inferred using TOGA (Tool to infer Orthologs from Genome Alignments), accessed via the file <u>orthologsClassification.tsv.gz</u> from http://genome.senckenberg.de/download/TOGA/. Dog and human brain single nuclei RNA-seq datasets used in this study are available as Seurat objects here: https://doi.org/10.6084/m9.figshare.33787129. The dog brain snRNA-seq expression dataset (DBEx) can be explored via a web app here: https://dog-brain-single-cell.serve.scilifelab.se.

## Code Availability

The analysis pipeline, implemented in Nextflow with dependencies managed using pixi, is available on GitHub at https://github.com/KatarinaTe/comparativeCD-publication (version at submission: tag v1.0.0-submission; Apache 2.0 licence). It includes all code used for mapping raw sequence data, imputation, GWAS, functional annotation of GWAS regions with chromatin states, network analyses and cross-species comparisons, as well as plotting scripts. Two analyses fall outside this pipeline: SNP-based heritability was estimated using code from https://github.com/VistaSohrab/dog-gwas-heritability-nextflow, and detailed analyses and plotting of dog GWAS regions were performed using scripts available from https://github.com/JenniferMeadowsLab/Plotting_GWAS_results, (https://doi.org/10.5281/zenodo.23054418).

## Acknowledgements

We would like to thank the research participants and employees of 23andMe Research Institute for making this work possible. Computation and data handling were enabled by resources provided by the Swedish National Infrastructure for Computing (SNIC) at Uppsala Multidisciplinary Center for Advanced Computational Science (UPPMAX projects; SNIC 2022/6-229, SNIC 2021/6-208, SNIC 2025/2-420), partially funded by the Swedish Research Council through grant agreement no. 2018-05973. This work was supported by the National Bioinformatics Infrastructure Sweden (NBIS) at SciLifeLab, specifically Mahesh Binzer-Panchal assisting in the reproducibility refactoring. NBIS is a national research infrastructure funded by the Swedish Research Council (Vetenskapsrådet Dnr 2023-00153), and SciLifeLab. Illustrations of a human, dog and mouse were kindly provided by Don Meadows (Billund, Denmark). Statistical and fine-mapping advice was provided by Mats Pettersson (Uppsala University, Sweden) and Steven Gazal (University of Southern California, USA). Abraham Palmer (University of California San Diego, USA) provided computation advice on inputs for NetColoc.

## Funding Statement

NORDiC is funded by National Institute of Mental Health (NIMH) R01 MH110427 (PI J.J.C.), NIMH R01 MH105500 (PI J.J.C.) and the Swedish Research Council grants 2015– 02271 (PI D.M.-C.), 2020-01343 (PI D.M.-C.) and 2025-06864 (PI D.M.-C.). NORDiC was further supported by the Swedish Research Council (grants 2012-07111 and 2018-02487), Swedish Research Council for Health, Working Life and Welfare 2018-00221 and the Center for Innovative Medicine (CIMED). K.L.-T is a Distinguished Professor for the Swedish Research Council (2023-00391) a KAW scholar (2023.0307) and a Torsten Söderberg Academy professor in Medicine 2024. E.K.K is funded by NIH NHGRI R56HG013639 and NIH NIA U19 AG057377.

## Author Contributions

K.L.-T., K.T., J.R.S.M. and E.K.K. conceived the study. K.L.-T., K.T., J.R.S.M. and E.K.K. supervised the study. K.T., V.S., E.K.K., J.R.S.M and K.L-T performed and interpreted most of the experiments and analysed the data. K.T. and J.R.S.M developed the GWAS and NetColoc bioinformatics pipeline with assistance from R.P. M.J.C., Å.K. O.W., E.S., C.W., J.R.S.M, K.L-T and E.P. developed the dog candidate cis-regulatory resource, with formal analysis from M.J.C. V.S. performed heritability analyses and developed the dogCD PheWeb browser resource. K.A.L., B.K. and E.K.K contributed the Darwin’s Ark data collection and curation resource. G.B., C.R. and D.M.-C. contributed human OCD clinical cohort data and clinical expertise. The Nordic OCD and Related Disorders Consortium (NORDiC) and the PGC TS/OCD working group provided access to genome-wide association summary statistics prior to their formal publication. C.S. and L.L.M. contributed to GWAS fine-mapping. K.T. and E.K.K wrote the first draft of the manuscript together with J.R.S.M, V.S., M.J.C. and K.L-T. Additional comments were provided by N.I.S., M.M. and J.J.C. All authors reviewed and approved the final manuscript.

## Competing Interest Declaration

The authors declare no competing interests.

## Additional Information

Supplementary Information is available for this paper.

Correspondence and requests for materials should be addressed to Katarina Tengvall.

## Extended Data Figures

**Extended Data Fig. 1.**
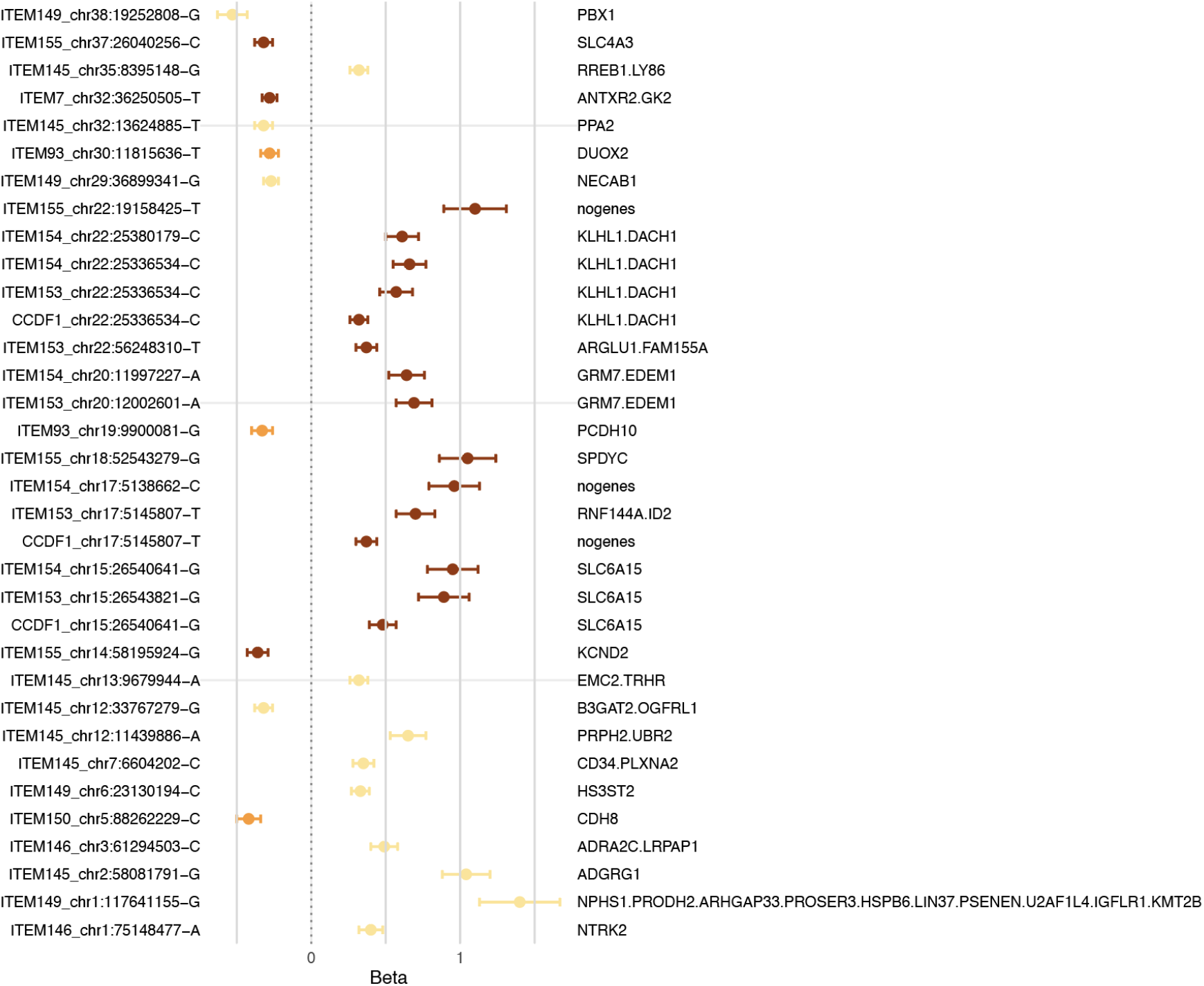
| Per statistical fine-mapped region lead SNP, effect size and implicated gene(s). For each genome-wide significant region, SuSiE statistical fine mapping was used to refine or break regions into multiple blocks, and to define the genes taken forward to NetColoc. For each SuSiE region, this plot shows the most significant SNP and its effect allele and size (Beta ± SE). Each SNP is labelled for its originating GWAS, with each effect coloured by factor grouping: Factor 1 - repetition severity (maroon), Factor 2 - trancing/pacing (orange), Factor 3 - repetition frequency (tan).

**Extended Data Fig. 2.**
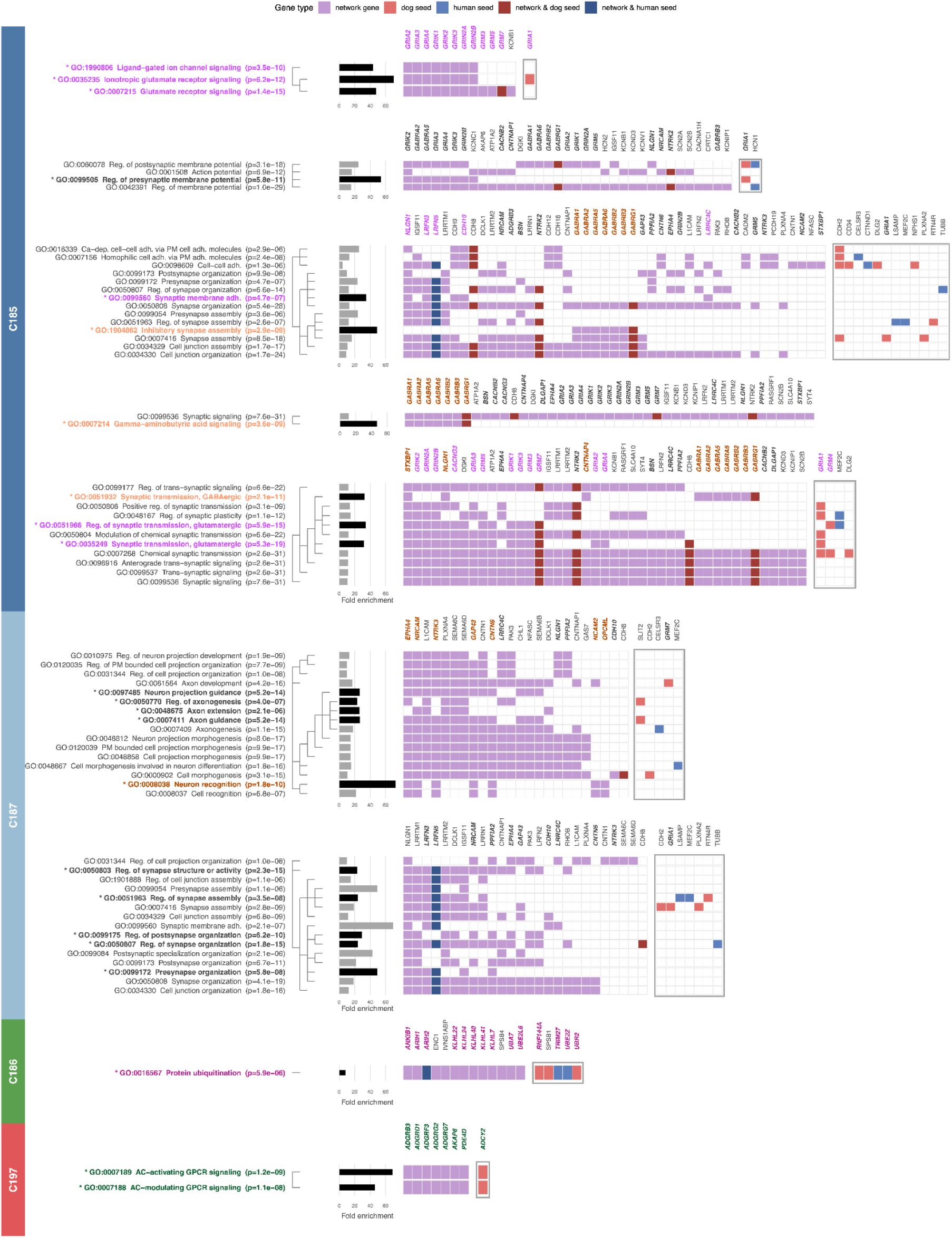
| GO Biological Process enrichment across OCD/dogCD network communities. GO Biological Process enrichment for the four gene network communities (C185, C187, C186, C197) identified by hierarchical community detection (HiDeF). Each row represents a GO term; each column a gene; colour indicates gene category: network gene (light purple), dog seed (dark red), human seed (dark blue), network & dog seed (brown), network & human seed (dark purple). Bar lengths show fold enrichment; p-values from hypergeometric tests are shown. Asterisks mark GO terms highlighted in Fig. 3b. Genes shown in bold are presented in Fig. 5. C185 is enriched for glutamatergic receptor signalling (GO:0035235, GO:0007215), GABA-A receptor signalling (GO:0007214), and synapse organisation. C187 is enriched for neuron recognition and axon guidance. C186 shows enrichment for protein ubiquitination. C197 is enriched for adenylate cyclase-activating and adenylate cyclase-modulating GPCR signalling.

**Extended Data Fig. 3.**
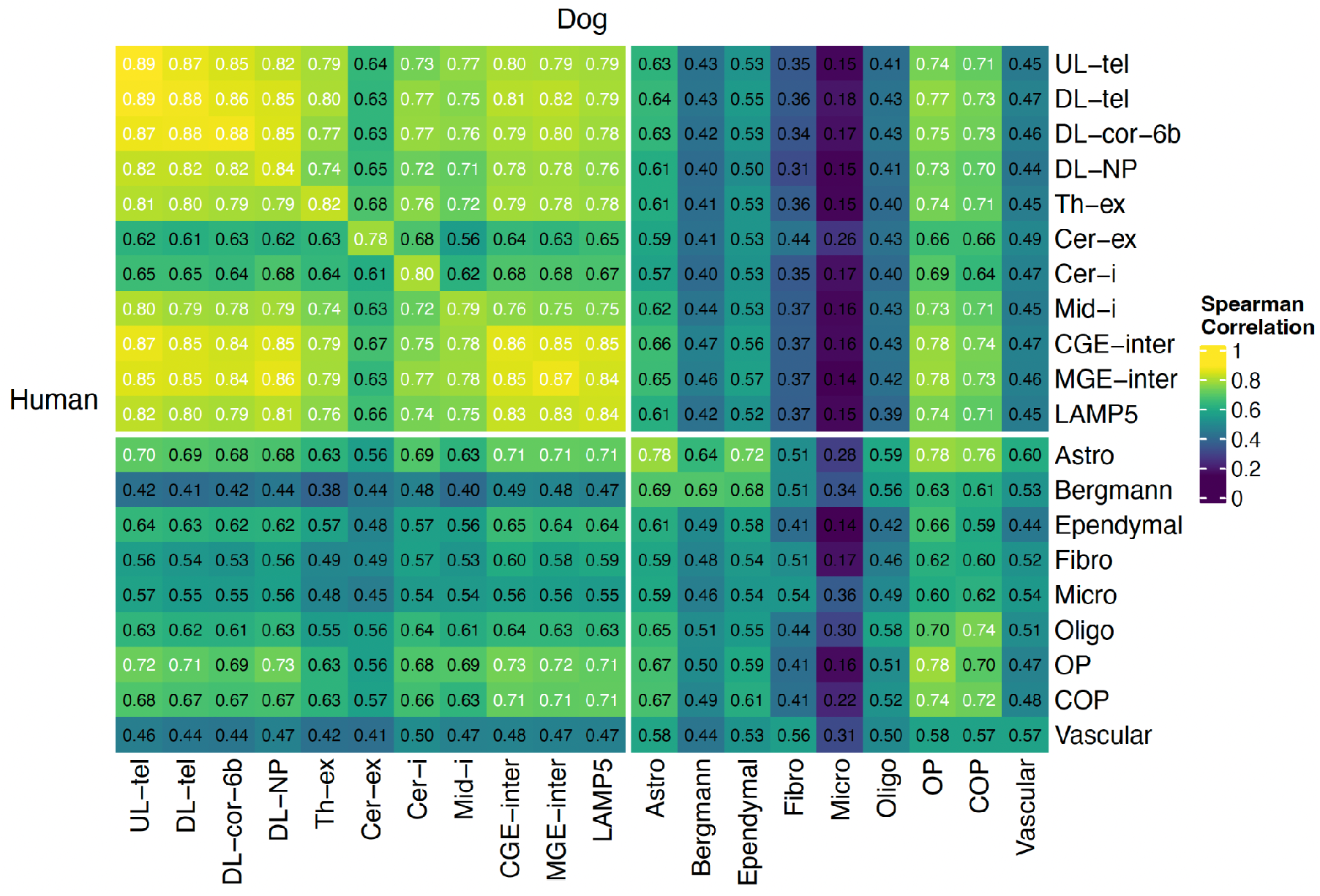
| Cross-species transcriptomic concordance of OCD/dogCD network genes. Heatmap of pairwise Spearman rank correlation coefficients between human and dog pseudo-bulk expression profiles for the 165 OCD/dogCD network genes across matched cell types. Axes are ordered by biological cell-type hierarchy and partitioned into neurons and glia. Cross-species correlations are highest in matching neuronal cell types (Spearman r = 0.78–0.89), compared with 0.36–0.78 across glial, vascular, and other non-neuronal cell types, consistent with conserved cell-type-specific transcriptional programmes across placental mammals. Neuronal cell types include: UL = upper layer; DL = deep layer; tel = intratelencephalic; cor-6b = corticothalamic and 6b; NP = near projecting; Th = thalamic; cer = cerebellum; Mid = mid-layer; CGE = caudal ganglionic eminence; MGE = medial ganglionic eminence; inter = interneurons; ex = excitatory; i = inhibitory. Glial, vascular, and other non-neuronal cell types include: Astro = astrocyte; Bergmann = Bergmann glia; COP = committed oligodendrocyte precursor; Ependymal = ependymal; Fibro = fibroblast; Micro = microglia; Oligo = oligodendrocyte; OP = oligodendrocyte precursor; Vascular = vascular. s. Of the 165 OCD/dogCD network genes, *DAOA* was not detected in this dataset’s RNA assay and is excluded from the correlations shown (n = 164 genes).

**Extended Data Fig. 4.**
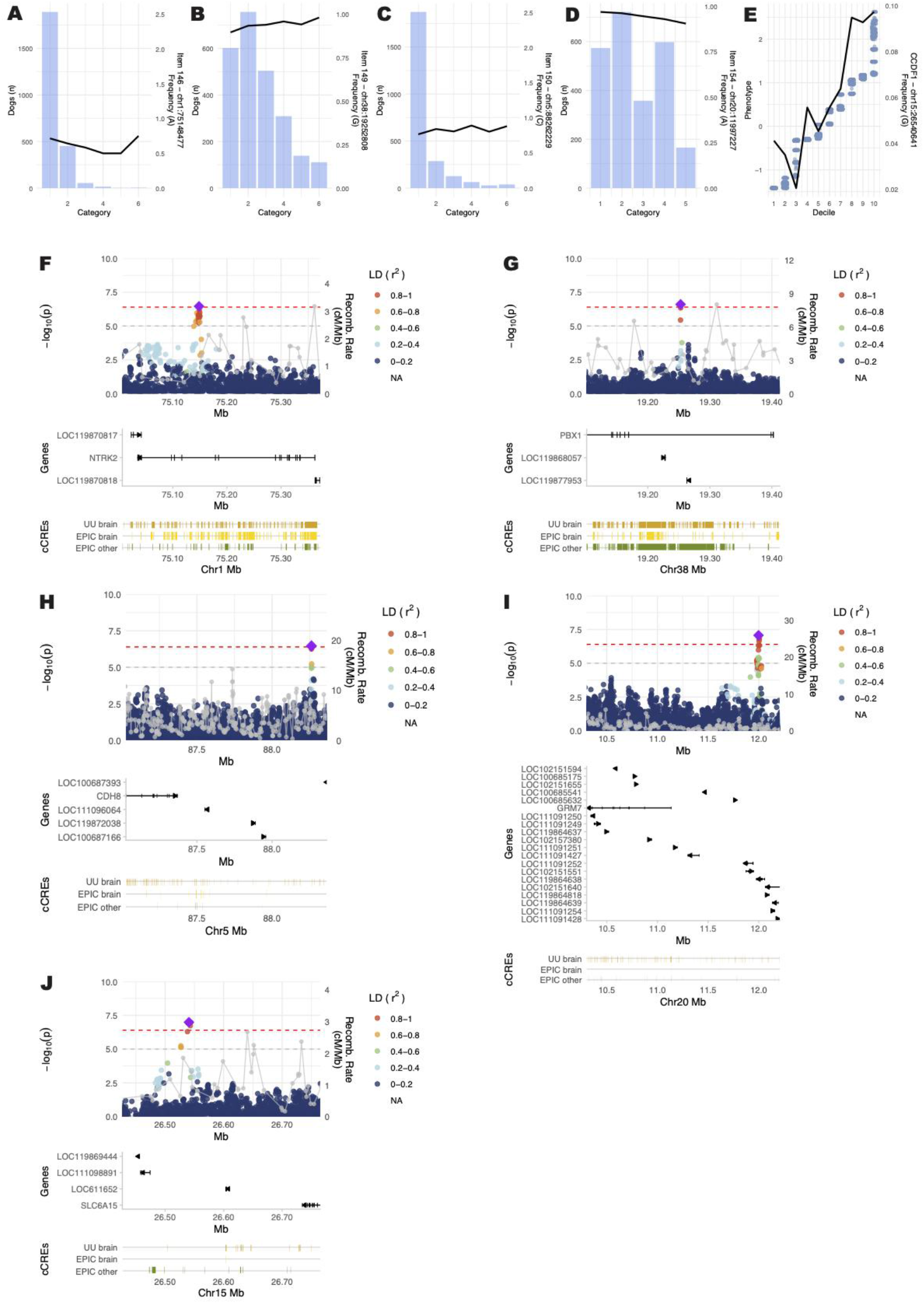
| Regional plots for a subset of dogCD GWAS hits. a–e, Distribution of the lead SNP’s effect allele across response categories of the dogCD items (a–d) or deciles of the continuous factor score (e). Bars show the number of dogs per category (left axis); the black line shows the effect allele frequency in each category or decile (right axis). f–j, Regional association plots for the same loci. Top: −log10(p) of SNPs within the region, coloured by linkage disequilibrium (r²) with the lead SNP (purple diamond); the grey line shows the recombination rate (cM/Mb), and the red and grey dashed lines mark the genome-wide (p ≤ 4×10⁻⁷) and suggestive (p ≤ 1×10⁻⁶) thresholds. Middle: NCBI v106 gene models. Bottom: cCREs from our dog brain atlas (UU brain) and from EpiC Dog brain and non-brain tissues. a,f, Q146, chr1, *NTRK2*; b,g, Q149, chr38, *PBX1*; c,h, Q150, chr5, *CDH8*; d,i, Q154, chr20, *GRM7*; e,j, Factor 1, chr15, *SLC6A15*.

## Notes

https://www.ncbi.nlm.nih.gov/bioproject/PRJNA675863

https://www.ncbi.nlm.nih.gov/bioproject/PRJNA683923

https://doi.org/10.17044/scilifelab.33339309

https://dog-cd-pheweb.serve.scilifelab.se

https://www.ebi.ac.uk/ena/browser/view/PRJEB124064

https://doi.org/10.6084/m9.figshare.31971747

https://doi.org/10.6084/m9.figshare.33787129

https://dog-brain-single-cell.serve.scilifelab.se

https://www.ncbi.nlm.nih.gov/datasets/genome/GCF_011100685.1/

https://www.ebi.ac.uk/eva/?eva-study=PRJEB62420

### Competing Interest Statement

The authors have declared no competing interest.

## Main References

1. Strom, N. I. et al. Genome-wide analyses identify 30 loci associated with obsessive-compulsive disorder. Nat. Genet. 57, 1389–1401 (2025).

2. Overall, K. L. Natural animal models of human psychiatric conditions: assessment of mechanism and validity. Prog. Neuropsychopharmacol. Biol. Psychiatry 24, 727–776 (2000).

3. Maurano, M. T. et al. Systematic localization of common disease-associated variation in regulatory DNA. Science 337, 1190–1195 (2012).

4. Jeong, Y. D. et al. Global prevalence of obsessive-compulsive and related disorders: A systematic review and modeling study. Am. J. Psychiatry appiajp20250944 (2026).

5. Fawcett, E. J., Power, H. & Fawcett, J. M. Women are at greater risk of OCD than men: A meta-analytic review of OCD prevalence worldwide: A meta-analytic review of OCD prevalence worldwide. J. Clin. Psychiatry 81, 19r13085 (2020).

6. World Health Organization. The Global Burden of Disease 2004 Update. (World Health Organization, Genève, Switzerland, 2008).

7. Pallanti, S. & Quercioli, L. Treatment-refractory obsessive-compulsive disorder: methodological issues, operational definitions and therapeutic lines. Prog. Neuropsychopharmacol. Biol. Psychiatry 30, 400–412 (2006).

8. Torbecke, J., Langhammer, T., Mewes, L., Lueken, U. & Fendel, J. C. Augmentation of cognitive-behavioural therapy for obsessive-compulsive and anxiety disorders: a protocol for a systematic review and meta-analysis. BMJ Open 14, e090431 (2024).

9. Goodman, W. K., Storch, E. A. & Sheth, S. A. Harmonizing the neurobiology and treatment of obsessive-compulsive disorder. Am. J. Psychiatry 178, 17–29 (2021).

10. Doppenberg, A. C. et al. Advancements in the understanding of the genetics of obsessive-Compulsive Disorder (OCD). Curr. Psychiatry Rep. 28, (2026).

11. Biria, M. et al. Cortical glutamate and GABA are related to compulsive behaviour in individuals with obsessive compulsive disorder and healthy controls. Nat. Commun. 14, 3324 (2023).

12. Pittenger, C., Bloch, M. H. & Williams, K. Glutamate abnormalities in obsessive compulsive disorder: neurobiology, pathophysiology, and treatment. Pharmacol. Ther. 132, 314– 332 (2011).

13. Burguière, E., Monteiro, P., Mallet, L., Feng, G. & Graybiel, A. M. Striatal circuits, habits, and implications for obsessive-compulsive disorder. Curr. Opin. Neurobiol. 30, 59–65 (2015).

14. Coelho, D. R. A. et al. Glutamatergic medications for obsessive-compulsive and related disorders: A systematic review and meta-analysis: A systematic review and meta-analysis. *JAMA Netw*. Open 8, e2452963 (2025).

15. Sullivan, P. F. et al. Psychiatric genomics: An update and an agenda. Am. J. Psychiatry 175, 15–27 (2018).

16. Trubetskoy, V. et al. Mapping genomic loci implicates genes and synaptic biology in schizophrenia. Nature 604, 502–508 (2022).

17. King, E. A., Davis, J. W. & Degner, J. F. Are drug targets with genetic support twice as likely to be approved? Revised estimates of the impact of genetic support for drug mechanisms on the probability of drug approval. PLoS Genet. 15, e1008489 (2019).

18. Hirschtritt, M. E., Bloch, M. H. & Mathews, C. A. Obsessive-compulsive disorder: Advances in diagnosis and treatment: Advances in diagnosis and treatment. JAMA 317, 1358–1367 (2017).

19. Derks, E. M., Thorp, J. G. & Gerring, Z. F. Ten challenges for clinical translation in psychiatric genetics. Nat. Genet. 54, 1457–1465 (2022).

20. Christmas, M. J. et al. Evolutionary constraint and innovation across hundreds of placental mammals. Science 380, eabn3943 (2023).

21. Villar, D. et al. Enhancer evolution across 20 mammalian species. Cell 160, 554–566 (2015).

22. Hong, H. et al. Comparative proteome and Cis-regulatory element analysis reveals specific molecular pathways conserved in dog and human brains. Mol. Cell. Proteomics 21, 100261 (2022).

23. Rosenthal, S. B. et al. Mapping the common gene networks that underlie related diseases. Nat. Protoc. 18, 1745–1759 (2023).

24. Wright, S. N. et al. Genome-wide association studies of human and rat BMI converge on synapse, epigenome, and hormone signaling networks. Cell Rep. 44, 116044 (2025).

25. Morrill, K., Chen, F. & Karlsson, E. Comparative neurogenetics of dog behavior complements efforts towards human neuropsychiatric genetics. Hum. Genet. 142, 1231–1246 (2023).

26. Overall, K. L. & Dunham, A. E. Clinical features and outcome in dogs and cats with obsessive-compulsive disorder: 126 cases (1989-2000). J. Am. Vet. Med. Assoc. 221, 1445–1452 (2002).

27. American Psychiatric Association. Diagnostic and statistical manual of mental disorders, 5th edn (American Psychiatric Publishing, 2013).

28. Chutter, M., Perry, P. & Houpt, K. Efficacy of fluoxetine for canine behavioral disorders. J. Vet. Behav. 33, 54–58 (2019).

29. Irimajiri, M. et al. Randomized, controlled clinical trial of the efficacy of fluoxetine for treatment of compulsive disorders in dogs. J. Am. Vet. Med. Assoc. 235, 705–709 (2009).

30. Garcia, A. M. et al. Phenomenology of early childhood onset obsessive compulsive disorder. J. Psychopathol. Behav. Assess. 31, 104–111 (2009).

31. Ogata, N. et al. Brain structural abnormalities in Doberman pinschers with canine compulsive disorder. Prog. Neuropsychopharmacol. Biol. Psychiatry 45, 1–6 (2013).

32. Dodman, N. H. et al. A canine chromosome 7 locus confers compulsive disorder susceptibility. Mol. Psychiatry 15, 8–10 (2010).

33. Tang, R. et al. Candidate genes and functional noncoding variants identified in a canine model of obsessive-compulsive disorder. Genome Biol. 15, R25 (2014).

34. Noh, H. J. et al. Integrating evolutionary and regulatory information with a multispecies approach implicates genes and pathways in obsessive-compulsive disorder. Nat. Commun. 8, 774 (2017).

35. Moya, P. R. et al. Rare missense neuronal cadherin gene (CDH2) variants in specific obsessive-compulsive disorder and Tourette disorder phenotypes. Eur. J. Hum. Genet. 21, 850– 854 (2013).

36. Karlsson, E. K. et al. Efficient mapping of mendelian traits in dogs through genome-wide association. Nat. Genet. 39, 1321–1328 (2007).

37. Lindblad-Toh, K. et al. Genome sequence, comparative analysis and haplotype structure of the domestic dog. Nature 438, 803–819 (2005).

38. Morrill, K. et al. Ancestry-inclusive dog genomics challenges popular breed stereotypes. Science 376, eabk0639 (2022).

39. Lord, K. A. et al. Genetic testing predicts appearance but not behavior in dogs. Proc. Natl. Acad. Sci. U. S. A. 122, e2421752122 (2025).

40. Overall, K. Manual of Clinical Behavioral Medicine for Dogs and Cats. (Saunders, 2013).

41. Salvin, H. E., McGreevy, P. D., Sachdev, P. S. & Valenzuela, M. J. The canine cognitive dysfunction rating scale (CCDR): a data-driven and ecologically relevant assessment tool. Vet. J. 188, 331–336 (2011).

42. Wright, H. F., Mills, D. S. & Pollux, P. M. J. Development and validation of a psychometric tool for assessing impulsivity in the domestic dog (Canis familiaris). Int. J. Comp. Psychol. 24, 210–225 (2011).

43. Revelle, W. Psych: Procedures for Psychological, Psychometric, and Personality Research. (2024).

44. Chalmers, R. P. Mirt: A multidimensional item response theory package for theREnvironment. J. Stat. Softw. 48, 1–29 (2012).

45. ThermoFisher Scientific. Axiom Canine Genotyping Array. https://web.archive.org/web/20251210022639/https://www.thermofisher.com/order/catalog/product/550771.

46. Meadows, J. R. S. et al. Genome sequencing of 2000 canids by the Dog10K consortium advances the understanding of demography, genome function and architecture. Genome Biol. 24, 187 (2023).

47. Yang, J., Lee, S. H., Goddard, M. E. & Visscher, P. M. GCTA: a tool for genome-wide complex trait analysis. Am. J. Hum. Genet. 88, 76–82 (2011).

48. Bi, W. et al. Efficient mixed model approach for large-scale genome-wide association studies of ordinal categorical phenotypes. Am. J. Hum. Genet. 108, 825–839 (2021).

49. Sohrab, V., Tengvall, K., Karlsson, E. K., Meadows, J. R. S. & Lindblad-Toh, K. Darwin’s Ark Dog Compulsive Disorder PheWeb. https://dog-cd-pheweb.serve.scilifelab.se (2026).

50. Gagliano Taliun, S. A., et al. Exploring and visualizing large-scale genetic associations by using PheWeb. Nat. Genet. 52, 550–552 (2020).

51. Sohrab, V. et al. Blood biomarkers and breed genetics of aging in pet dogs. bioRxivorg 2026.03.07.710297 (2026) doi:10.64898/2026.03.07.710297.

52. Wang, G., Sarkar, A., Carbonetto, P. & Stephens, M. A simple new approach to variable selection in regression, with application to genetic fine mapping. J. R. Stat. Soc. Series B Stat. Methodol. 82, 1273–1300 (2020).

53. Pettersson, M. E., Tengvall, K. & Meadows, J. R. S. Towards a standard threshold for genome wide significance in dogs. Anim. Genet. 57, e70208 (2026).

54. Wang, C. et al. A novel canine reference genome resolves genomic architecture and uncovers transcript complexity. *Commun*. Biol. 4, 185 (2021).

55. Kirilenko, B. M. et al. Integrating gene annotation with orthology inference at scale. Science 380, eabn3107 (2023).

56. Zollner, S. & Pritchard, J. K. Overcoming the winner’s curse: estimating penetrance parameters from case-control data. Am. J. Hum. Genet. 80, 605–615 (2007).

57. Palmer, C. & Pe’er, I. Statistical correction of the Winner’s Curse explains replication variability in quantitative trait genome-wide association studies. PLoS Genet. 13, e1006916 (2017).

58. Huang, J. K. et al. Systematic evaluation of molecular networks for discovery of disease genes. Cell Syst. 6, 484–495.e5 (2018).

59. Wright, S. N. et al. State of the interactomes: an evaluation of molecular networks for generating biological insights. Mol. Syst. Biol. 21, 1–29 (2025).

60. Wood, A. R. et al. Defining the role of common variation in the genomic and biological architecture of adult human height. Nat. Genet. 46, 1173–1186 (2014).

61. Kemper, K. E., Visscher, P. M. & Goddard, M. E. Genetic architecture of body size in mammals. Genome Biol. 13, 244 (2012).

62. Okada, Y. et al. Genetics of rheumatoid arthritis contributes to biology and drug discovery. Nature 506, 376–381 (2014).

63. Morris, A. P. et al. Large-scale association analysis provides insights into the genetic architecture and pathophysiology of type 2 diabetes. Nat. Genet. 44, 981–990 (2012).

64. Ha, E., Bae, S.-C. & Kim, K. Large-scale meta-analysis across East Asian and European populations updated genetic architecture and variant-driven biology of rheumatoid arthritis, identifying 11 novel susceptibility loci. Ann. Rheum. Dis. 80, 558–565 (2021).

65. Mahajan, A. et al. Fine-mapping type 2 diabetes loci to single-variant resolution using high-density imputation and islet-specific epigenome maps. Nat. Genet. 50, 1505–1513 (2018).

66. Yengo, L. et al. A saturated map of common genetic variants associated with human height. Nature 610, 704–712 (2022).

67. Zheng, F. et al. HiDeF: identifying persistent structures in multiscale ’omics data. Genome Biol. 22, 21 (2021).

68. Ashburner, M. et al. Gene ontology: tool for the unification of biology. The Gene Ontology Consortium. Nat. Genet. 25, 25–29 (2000).

69. Gene Ontology Consortium. The Gene Ontology knowledgebase in 2026. Nucleic Acids Res. 54, D1779–D1792 (2026).

70. Yu, G. et al. GOSemSim: an R package for measuring semantic similarity among GO terms and gene products. Bioinformatics 26, 976–978 (2010).

71. Anttila, V. et al. Analysis of shared heritability in common disorders of the brain. bioRxiv 048991 (2016) doi:10.1101/048991.

72. Smith, C. L. & Eppig, J. T. The mammalian phenotype ontology: enabling robust annotation and comparative analysis: The mammalian phenotype ontology. Wiley Interdiscip. Rev. Syst. Biol. Med. 1, 390–399 (2009).

73. Ahmari, S. E. & Dougherty, D. D. Dissecting ocd circuits: From animal models to targeted treatments. Depress. Anxiety 32, 550–562 (2015).

74. Shmelkov, S. V. et al. Slitrk5 deficiency impairs corticostriatal circuitry and leads to obsessive-compulsive-like behaviors in mice. Nat. Med. 16, 598–602, 1p following 602 (2010).

75. Welch, J. M. et al. Cortico-striatal synaptic defects and OCD-like behaviours in Sapap3-mutant mice. Nature 448, 894–900 (2007).

76. Dantuma, N. P. & Bott, L. C. The ubiquitin-proteasome system in neurodegenerative diseases: precipitating factor, yet part of the solution. Front. Mol. Neurosci. 7, 70 (2014).

77. Ebstein, F., Küry, S., Papendorf, J. J. & Krüger, E. Neurodevelopmental disorders (NDD) caused by genomic alterations of the ubiquitin-proteasome system (UPS): The possible contribution of immune dysregulation to disease pathogenesis. Front. Mol. Neurosci. 14, 733012 (2021).

78. Schmidt, M. F., Gan, Z. Y., Komander, D. & Dewson, G. Ubiquitin signalling in neurodegeneration: mechanisms and therapeutic opportunities. Cell Death Differ. 28, 570–590 (2021).

79. Shi, X. et al. Kelch-like proteins: Physiological functions and relationships with diseases. Pharmacol. Res. 148, 104404 (2019).

80. Lee, B. H., Meyer, C. M., Speca, D. J. & Díaz, E. Adhesion G-protein coupled receptors in neurological and psychiatric disorders. J. Psychiatr. Brain Sci. 10, (2025).

81. Unroe, K. A., Glover, M. E., Shupe, E. A., Feng, N. & Clinton, S. M. Perinatal SSRI exposure disrupts G protein-coupled receptor BAI3 in developing dentate gyrus and adult emotional behavior: Relevance to psychiatric disorders. Neuroscience 471, 32–50 (2021).

82. Zhang, X.-H. et al. Increased anxiety-like behaviors in Adgra1-/-male but not female mice are attributable to elevated neuron dendrite density, upregulated PSD95 expression, and abnormal activation of the PI3K/AKT/GSK-3β and MEK/ERK pathways. Neuroscience 503, 131–145 (2022).

83. Dodge-Kafka, K. L. et al. The protein kinase A anchoring protein mAKAP coordinates two integrated cAMP effector pathways. Nature 437, 574–578 (2005).

84. Cross-Disorder Group of the Psychiatric Genomics Consortium. Electronic address: & Cross-Disorder Group of the Psychiatric Genomics Consortium. Genomic relationships, novel loci, and pleiotropic mechanisms across eight psychiatric disorders. Cell 179, 1469–1482.e11 (2019).

85. Sen, P. et al. A bipolar disorder-associated missense variant alters adenylyl cyclase 2 activity and promotes mania-like behavior. Mol. Psychiatry 30, 97–110 (2025).

86. Giera, S. et al. The adhesion G protein-coupled receptor GPR56 is a cell-autonomous regulator of oligodendrocyte development. Nat. Commun. 6, 6121 (2015).

87. Luo, R. et al. A novel GPR56 mutation causes bilateral frontoparietal polymicrogyria. Pediatr. Neurol. 45, 49–53 (2011).

88. Bruno, K. J. & Hess, E. J. The alpha(2C)-adrenergic receptor mediates hyperactivity of coloboma mice, a model of attention deficit hyperactivity disorder. Neurobiol. Dis. 23, 679– 688 (2006).

89. Howard, D. M. et al. Genome-wide meta-analysis of depression identifies 102 independent variants and highlights the importance of the prefrontal brain regions. Nat. Neurosci. 22, 343–352 (2019).

90. Friligkou, E. et al. Gene discovery and biological insights into anxiety disorders from a large-scale multi-ancestry genome-wide association study. Nat. Genet. 56, 2036–2045 (2024).

91. Watanabe, K. et al. Genome-wide meta-analysis of insomnia prioritizes genes associated with metabolic and psychiatric pathways. Nat. Genet. 54, 1125–1132 (2022).

92. Tsai, N.-P. et al. Multiple autism-linked genes mediate synapse elimination via proteasomal degradation of a synaptic scaffold PSD-95. Cell 151, 1581–1594 (2012).

93. Hoshina, N. et al. ASD/OCD-linked protocadherin-10 regulates synapse, but not axon, development in the amygdala and contributes to fear- and anxiety-related behaviors. J. Neurosci. 42, 4250–4266 (2022).

94. Meng, X. et al. Multi-ancestry genome-wide association study of major depression aids locus discovery, fine mapping, gene prioritization and causal inference. Nat. Genet. 56, 222– 233 (2024).

95. Als, T. D. et al. Depression pathophysiology, risk prediction of recurrence and comorbid psychiatric disorders using genome-wide analyses. Nat. Med. 29, 1832–1844 (2023).

96. Purves, K. L. et al. A major role for common genetic variation in anxiety disorders. Mol. Psychiatry 25, 3292–3303 (2020).

97. Demontis, D. et al. Genome-wide analyses of ADHD identify 27 risk loci, refine the genetic architecture and implicate several cognitive domains. Nat. Genet. 55, 198–208 (2023).

98. Chen, Y. et al. Investigating the shared genetic architecture between attention-deficit/hyperactivity disorder and risk taking behavior: A large-scale genomewide cross-trait analysis. J. Affect. Disord. 356, 22–31 (2024).

99. Rao, S., Baranova, A., Yao, Y., Wang, J. & Zhang, F. Genetic relationships between attention-deficit/hyperactivity disorder, autism spectrum disorder, and intelligence. Neuropsychobiology 81, 484–496 (2022).

100. Verma, A. et al. Diversity and scale: Genetic architecture of 2068 traits in the VA Million Veteran Program. Science 385, eadj1182 (2024).

101. Milad, M. R. & Rauch, S. L. Obsessive-compulsive disorder: beyond segregated cortico-striatal pathways. Trends Cogn. Sci. 16, 43–51 (2012).

102. Siletti, K. et al. Transcriptomic diversity of cell types across the adult human brain. Science 382, eadd7046 (2023).

103. Christmas, M. J., et al. A single nuclei expression resource for exploring dog brain cell transcriptomic diversity. bioRxiv (2026) doi:10.64898/2026.09.14.751376.

104. Corrigan, E. K., et al. Conservation, alteration, and redistribution of mammalian striatal interneurons. bioRxivorg (2024) doi:10.1101/2024.07.29.605664.

105. Son, K. H. et al. Integrative mapping of the dog epigenome: Reference annotation for comparative intertissue and cross-species studies. Sci. Adv. 9, eade3399 (2023).

106. ENCODE Project Consortium et al. Expanded encyclopaedias of DNA elements in the human and mouse genomes. Nature 583, 699–710 (2020).

107. Ernst, J. & Kellis, M. ChromHMM: automating chromatin-state discovery and characterization. Nat. Methods 9, 215–216 (2012).

108. Traynelis, S. F. et al. Glutamate receptor ion channels: structure, regulation, and function. Pharmacol. Rev. 62, 405–496 (2010).

109. Niswender, C. M. & Conn, P. J. Metabotropic glutamate receptors: physiology, pharmacology, and disease. Annu. Rev. Pharmacol. Toxicol. 50, 295–322 (2010).

110. Escobar, A. P., Wendland, J. R., Chávez, A. E. & Moya, P. R. The neuronal glutamate transporter EAAT3 in obsessive-compulsive disorder. Front. Pharmacol. 10, 1362 (2019).

111. Piantadosi, S. C., Chamberlain, B. L., Glausier, J. R., Lewis, D. A. & Ahmari, S. E. Lower excitatory synaptic gene expression in orbitofrontal cortex and striatum in an initial study of subjects with obsessive compulsive disorder. Mol. Psychiatry 26, 986–998 (2021).

112. Pittenger, C. Biological mechanisms and treatment of obsessive-compulsive disorder. Annu. Rev. Clin. Psychol. 22, 455–480 (2026).

113. Brennan, B. P., Rauch, S. L., Jensen, J. E. & Pope, H. G., Jr. A critical review of magnetic resonance spectroscopy studies of obsessive-compulsive disorder. Biol. Psychiatry 73, 24–31 (2013).

114. Restifo-Bernstein, G., Guo, J., Kegeles, L. S., Shungu, D. C. & Simpson, H. B. Reconsidering the glutamate hypothesis of obsessive-compulsive disorder: A systematic review of proton magnetic resonance spectroscopy studies in unmedicated participants. J. Mood Anxiety Disord. 13, 100168 (2026).

115. Schizophrenia Working Group of the Psychiatric Genomics Consortium. Biological insights from 108 schizophrenia-associated genetic loci. Nature 511, 421–427 (2014).

116. Mäki-Marttunen, T. et al. Functional effects of schizophrenia-linked genetic variants on intrinsic single-neuron excitability: A modeling study. Biol. Psychiatry Cogn. Neurosci. Neuroimaging 1, 49–59 (2016).

117. Sohal, V. S. & Rubenstein, J. L. R. Excitation-inhibition balance as a framework for investigating mechanisms in neuropsychiatric disorders. Mol. Psychiatry 24, 1248–1257 (2019).

118. Menzies, L. et al. White matter abnormalities in patients with obsessive-compulsive disorder and their first-degree relatives. Am. J. Psychiatry 165, 1308–1315 (2008).

119. Mabb, A. M. & Ehlers, M. D. Ubiquitination in postsynaptic function and plasticity. Annu. Rev. Cell Dev. Biol. 26, 179–210 (2010).

120. Békés, M., Langley, D. R. & Crews, C. M. PROTAC targeted protein degraders: the past is prologue. Nat. Rev. Drug Discov. 21, 181–200 (2022).

121. Kennedy, C., McPhie, K. & Rittinger, K. Targeting the ubiquitin system by fragment-based drug discovery. Front. Mol. Biosci. 9, 1019636 (2022).

122. Gurney, M. E., Cogram, P., Deacon, R. M., Rex, C. & Tranfaglia, M. Multiple behavior phenotypes of the fragile-X syndrome mouse model respond to chronic inhibition of phosphodiesterase-4D (PDE4D). Sci. Rep. 7, 14653 (2017).

123. Greer, J. M. & Capecchi, M. R. Hoxb8 is required for normal grooming behavior in mice. Neuron 33, 23–34 (2002).

124. Ahmari, S. E. et al. Repeated cortico-striatal stimulation generates persistent OCD-like behavior. Science 340, 1234–1239 (2013).

125. Grotzinger, A. D. et al. Mapping the genetic landscape across 14 psychiatric disorders. Nature 649, 406–415 (2026).

126. Sarris, J. et al. N-acetyl cysteine (NAC) augmentation in the treatment of obsessive-compulsive disorder: A phase III, 20-week, double-blind, randomized, placebo-controlled trial. Prog. Neuropsychopharmacol. Biol. Psychiatry 117, 110550 (2022).

127. Biohaven Pharmaceuticals, Inc. A Study to Evaluate the Efficacy and Safety of Adjunctive Troriluzole in Obsessive-Compulsive Disorder. ClinicalTrials.gov https://clinicaltrials.gov/study/NCT04693351 (2021).

128. Maiti, R., Mishra, A., Srinivasan, A. & Mishra, B. R. Pharmacological augmentation of serotonin reuptake inhibitors in patients with obsessive-compulsive disorder: A network meta-analysis. Acta Psychiatr. Scand. 148, 19–31 (2023).

129. Rodriguez, C. I. et al. Randomized controlled crossover trial of ketamine in obsessive-compulsive disorder: proof-of-concept. Neuropsychopharmacology 38, 2475–2483 (2013).

130. Eissazade, N. et al. Efficacy and safety of 5-hydroxytryptamine-3 (5-HT3) receptor antagonists in augmentation with selective serotonin reuptake inhibitors (SSRIs) in the treatment of moderate to severe obsessive-compulsive disorder: a systematic review and meta-analysis of randomized clinical trials. Sci. Rep. 13, 20837 (2023).

131. Stern, E. R. et al. Randomized controlled trial of the effects of high-dose ondansetron on clinical symptoms and brain connectivity in obsessive-compulsive and tic disorders. Am. J. Psychiatry 182, 285–296 (2025).

132. Lee, S., Hjerling-Leffler, J., Zagha, E., Fishell, G. & Rudy, B. The largest group of superficial neocortical GABAergic interneurons expresses ionotropic serotonin receptors. J. Neurosci. 30, 16796–16808 (2010).

133. Endres, D. et al. Obsessive-compulsive disorder - A state-of-the-art review. Neurosci. Biobehav. Rev. 177, 106320 (2025).

134. Weissbrod, O. et al. Functionally informed fine-mapping and polygenic localization of complex trait heritability. Nat. Genet. 52, 1355–1363 (2020).

135. Shin, L. M. & Liberzon, I. The neurocircuitry of fear, stress, and anxiety disorders. Focus (Am. Psychiatr. Publ.) 9, 311–334 (2011).

136. Sun, T. et al. Basolateral amygdala input to the medial prefrontal cortex controls obsessive-compulsive disorder-like checking behavior. Proc. Natl. Acad. Sci. U. S. A. 116, 3799– 3804 (2019).

137. Lord, K. A., Chen, F. L. & Karlsson, E. K. An evolutionary perspective on dog behavioral genetics. Annu. Rev. Anim. Biosci. 13, 167–188 (2025).

## Methods References

138. Bartlett, M. S. Tests of significance in factor analysis. Br. J. Stat. Psychol. 3, 77–85 (1950).

139. Kaiser, H. F. A second generation little jiffy. Psychometrika 35, 401–415 (1970).

140. Cattell, R. B. The scree test for the number of factors. Multivariate Behav. Res. 1, 245–276 (1966).

141. Horn, J. L. A rationale and test for the number of factors in factor analysis. Psychometrika 30, 179–185 (1965).

142. Raiche, G. & Magis, D. nFactors: Parallel Analysis and Other Non Graphical Solutions to the Cattell Scree Test. R Package Version 2.4.1.2. https://CRAN.R-project.org/package=nFactors. (2025).

143. DePristo, M. A. et al. A framework for variation discovery and genotyping using next-generation DNA sequencing data. Nat. Genet. 43, 491–498 (2011).

144. Li, H. et al. The Sequence Alignment/Map format and SAMtools. Bioinformatics 25, 2078–2079 (2009).

145. Chang, C. C. et al. Second-generation PLINK: rising to the challenge of larger and richer datasets. Gigascience 4, 7 (2015).

146. Rubinacci, S., Ribeiro, D. M., Hofmeister, R. J. & Delaneau, O. Efficient phasing and imputation of low-coverage sequencing data using large reference panels. Nat. Genet. 53, 120–126 (2021).

147. Danecek, P. et al. Twelve years of SAMtools and BCFtools. Gigascience 10, giab008 (2021).

148. Browning, B. L., Zhou, Y. & Browning, S. R. A one-penny imputed genome from next-generation reference panels. Am. J. Hum. Genet. 103, 338–348 (2018).

149. Manichaikul, A. et al. Robust relationship inference in genome-wide association studies. Bioinformatics 26, 2867–2873 (2010).

150. Yang, J. et al. Common SNPs explain a large proportion of the heritability for human height. Nat. Genet. 42, 565–569 (2010).

151. Bi, W., Zhou, W., Dey, R., Zhao, Z. & Lee, S. GRAB: Genome-wide robust analysis for biobank data (GRAB). CRAN: Contributed Packages The R Foundation 10.32614/cran.package.grab (2025).

152. D. Turner, S. qqman: an R package for visualizing GWAS results using Q-Q and manhattan plots. J. Open Source Softw. 3, 731 (2018).

153. Thorvaldsdóttir, H., Robinson, J. T. & Mesirov, J. P. Integrative Genomics Viewer (IGV): high-performance genomics data visualization and exploration. Brief Bioinform 14, 178– 192 (2013).

154. Zou, Y., Carbonetto, P., Wang, G. & Stephens, M. Fine-mapping from summary data with the ‘Sum of Single Effects’ model. PLoS Genet 18, e1010299 (2022).

155. Kidd, J. M. Fine-scale recombination rates inferred using the canFam4 assembly are strongly correlated with previous maps of dog recombination. Mamm. Genome 37, 12 (2025).

156. Picelli, S. et al. Tn5 transposase and tagmentation procedures for massively scaled sequencing projects. Genome Res. 24, 2033–2040 (2014).

157. Lee, J., et al. Kundajelab/atac_dnase_pipelines: 0.3.0. (Zenodo, 2016). doi:10.5281/ZENODO.156534.

158. Bolger, A. M., Lohse, M. & Usadel, B. Trimmomatic: a flexible trimmer for Illumina sequence data. Bioinformatics 30, 2114–2120 (2014).

159. Gaspar, J. Improved peak-calling with MACS2. bioRxiv (2018) doi:10.1101/496521.

160. Ernst, J. & Kellis, M. Chromatin-state discovery and genome annotation with ChromHMM. Nat Protoc 12, 2478–2492 (2017).

161. Dong, M. CanFam4-referenced PhastCons scores and regions, estimated from Zoonomia 241 Mammals (v2.1) alignment. Zenodo 10.5281/ZENODO.20136916 (2022).

162. Hinrichs, A. S. et al. The UCSC Genome Browser Database: update 2006. Nucleic Acids Res. 34, D590–8 (2006).

163. Quinlan, A. R. & Hall, I. M. BEDTools: a flexible suite of utilities for comparing genomic features. Bioinformatics 26, 841–842 (2010).

164. Riemondy, K. A. et al. valr: Reproducible genome interval analysis in R. F1000Res 6, 1025 (2017).

165. Dong, M. CanFam4-referenced PhyloP scores, estimated from Zoonomia 241 Mammals (v2.1) alignment. Zenodo 10.5281/ZENODO.20137136 (2022).

166. Pardiñas, A. F. et al. Common schizophrenia alleles are enriched in mutation-intolerant genes and in regions under strong background selection. Nat. Genet. 50, 381–389 (2018).

