## Supplementary Methods 1-4 for "Dog-human gene network colocalization links synaptic signalling and ubiquitination to compulsive disorders"

### Supplementary Methods 1 | Candidate cis-regulatory element identification from dog brain

#### Commercial Tissue Sampling

Brain samples from 3 year old beagles that were sourced from a commercial provider, Zyagen, USA (**Supplementary Table 18**). Zyagen declared that all tissues were obtained from AAALAC and ISO certified facilities. Dogs were raised in compliance with SDA Animal Welfare Act (AWA) and American Kennel Club standards, and as such received regular health checks, including vaccinations and veterinary visits. Dogs were housed in spacious and proper ventilated conditions with a consistent feeding schedule and high-quality nutrition. Dogs received regular opportunities for exercise and socialization, ensuring a safe and stimulating environment with toys and enrichment activities. Dogs were euthanised by a licensed veterinarian using a sedative followed by injection of sodium pentobarbital.

#### Tissue & Extraction

Regional dissection of brain regions was performed by Zyagen (USA). Anterior cingulate cortex, cerebellum, frontal lobe, hypothalamus, occipital cortex, striatum, temporal cortex, thalamus were collected by first freezing the entire brain, then thawing and separating the tissue before storing in -80°C. Approximately 25-50 mg of tissue was cut and placed in a sterile petri dish prior to extraction or tissue disruption. Nuclei extraction was performed using 2 ml Dounce homogenizer (KIMBLE) and pestle method where pestle ‘A’ performed 10 strokes and pestle ‘B’ performed 20 strokes. The resulting solution of nuclei and nuclei extraction solution was then filtered through a 30 uM filter (Miltenyi).

#### Bulk ATAC-seq

Bulk ATAC-seq (Assay for Transposase-Accessible Chromatin with high-throughput sequencing) was performed on extracted nuclei to study chromatin accessibility and identifying open regions in the genome. Before the start of the experiment the Tn5 assembly was produced by adding 10uM Tn5ME-A primer and 10 uM Tn5MErev primer as well as 10 uM Tn5-B primer and 10 uM Tn5ME-B primer in separate tubes and run on a Thermocycler (95°C to 25°C cooling -0.1°C/second) (**Supplementary Table 19**)[1]. Subsequently, the resulting tubes were mixed with two times more of 2X DF buffer (100 mM HEPES (pH 7.2), 200 mM NaCl, 0.2 mM EDTA, 2 mM DTT, 2% Triton-X, 20% Glycerol) and 65 uM Tn5 enzyme (Picelli et al., 2014; Protein Science Facility at the Karolinska Institutet in Stockholm) for 1 hour at room temperature, which can then be placed in the -20°C freezer or used directly. First the nuclei was extracted by utilizing the Dounce extraction method with a high sugar buffer (100 mM PMSF, 14.3 mM B-mercaptoethanol, 1/2 cOmplete EDTA-free Protease Inhibitor Cocktail tablet (Merck), 1 M sucrose, 500 mM EDTA, 10% NP10) followed by filtering through the 30 uM filter (Miltenyi). The extracted nuclei was purified by centrifugation at 4°C and 900 rpm for 10 minutes and resuspended in ATAC-RSB (1 M Tris-HCl ph 7.5, 5 M NaCl, 1 M $MgCl_2$, 10% Tween). The purified nuclei were then centrifuged and resuspended with the reaction mix (2X TD buffer (1 M Tris-HCl pH 7.5, 1 M $MgCl_2$, 100% Dimethyl Formamide, pH adjusted with acetic acid to pH 7.6), 1X PBS, 1% digitonin, 10% Tween -20, 2.5 ul TN5 assembly) and placed in a Thermomixer at 37°C and 1000 rpm for 30 minutes to fragment the chromatin into smaller pieces using the transposase enzyme. The samples were immediately cleaned using the Zymo DNA Clean and Concentrator -5 kit (Zymo Research) and eluted in 20 ul of elution buffer. Afterwards adapters were added to the ends of the DNA fragments by mixing 10 uM each of primer Ad1, one of 24 primer Ad2.'s (**Supplementary Table 19**), plus NEBNext High-Fidelity 2X PCR Master Mix (New England BioLabs Inc., M0541S) and running the reaction on a Thermocycler (72°C for 5 minutes, 95°C for 5 minutes, 98°C for 30 seconds followed by 5 cycles of 98°C for 10 seconds, 63°C for 30 seconds and 72°C for 3 minutes). To test if 5 cycles were sufficient, we performed a RT-qPCR using this PCR material as input and the same Ad1/Ad2.X combination as well as 25X SYBR Green I nuclei acid gel stain (ThermoFisher Scientific) and NEBNext High-Fidelity 2X PCR Master Mix. If the amplification plateau had not been reached, additional PCR cycles were added. The final PCR product was purified with the Zymo DNA Clean and Concentrator -5 kit and quality control of the samples assessed with Qubit quantification with the dsDNA HS kit (ThermoFisher Scientific) and TapeStation (4150, Agilent) visualization with the HS D1000 ScreenTape (Agilent) (**Supplementary Table 20**).

**Histone mark identification with CUT & RUN**

We used the commercially available CUTANA ChIC/CUT & RUN Kit (EpiCypher) to map two histone modifications in the extracted tissues. The antibodies H3K4me3 (Monoclonal, MA5-11199, ThermoFisher Scientific) and H3K27me3 (Polyclonal, PA5-120811, ThermoFisher Scientific) were run on extracted nuclei. All of the experiments utilized the nuclei sample preparation method extension in the protocol with the nuclear extraction buffer (20 mM HEPES (pH 7.9), 10 mM KCl, 0.1% Triton X-100, 20% glycerol, 1 mM $MnCl_2$). Following the DNA purification and quality controls a library was prepared for each sample using the CUTANA Library Prep Kit (EpiCypher 14-1001/14-1002. Library construction was performed using 5 ng of input CUT&RUN DNA (or the total DNA available if $< 5$ ng), which underwent successive end-repair, A-tailing, and ligation of Illumina-compatible adapters. All enzymatic steps were performed under conditions optimized for CUT&RUN-derived DNA fragments, which exhibit predominantly mono- and oligo-nucleosome distributions. Antibody and library information is summarised in **Supplementary Tables 21-22**.

**Histone mark identification with ChIP-seq**

For ChIP-seq targeting H3K27ac (Acetyl-Histone H3-K27 Rabbit pAb, A7253, Abclonal) tissue samples were processed via direct SDS lysis. The IgG antibody (Normal Mouse IgG, 12-371, MilliporeSigma) was used as the control. Fresh or frozen brain tissue was scraped or finely sliced using scalpels and suspended in PBS supplemented with protease inhibitors (1× final concentration) and 10 mM sodium butyrate. After brief vortexing, cells were pelleted and the supernatant discarded. Crosslinking was performed by resuspending the pellet in PBS containing 1% formaldehyde, followed by a 10-minute incubation. Crosslinking was quenched with 2.5 M glycine and incubated for 5 minutes. Cells were pelleted and washed twice with PBS to remove residual formaldehyde. Lysis was carried out in 1% SDS lysis buffer containing protease inhibitors and sodium butyrate, with brief vortexing and mixing using a cut pipette tip. Samples were incubated on ice for 10 minutes (room temperature may be preferable to prevent SDS precipitation). Additional homogenization using a dounce homogenizer was performed if necessary. Lysates were transferred to 1.5 ml sonication tubes and sonicated using a 30-second on/off cycle for 30–45 cycles with the Bioruptor® Pico Sonication System (Diagenode AB). Post-sonication, samples were cleaned and diluted with ChIP dilution buffer (1:4 ratio) prior to immunoprecipitation. Antibody-bead complexes were prepared by incubating H3K27ac antibody with pre-washed protein A/G magnetic beads (Pierce, Thermo Scientific, 88802) for ~20 minutes at room temperature, then added to the diluted chromatin. Immunoprecipitated samples were washed sequentially with the following buffers: twice with RIPA (150 mm NaCl, 1% Nonidet P-40, 0.5% DOC, 0.1% SDS, 50 mm Tris (pH 7.4)), twice with IPWB2 (10 mM Tris-HCl (pH 7.9), 250 mM LiCl: final concentration 250 mM, 1 mM EDTA, 1% NP-40, 1% Sodium deoxycholate), twice with high salt buffer (50 mM Tris-HCl (pH 7.5), 500 mM NaCl, 0.10% SDS, 0.10% NaDOC, 1.00% Triton-X, 1 mM EDTA), twice with LiCl buffer (250 mM), and once with TE. Each buffer wash included one magnetic separation and one resuspension step. Elution was performed in IPEB buffer (100 mM $NaHCO_3$, 1% SDS) with RNase at 37°C for 15 minutes, followed by addition of 5 M NaCl and proteinase K (Qiagen AllPrep kit) for decrosslinking at 65°C for ~4 hours. DNA was purified using Ampure XP beads at a bead-to-sample ratio of at least 1.8× to retain short fragments. The purified ChIP DNA was end-repaired using the NEBNext Ultra II End Repair/dA-Tailing Module (NEB, E7546S), fragments were A-tailed with NEBNext High-Fidelity 2X PCR Master Mix and the NEBNext Multiplex Oligos for Illumina (NEB, E6440S) were ligated with Quick T4 DNA Ligase (NEB, M0202S). Antibody and quality control information provided in **Supplementary Tables 21 and 23**.

### Sequencing, mapping and peak calling

Libraries were pooled and sequenced on an Illumina NovaSeq 6000 using paired-end 2x150-bp sequencing. ATAC-seq and ChIP-seq data were processed using the ENCODE ATAC-seq v0.3.0[2] and ChIP-seq v1.9 pipelines (https://github.com/ENCODE-DCC/chip-seqpipeline2), respectively. Reads were cropped to 150 bp and reads < 100 bp removed with trimmomatic v0.41[3] prior to alignment to the canine UU_Cfam_GSD_1.0/CanFam4 reference genome[4] using BWA-MEM v0.7.17. PCR duplicates were removed using Picard MarkDuplicates, and multimapping and low-quality reads (MAPQ < 30) were excluded. Library complexity was assessed using NRF (>0.7), PBC1 (>0.5) and PBC2 (>1.0), with at least two libraries required for each brain region/histone mark combination. For ChIP-seq, NSC (>1.05) and RSC (>0.8) were used to assess signal-to-noise ratios. Peaks were called with MACS2[5] using matched input controls with the number of peaks called capped at 500 000, a p-value threshold of 0.01 and an irreproducible discovery rate (IDR) threshold of 0.05 for calling peaks. Narrow-peak mode was used for H3K4me3 and H3K27ac, and broad-peak mode for H3K27me3. For ATAC-seq, a nucleosome-free region (NFR) was required to be present. Fold-enrichment BigWig tracks were generated relative to Input. For ATAC-seq, reads were shifted by +4 bp on the plus strand and −5 bp on the minus strand. Mitochondrial reads were removed before MACS2 peak calling, without an Input control. We required peak-calling reproducibility between replicates to be ‘borderline’ or ‘pass’ for ATAC-seq and narrow-mark peaks, whereas H3K27me3 peaks were retained when present in at least two biological replicates. Peak files, signal tracks and sample-level QC metrics are available as described in Data Availability and are provided as **Supplementary Table 24**.

**Chromatin-state characterisation**

ChromHMM[6] v1.27 was used to infer states from ATAC-seq, H3K4me3, H3K27ac and H3K27me3 peak files. Peaks were binarized using BinarizeBed with a 200-bp bin size. Models containing two to 15 states were trained using LearnModel with default parameters and with the binarized bed files from all brain regions as input. Emission files from models with 2 to 14 states were then compared to the emissions file from the 15 state model using CompareModels. The nine-state model was selected as the optimum model, as it was deemed that all models with >9 states captured as much information as the 9 state model, as evidenced by the median maximum-correlation values. The likely functional element type underlying each state was determined by the combination of histone marks and open chromatin present as defined in the literature: States 1, 4, 5 and 6 were classified as promoter-like due to the presence of H3K4me3; states 2 and 8 as active enhancers due to the presence of H3K27ac (and absence of H3K4me3); state 3 as open chromatin due to only an ATAC signal present; state 9 as repressed due to the presence of H3K27me3; and state 7 as lacking detectable signal due to the absence of any peaks. Brain-region overlap was assessed using Jaccard similarity in R, and genomic coverage was calculated for each functional class and brain region as the total length occupied by the functional class divided by the genome length (UU_Cfam_GSD_1.0/CanFam4). Chromatin-state enrichment near expressed hypothalamic genes was tested using bulk gene-expression data. Here, bed files were created from an expressed genes gff to define the coordinates for all genes, exons, TSS, TES, TSS +/- 2 Kbp and TES +/- 2 Kbp. Overlap between each element and the chromatin states defined for the hypothalamus[4] were then identified using BEDtools[7] intersect. BEDtools intersect was also used to identify overlaps between chromatin states from all brain regions and Zoonomia phastCons elements[8] to assess the level of evolutionary conservation of the identified regulatory elements. Overlaps between our regulatory elements and those presented in the EpicDog resource were identified in R using ‘bed_intersect’ and ‘bed_subtract’ tools in the ‘valr’ package after liftover of EpicDog elements from CanFam3.1 to CanFam4 using the UCSC liftover tool (https://genome.ucsc.edu/cgi-bin/hgLiftOver). ChromHMM models, state assignments, are provided in **Supplementary Tables 25-26**, state files are available as described in Main Text - Data Availability.

### Supplementary Methods 2 | Phenotypic data and Exploratory Factor Analysis

#### Behavioural Questionnaire

Out of the full set of 110 Darwin's Ark behavioral questions[9], we selected 14 that cover behaviours related to compulsion in dogs (**Supplementary Table 1**). The selected behavioural survey questions were scored either on a 5-point Likert scale: (1) four questions (items 7, 93, 153 and 154) had options of strongly agree, agree, neither agree nor disagree, disagree, or strongly disagree; (2) two (items 95 and 155) had options of never, rarely, sometimes, often, or always. or on a Frequency scale: eight questions (items 145-152) had options of none, less than 30 min, 30 min - 1 hr, 1 - 2 hr, 3 - 5 hr, 5+ hr. Likert scale responses saved were codified as values {0,1,2,3,4} and frequency scale were coded as values {0,1,2,3,4,5,6}. For exploratory factor analysis (EFA) the responses were transformed to scale 1-5 or 1-6 for likert and frequency scale, respectively. Thus, for frequency scale response groups 3 - 5 hr and 5+ hr were merged resulting in a top response of 3+ hr, including proportionally few dogs with extreme repetitive behaviours. Responses for questions 7, 93, 153 and 154 were switched to make all questions comparable in terms of higher scores equals to more compulsion. In total, 12,989 dogs had responses for all 14 questions and were included in EFA.

#### Determining factors

EFA was performed on 14 survey response questions (items) using R (v4.5.3) and the R-packages nFactors (v. 2.4.1.1)[10] and base (v. 4.5.3) for evaluating the number of factors to use, and psych (v. 2.6.5)[11] for EFA. The optimal number of factors for the 14 items were estimated using nFactors::parallel and base::eigen. In the scree test and in the parallel analysis, four factors were indicated, however when performing EFA with four factors, one factor had Sum of Squares (SS)-loading below 1. With three factors, all SS-loadings were above 1. Per-item MSA values are given in **Supplementary Table 16** (overall MSA and Cronbach's alpha reported in Methods). We computed the final EFA with polychoric varimax rotation using three factors, fm="pa" (principal factor solution), max.iter=200, squared multiple correlations (SMC) set to FALSE (**Supplementary Table 1**). The three factors were: Factor 1 – items 7, 153, 154, 155; Factor 2 – items 93, 95, 150; and Factor 3 – items 145, 146, 147, 148, 149, 151, 152. For each of the three factors, the SS-loading, Proportion- and CumulativeVariance are reported in **Supplementary Table 17**.

#### Item Response Theory Factor Scoring

We calculated continuous phenotypes (factor scores) for genome-wide association analysis using Item Response Theory (IRT), which accounts for variation in question difficulty and discrimination. For each factor, items were selected based on the factor structure, and a graded response model (GRM) was fitted to the polytomous item responses using themirt (v. 1.47)[12]. Where necessary, item responses were reverse-coded prior to analysis to ensure consistent directional scoring, and all response scales were rescaled from a 0–4 to a 1–5 range. Model fit was evaluated using the M2* statistic (or C2 for Factor 2), and item-level fit was inspected via the itemfit function. Parallel solutions were examined within a confirmatory factor analysis (CFA) framework using the lavaan package (v.0.7.2)[13], allowing comparison of standardized factor loadings between the IRT and classical test theory (CTT) approaches.

Factor scores were estimated using both Expected A Posteriori (EAP) and Maximum A Posteriori (MAP) methods, with item weights derived from IRT discrimination parameters applied during scoring. MAP-based scores were retained for F1 and F3, while EAP-based scores were retained for F2, based on inspection of the score-by-standard-error distributions. The standard error of each estimate was retained for downstream assessment of measurement precision. Dogs were assigned a factor score only if responses were available for more than one item within the respective factor.

### Supplementary Methods 3 | Genotyping

#### Dog reference genomes

Two versions of the reference genome have been used through the project. The backbone (i.e., autosomes and chromosome X) is always the female UU_Cfam_GSD_1.0/CanFam4 reference genome (GCF_011100685.1)[4]. For the process of sequence alignment, and low-pass sequence imputation, we used the Dog10K UU_Cfam_GSD_1.0(CanFam4)/ROS_Cfam_1.0_chrY hybrid reference genome[14]. This reference combines UU_Cfam_GSD_1.0/CanFam4 reference genome (GCF_011100685.1)[4] with three Y chromosome contigs from Ros1.0. In this way the Y contigs could act as a read sink for the pseudoautosomal regions. Currently, no high quality, full length Y chromosome for the dog has been published.

#### Axiom array

Genotype data from 411 dogs with Axiom canine genotype array A+B variants (1,011,992 sites) were available for analysis[9]. These were positioned according to the dog reference genome, canfam3.1 (GCF_000002285.3). The LiftOver command-line program (LINUX 64-bit x86, 2022-01-31)[15] and canFam3ToCanFam4.over.chain.gz (2020-05-12) were able to successfully convert 974891 to CanFam4 positions. Quality control was performed using PLINK v1.90b7.7[16]. Filters removed variants due to --geno 0.05 (68,377 variants), --hwe midp 1e-15 (3,738 variants), non-A/C/G/T alleles --exclude variant name (37 variants), and --maf 0.01 (156,001 variants). In addition, variants that were found to lift to alternate chromosomes were removed (530 variants). The final dataset contained 746,059 variants.

#### Imputation - Low-pass sequencing

For this process, the hybrid Dog10K UU_Cfam_GSD_1.0(CanFam4)/ROS_Cfam_1.0_chrY was used. Genotype Likelihoods IMputation and PhaSing mEthod (GLIMPSE)[17] v1.1.1 was used to impute the Low-pass data. First, BCFTools v1.20[18] was used to extract polymorphic, biallelic SNP from each autosome of the phased Dog10K panel[14], (view -G -m 2 -M 2 -v snps) and then these tsv formatted files were broken into overlapping chunks for later imputation processes using supplied GLIMPSE_chunk_static script. Next, BCFTools v1.20[18] was used to calculate genotype likelihoods (bcftools mpileup -f referenceGenome -I -E -a 'FORMAT/DP' -T) from the input bams and these were used for chunked imputation using the supplied, GLIMPSE_phase_static. In the last step, the supplied GLIMPSE_ligate_static script was used to merge imputed bcf chunks, before all chromosomes were moved to vcf format and then concatenated together (BCFTools v1.20[18] convert and concat). The resultant imputed SNPs were filtered, and only those with INFO ≥ 0.8 were taken forward (SNP n = 14,274,129).

**Imputation - Axiom array**

Prior to imputation with Beagle 5.0[19], we used conform-gt.24May16.cee.jar to ensure consistency between the Axiom data and Dog10K phased reference panel[14]. Beagle 5.0 (beagle.22Jul22.46e.jar) was run using the Dog10K phased reference panel and the following settings: 30 burn-in and 40 iterations. BCFTools v1.20[18] was used to retain sites with a DR2 ≥ 0.8 resulting in a set of 14,042,437 variants.

**Downsampling for sample inclusion threshold and Genotype concordance**

All genotype concordance comparisons used GATK v4.4.0.0[20] GenotypeConcordance (with picard v3.1.1 and java/OpenJDK_17+35). First, concordance was compared between 11 sets of High-coverage and matched Low-pass samples. High-coverage samples ranged between 35.17 - 45.67X, while Low-pass samples ranged between 0.11-2.81X and non-reference concordance was calculated as >0.97 in each comparison (**Supplementary Table 3**).

To formally test the limit of Low-coverage sample depth that should be taken forward, 10 High-coverage samples were downsampled and genotype concordance between these and their original sample calculated. We used GATK v4.4.0.0 DownsampleSam with settings of P based on the estimated depth from that sample to generate 0.1X-1.0X in 10 steps. From the empirical results, we set a coverage threshold of ≥ 0.3X in order to maintain a tight distribution of non-reference discordance ≥ 0.95. This removed 85 samples.

We also compared genotype concordance between the imputed Axiom array and the imputed Low-pass sequencing data using five paired samples and a mismatched set as a negative control (shared SNPs = 2,404,919). For the matched sets, concordance and heterozygous site sensitivity were greater than 0.96, in stark contrast to the negative control values of 0.27 and 0.53 respectively (**Supplementary Table 3**). High levels of concordance with each type of genetic data allowed for dataset type merging.

**Variant dataset merging and filtering**

PLINK v1.90b7.7[16] was used for both filtering and merging the Array and Low-pass sequencing datasets. For the Low-pass dataset, filtering included --maf 0.001 (removing 854,063 variants) and --hwe midp 1e-20 (removing 61,417 variants), leaving 13,358,649 variants in 3,044 samples. The Array dataset of 411 samples was filtered as follows --hwe midp 1e-20 (removing 116 variants), --maf (0, filtered at 0.01 before imputation) leaving 14,042,321 variants. The command --bmerge merge-mode 1, merge-equal-pos was then run. For quality control, we ran plink flip-scan on the merged output and identified 7,163 variants with at least one negative LD match; these were excluded in both starting datasets and the merging process repeated. The second run identified 264 variants in flip-scan. The process was repeated again and the third time no problematic variants were identified. A final QC-filter of --geno 0.05 and --hwe midp 1e-20 removed 2,275,000 and 37,481 variants respectively, leaving 12,518,078 variants and 3,455 samples.

### Supplementary Methods 4 | Effect of batch covariate on GWAS calibration

As a quality control step prior to finalizing our GWAS pipeline, we assessed the impact of including batch as a covariate on association calibration. GWAS was performed using MLMA (GCTA v1.94.1) with a genomic relationship matrix (GRM) fitted as a random effect to account for population structure and relatedness. As a representative test, we compared results for Factor 1 between models with and without batch adjustment (age + sex vs. age + sex + batch). Genomic inflation factors were close to 1 for both models ($\lambda\_GC = 1.0003$ without batch and $\lambda\_GC = 0.9997$ with batch), a difference of only 0.0006, indicating no evidence of inflation in either analysis and no meaningful shift in calibration from including batch as a covariate. Consistent with this, per-SNP association statistics were highly concordant between the two models across 9,875,386 SNPs present in both analyses (-log10(P) Pearson's $r = 0.997$, Spearman's $\rho = 0.995$).

The largest per-SNP shifts in -log10(P) occurred exclusively among SNPs with no evidence of association. Twenty-three SNPs showed a shift in p-value of at least one order of magnitude between models (absolute change in -log10(P) $\geq 1$; range 1.00–1.88), yet none of these reached suggestive ($p < 1\times10^{-6}$) or genome-wide ($p < 4\times10^{-7}$) significance in either model, and the most significant p-value among this group was $5.18\times10^{-4}$. Restricting instead to the 11 SNPs that did reach suggestive or genome-wide significance in either model, the maximum shift was only 0.19 (mean = 0.11), confirming that batch adjustment had negligible impact on the SNPs of primary interest. Given the consistency observed for F1, batch was not included as a covariate in the GWAS models for the remaining factors or for any of the item-level GWAS runs.

### Supplementary References

167. Picelli, S. *et al.* Tn5 transposase and tagmentation procedures for massively scaled

sequencing projects. *Genome Res.* **24**, 2033–2040 (2014).

168. Lee, J. *et al. Kundajelab/atac_dnase_pipelines: 0.3.0*. (Zenodo, 2016). doi:10.5281/ZENODO.156534.

169. Bolger, A. M., Lohse, M. & Usadel, B. Trimmomatic: a flexible trimmer for Illumina sequence data. *Bioinformatics* **30**, 2114–2120 (2014).

170. Wang, C. *et al.* A novel canine reference genome resolves genomic architecture and uncovers transcript complexity. *Commun. Biol.* **4**, 185 (2021).

171. Gaspar, J. Improved peak-calling with MACS2. *bioRxiv* (2018) doi:10.1101/496521.

172. Ernst, J. & Kellis, M. Chromatin-state discovery and genome annotation with ChromHMM. *Nat. Protoc.* **12**, 2478–2492 (2017).

173. Quinlan, A. R. & Hall, I. M. BEDTools: a flexible suite of utilities for comparing genomic features. *Bioinformatics* **26**, 841–842 (2010).

174. Dong, M. CanFam4-referenced PhastCons scores and regions, estimated from Zoonomia 241 Mammals (v2.1) alignment. Zenodo https://doi.org/10.5281/ZENODO.20136916 (2022).

175. Morrill, K. *et al.* Ancestry-inclusive dog genomics challenges popular breed stereotypes. *Science* **376**, eabk0639 (2022).

176. Raîche, G., Walls, T. A., Magis, D., Riopel, M. & Blais, J.-G. Non-graphical solutions for Cattell's scree test. *Methodology (Gott.)* **9**, 23–29 (2013).

177. Revelle, W. psych: Procedures for psychological, psychometric, and personality research. *R package version* (2018).

178. Chalmers, R. P. Mirt: A multidimensional item response theory package for theREnvironment. *J. Stat. Softw.* **48**, (2012).

179. Rosseel, Y. lavaan: AnRPackage for Structural Equation Modeling. *J. Stat. Softw.* **48**, (2012).

180. Meadows, J. R. S. *et al.* Genome sequencing of 2000 canids by the Dog10K consortium advances the understanding of demography, genome function and architecture. *Genome Biol.* **24**, 187 (2023).

181. Hinrichs, A. S. *et al.* The UCSC Genome Browser Database: update 2006. *Nucleic Acids Res.* **34**, D590–8 (2006).

182. Chang, C. C. *et al.* Second-generation PLINK: rising to the challenge of larger and richer datasets. *Gigascience* **4**, 7 (2015).

183. Rubinacci, S., Ribeiro, D. M., Hofmeister, R. J. & Delaneau, O. Efficient phasing and imputation of low-coverage sequencing data using large reference panels. *Nat. Genet.* **53**, 120–126 (2021).

184. Danecek, P. *et al.* Twelve years of SAMtools and BCFtools. *Gigascience* **10**, (2021).

185. Browning, B. L., Zhou, Y. & Browning, S. R. A one-penny imputed genome from next-generation reference panels. *Am. J. Hum. Genet.* **103**, 338–348 (2018).

186. McKenna, A. *et al.* The Genome Analysis Toolkit: a MapReduce framework for analyzing next-generation DNA sequencing data. *Genome Res.* **20**, 1297–1303 (2010).
